# A nucleotide-triggered molecular switch orchestrating septin polymerization

**DOI:** 10.64898/2026.05.07.723466

**Authors:** Benjamin Grupp, Benjamin Jäckel, Andreas Günter, Jakob Rehberger, Stefan Steimle, Jan Felix Gehrke, Christian Sieg, Jan Strittmatter, Tobias Wunder, Thomas Vomhof, Johannes Ruhnke, Erik Schleicher, Stefan Gerhardt, Nils Johnsson, Thomas Gronemeyer

## Abstract

Septins are conserved cytoskeletal GTP-binding proteins that form higher-order structures critical for cytokinesis and polarized growth across opisthokonts. In budding yeast and humans, four homologous septins assemble into hetero-octameric protofilaments through alternating interactions between adjacent G-domains (G-interface) or N- and C-termini (NC-interface), which then polymerize end-to-end into filaments. How nucleotide binding and hydrolysis control filament assembly has remained elusive. We uncover a septin-specific mechanism in which nucleotide binding stabilizes the monomeric G-interface and primes it for protofilament formation. Cryo-EM and DEER spectroscopy reveal nucleotide-induced conformational changes that engage the G-interface and enable an NC-compatible conformation absent in the nucleotide-free state. *In vitro* binding and reconstitution assays confirm that G-interface formation is strictly nucleotide-dependent and required for subsequent NC-interface assembly. These findings establish unidirectional allosteric signaling from the G-to the NC-interface, revealing the molecular basis for controlled septin protofilament assembly.

## Introduction

Septins are cytoskeletal guanine nucleotide-binding proteins that assemble into higher-order structures essential for eukaryotic cell organization. Originally, septins were identified in the budding yeast *Saccharomyces cerevisiae* through temperature-sensitive mutations in the genes CDC3, CDC10, CDC11, and CDC12, which caused cytokinesis defects and established them as key regulators of the cell division cycle^1^. Subsequent sequence analysis classified septins within the P-Loop NTPase superfamily^2^ and revealed their ubiquitous presence in opisthokonts with species-specific gene expansions^3^. The septin core comprises a guanine nucleotide-binding domain (G-domain) with a modified Rossmann fold, featuring the signature G1-G4 motifs of Ras-like GTPases, plus an N-terminal extension (including helix α0) and a conserved septin unique element (SUE). The SUE comprises a characteristic β-meander (SUE-βββ) followed by helices α5 and α6, often succeeded by a C-terminal extension harboring a coiled-coil-forming region^4,5^.

Septins polymerize into apolar protofilaments through two alternating interfaces: The G-interface, mediated by two adjacent G-domains, and the NC-interface, sustained by the N- and C-terminal extensions^6^. In budding yeast, these protofilaments are linear hetero-octameric rods containing two copies of four different septin subunits, whereas in mammals, hetero-hexameric and hetero-octameric rods have been described (Supplementary Fig. 1)^7–9^. Septin protofilaments assemble into higher-order structures such as rings or gauzes, serving as scaffolds or diffusion barriers in processes including cytokinesis, vesicle trafficking, cell migration, and polarity establishment^10–16^.

Despite their critical roles in eukaryotic cell biology, septin assembly mechanisms remain poorly understood. Guanine nucleotide binding and hydrolysis are implicated in septin polymerization dynamics, yet mechanistic insight remains elusive^4,17–20^. Most septins exhibit slow GTP hydrolysis rates (0.0005-0.064 min^−1^ in yeast and humans)^18,21^, lack classical GTPase-activating proteins (GAPs) or guanine nucleotide exchange factors (GEFs)^22^, and retain bound nucleotides throughout the cell cycle^20^, suggesting a primarily structural rather than dynamic role in assembly. Mutations in septin nucleotide-binding pockets cause severe cytokinetic defects in yeast^21^ and have been linked to human pathologies, including cancer and male infertility, underscoring the physiological importance of nucleotide binding and hydrolysis^23,24^. Computational studies have proposed that nucleotide binding stabilizes the G-interface, but direct biochemical evidence for nucleotide-controlled interface formation and for allosteric coupling between G- and NC-interfaces remains scarce^25–28^.

Previous structural studies have primarily examined pre-assembled septin complexes^6,29–31^, leaving the nucleotide-dependent conformational dynamics of monomeric subunits underexplored. The only monomeric, nucleotide-free septin structure, that of *S. cerevisiae* Cdc11, reveals striking differences compared with assembled, nucleotide-bound states^31,32^. In complexes, the SUE-βββ adopts an ordered conformation that interacts with the β-meander of the adjacent subunit across the G-interface. In Cdc11_apo_, this region is partially disordered and rotated by ∼90° relative to the filament axis, potentially preventing formation of the G-interface^31,32^. The functional significance of this apo conformation and its transition to an assembly-competent state remains unresolved.

Here, integrating biochemical, structural, and *in vivo* approaches, we demonstrate that nucleotide binding triggers conformational rearrangements in the conserved SUE that are essential for G- and NC-interface formation. G-interface assembly stabilizes the nucleotide-bound state and allosterically promotes NC-interface engagement. These findings establish a sequential, nucleotide-gated pathway for septin protofilament formation, resolving the long-standing question of how nucleotides contribute to protofilament assembly.

## Results

### The G-domains of yeast septins are monomeric apo proteins

To investigate how individual septin subunits interact with guanine nucleotides without confounding polymerization effects, we first turned to *S. cerevisiae* septins, whose G-domains we previously established to be monomeric in solution^27^. The five mitotic septins (Cdc10, Cdc3, Cdc12, Cdc11, and Shs1) were expressed as G-domains lacking the N- and C-terminal extensions (NTE, CTE), which were removed to prevent noncanonical dimerization and aggregation^32–34^. These termini are unresolved in available structures (PDB 2QAG, 9BHW, 9BHT) and are routinely omitted without depleting associated nucleotides^34^. Cdc10 and Cdc12 had to be tagged with maltose-binding protein (MBP) to remain soluble. Anion-exchange chromatography of denatured proteins revealed no co-purified GTP or GDP (Fig. 1a). Incubation of purified septins with excess GTP or GDP, followed by desalting, likewise revealed no retained nucleotide (Supplementary Fig. 2a), indicating that monomeric septins are largely nucleotide-free (apo) and do not form stable nucleotide complexes.

**Fig. 1.**
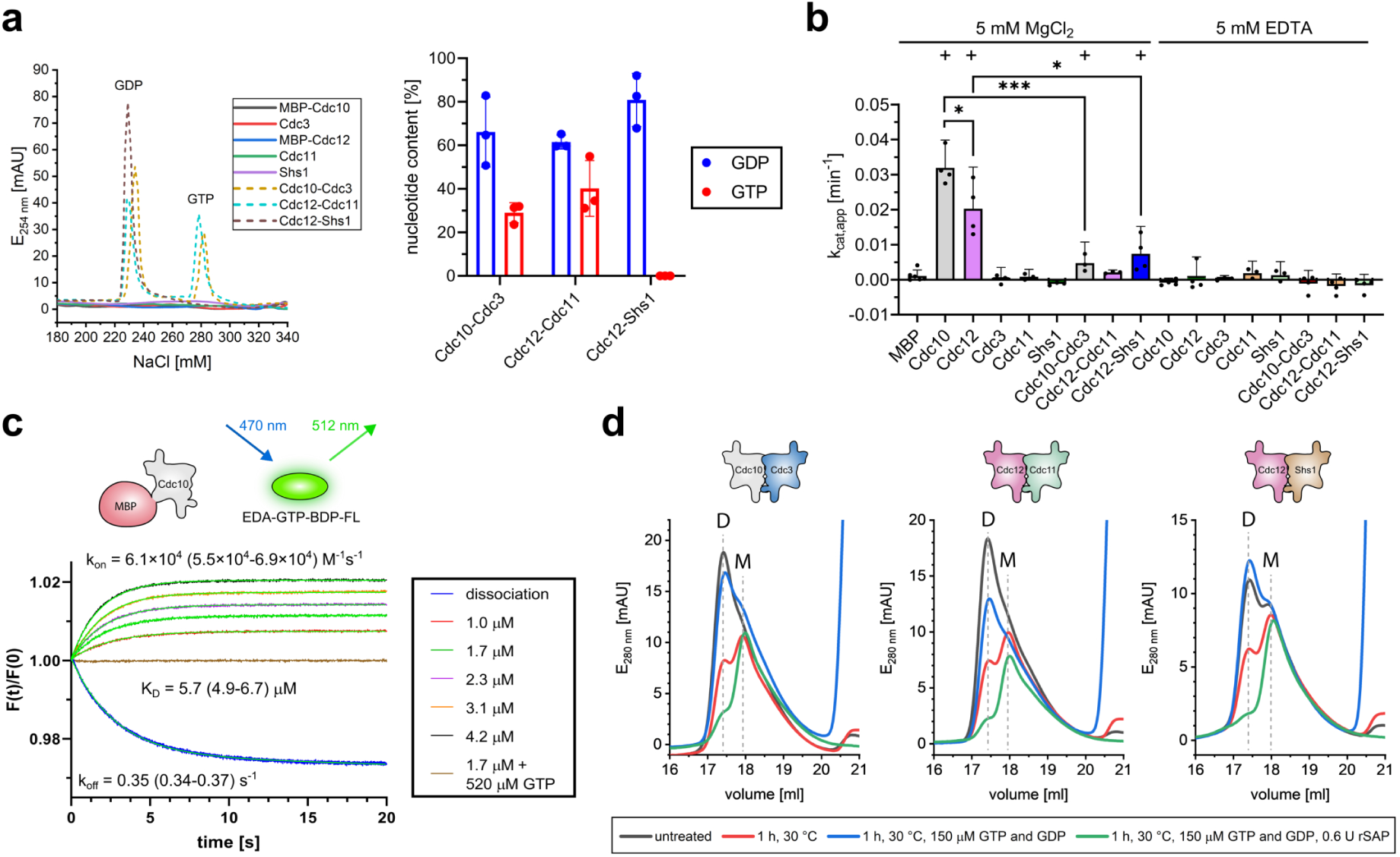
Nucleotide binding and GTPase activity of monomeric yeast septin G-domains and G-interface dimers. **a**, Nucleotide content of purified monomeric G-domains (Cdc10, Cdc3, Cdc12, Cdc11, Shs1) and co-expressed G-interface dimers (Cdc10-Cdc3, Cdc12-Cdc11, Cdc12-Shs1). Left: representative HPLC chromatograms from heat-denatured samples. Right: quantified nucleotide occupancy in dimers (arithmetic mean ± s.d., n = 3 independent experiments). Monomers are nucleotide-free; dimers are nucleotide-bound. **b**, Apparent GTP hydrolysis rates (*kcat,app*) measured by a malachite green assay at 100 µM GTP. Data are arithmetic means + symmetric 95% confidence intervals (CIs). Each data point represents the average of 2–12 technical replicates from independent experiments (Cdc10 n = 4, Cdc12 n = 4, Cdc3 n = 3, Cdc11 n = 3, Shs1 n = 3, Cdc10-Cdc3 n = 3 (MgCl2) or 4 (EDTA), Cdc12-Cdc11 n = 3 (MgCl2) or 4 (EDTA), Cdc12-Shs1 n = 4). For MBP, six replicates from the same snap-frozen protein preparation were measured on different occasions. Constructs with detectable activity are indicated (+). Significance brackets indicate results from one-way ANOVA with Tukey’s post hoc test. **c**, Representative stopped-flow fluorescence traces of EDA-GTP-BDP-FL association and dissociation with monomeric Cdc10. Global non-linear fits are shown in neon green. Kinetic parameters (*kon*, *koff*, *KD*) are geometric means with asymmetric 95% CIs from five independent experiments (association was measured at 1.0–4.2 µM Cdc10 (three preparations) or 1.0–3.1 µM (two preparations)). **d**, Investigation of dimer integrity via analytical SEC. Guanine nucleotide binding stabilizes G-interfaces. Co-expressed pairs (Cdc10-Cdc3, Cdc12-Cdc11, Cdc12-Shs1) form stable nucleotide-bound dimers. Incubation at 30 °C promotes dissociation, which is prevented by the addition of GTP and GDP. Phosphatase treatment (rSAP) amplifies dimer disruption.

Despite undetectable nucleotide retention, free phosphate detection assays revealed Mg^2+^-dependent GTP hydrolysis by Cdc10 and Cdc12. No activity was detected in the presence of EDTA and for Cdc3, Cdc11, and Shs1 (Fig. 1b). This pattern is consistent with the canonical septin hydrolysis mechanism that relies on magnesium coordination by a switch I threonine, present in Cdc10 (Thr74) and Cdc12 (Thr75) but absent in the other subunits^35^. Hydrolysis rates increased with GTP concentration in a saturable manner, approaching apparent plateau levels of 0.060 min^−1^ (Cdc10) and 0.036 min^−1^ (Cdc12) (Supplementary Fig. 2b). These catalytic rate constants are ∼60-fold (Cdc10) to ∼70-fold (Cdc12) higher than previously reported for MBP-free constructs^21^, likely reflecting improved protein stability in our preparations. Thus, although purified in an apo state and lacking detectable nucleotide retention, monomeric Cdc10 and Cdc12 retain catalytically competent active sites capable of GTP binding and turnover.

Assembled septin complexes exhibit stoichiometric amounts of retained nucleotides^20,31,33,35,36^, bind them with low-micromolar affinity^33^, and nucleotide exchange is characteristically slow, typically over hours^20,37^. By contrast, our monomeric yeast septin G-domains are nucleotide-free yet some catalytically active, implying much faster nucleotide dissociation. To quantify nucleotide binding kinetics, we performed stopped-flow fluorescence measurements with BDP-FL-labeled GTP. Cdc12 exhibited biphasic kinetics, with an initial bimolecular association (*k_1_* ≈ 4.4 × 10^4^ M^−1^ s^−1^, *k*_−_*_1_* ≈ 0.08 s^−1^) followed by a slower unimolecular step consistent with a conformational rearrangement (*k_2_* ≈ 0.023 s^−1^, *k_−2_* ≈ 0.013 s^−1^), yielding an overall *K_D_* of ∼0.7 µM (Supplementary Fig. 2c). Cdc10 displayed simpler apparent one-phase kinetics (*k_on_* ≈ 6.1 × 10^4^ M^−1^ s^−1^, *k_off_* ≈ 0.35 s^−1^, *K_D_* ≈ 5.7 µM) (Fig. 1c), whereas Cdc11 showed markedly slower association (*k_on_* ≈ 5.4 × 10^3^ M^−1^ s^−1^) and faster dissociation (*k_off_* ≈ 1.2 s^−1^), corresponding to weak binding (*K_D_* ≈ 223 µM) (Supplementary Fig. 2d). In all three cases, dissociation rate constants were orders of magnitude faster than in intact yeast septin complexes (*k_off_* ≈ 1.5 × 10^−5^ s^−1^)^20^, indicating that rapid nucleotide exchange is an intrinsic property of the monomeric state. Strikingly, no detectable binding was observed for Cdc3 or Shs1 with either BDP-FL-GTP or BDP-FL-GDP (Supplementary Fig. 2e,f), suggesting that stable nucleotide association in these subunits requires heteromeric partners. Collectively, these data reveal pronounced heterogeneity in nucleotide binding affinity and kinetics across septin subunits and support a model in which complex assembly is required for stable nucleotide binding.

### Nucleotide binding is essential for G-interface integrity in yeast septins

The rapid nucleotide exchange of monomeric septins raised the question of how assembled complexes achieve tight nucleotide binding. G-interface formation, which buries associated guanine nucleotides between two subunits, might be sufficient to provide this stability. To test this hypothesis, we purified three dimeric G-interface pairs by co-expression of the previously investigated G-domains. Nucleotide-content analysis showed that all complexes were nucleotide-associated, with average occupancies of ∼95% for Cdc10-Cdc3, ∼102% for Cdc12-Cdc11, and ∼81% for Cdc12-Shs1 (Fig. 1a). Cdc12-Shs1 contained exclusively GDP, whereas Cdc10-Cdc3 and Cdc12-Cdc11 carried mixtures of GTP and GDP, with GDP accounting for roughly 60-70% of the bound nucleotide in each case. Catalytic activities of the dimers were markedly reduced compared to the catalytically active monomers, consistent with slower nucleotide exchange upon G-interface formation (Fig. 1b).

Our previous *in silico* studies indicated that nucleotide binding enhances G-interface affinity^27,28^, suggesting that both processes act mutually stabilizing. To test this experimentally, we depleted nucleotides from purified dimers and monitored complex integrity by size-exclusion chromatography (SEC) (Fig. 1d). Upon incubation at 30 °C for 1 h, approximately 60% of the complexes dissociated into monomers, as judged from the SEC peak areas. Supplementing GTP and GDP restored the dimeric complexes. Conversely, phosphatase treatment of nucleotide-supplemented samples abolished the dimeric fraction entirely, demonstrating that nucleotide binding is a key determinant of G-interface integrity.

### Yeast septin complexes assemble in vitro from monomers in a nucleotide-dependent manner

Next, we asked whether nucleotides also govern *de novo* G-interface assembly and reconstituted G-interface pairs *in vitro* by incubating monomeric subunits with their respective G-interface partners in the presence or absence of guanine nucleotides and of either MgCl_2_ or EDTA. Formed complexes were subsequently pulled down on amylose resin and analyzed by SDS-PAGE. No pair exhibited detectable G-interface formation in the absence of nucleotide. Cdc10 and Cdc3 dimerized in the presence of MgCl_2_ with high efficiency with GTP, GDP, or a mixture of both; with reduced efficiency with GTPγS and not at all with GppNHp (Fig. 2a). EDTA abolished assembly entirely for all tested nucleotides. Cdc12-Cdc11 (Fig. 2b) and Cdc12-Shs1 (Fig. 2c) assembled with GTP, GDP, and GTPγS in both MgCl_2_ and EDTA. GppNHp failed to support Cdc12-Shs1 and yielded hardly detectable Cdc12-Cdc11 formation in EDTA.

**Fig. 2.**
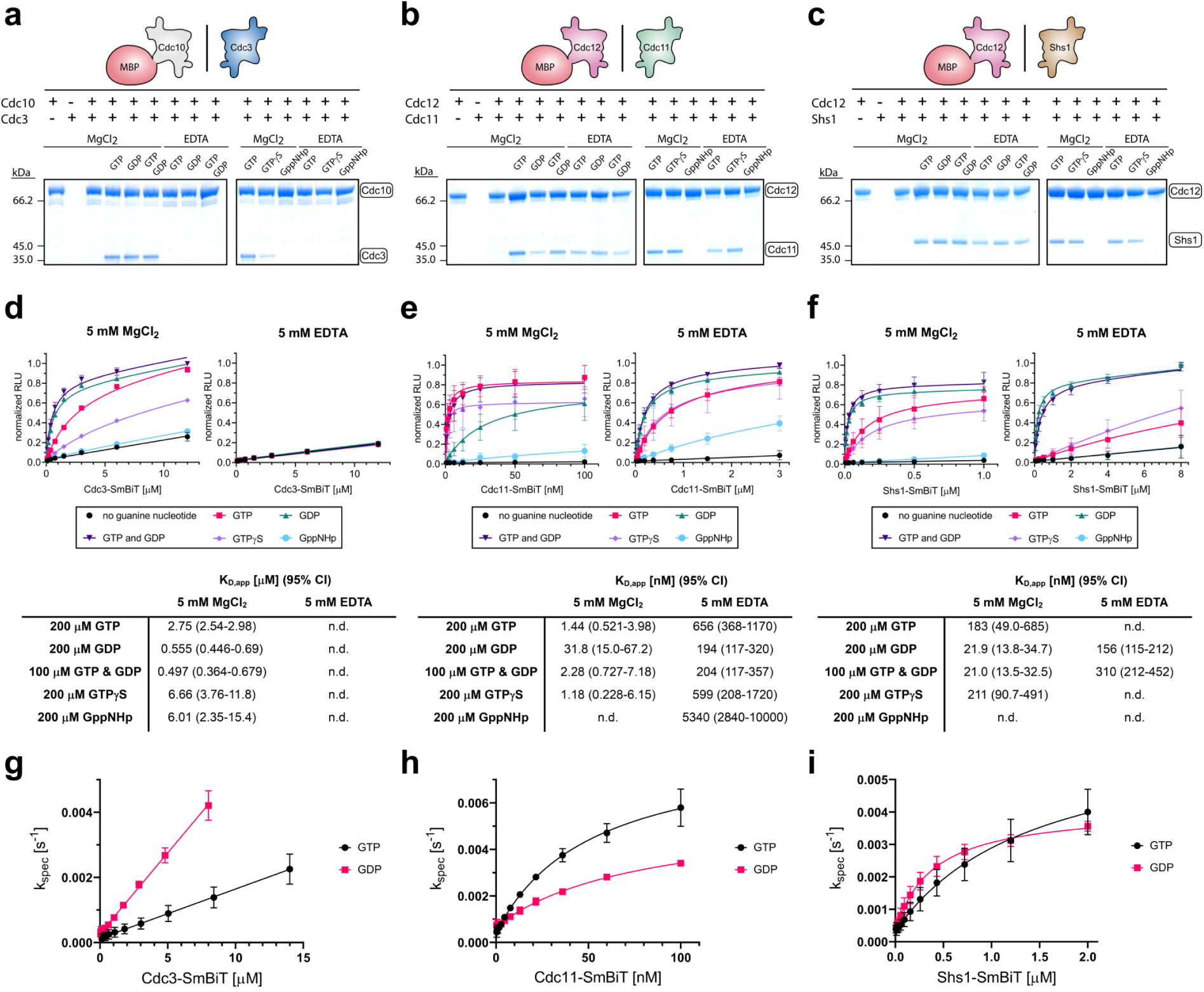
Guanine nucleotides control *de novo* assembly of yeast septin G-interface dimers. **a–c**, Reconstitution of G-interface pairs *in vitro* by co-incubation of MBP-tagged Cdc10 or Cdc12 with their respective partners (Cdc3, Cdc11, or Shs1) in the presence of the indicated guanine nucleotides and MgCl2 or EDTA, followed by pull-down on amylose resin and SDS-PAGE. **d–f**, Equilibrium titrations of Cdc10-Cdc3 (**d**), Cdc12-Cdc11 (**e**), and Cdc12-Shs1 (**f**) G-interface formation using the NanoBiT system. Curves show fits to data from n = 3 (Cdc10-Cdc3) or n = 4 (Cdc12-Cdc11, Cdc12-Shs1) independent experiments, each measured in 1–4 technical replicates. Data points are arithmetic means ± s.d. across biological replicates. Tables report geometric means for apparent *KD* values with asymmetric 95% CIs. n.d., not determined. **g–i**, Association kinetics of G-interface formation measured by the NanoBiT system in 3 mM GTP or GDP. Observed rate constants (*kspec*) from global fits are plotted against septin-SmBiT concentration and fitted by linear regression for Cdc10-Cdc3 (**g**) or a hyperbolic model for Cdc12-Cdc11 (**h**) and Cdc12-Shs1 (**i**). Data points are arithmetic means ± s.d. from n = 3 independent experiments, each measured in 2–6 technical replicates. Raw luminescence time courses and corresponding global fits are shown in Supplementary Fig. 3a–c.

A previous study on mammalian septins proposed that G-interface stability follows a high-affinity GDP/low-affinity GTP model^18^. To test whether this model applies to yeast septins, we performed equilibrium titrations using G-domain fusions to split-luciferase fragments (NanoBiT^38^). For Cdc10-Cdc3, dimerization was strictly MgCl_2_-dependent and favored by GDP or GTP/GDP (submicromolar affinities), whereas GTP and GTPγS shifted affinity into the low-micromolar range and GppNHp produced only minimal signal above background (Fig. 2d). For Cdc12-Cdc11, all nucleotides except GppNHp supported high-affinity complex formation in both MgCl_2_ and EDTA—with GTP, GTP/GDP and GTPγS near 1 nM and GDP around 30 nM in MgCl_2_ and ∼200–650 nM in EDTA—indicating substantially lower Mg^2+^-dependence than for Cdc10-Cdc3 and a GTP preference under physiological conditions (Fig. 2e). Cdc12-Shs1 behaved similarly but with a clear GDP preference: GDP and GTP/GDP gave low-(MgCl_2_) to mid-nanomolar (EDTA) affinities, GTP and GTPγS markedly weaker binding that saturated only in MgCl_2_, and GppNHp only minimal signal (^F^ig. 2f). These data indicate that yeast septin G-interfaces tolerate both GTP and GDP with complex-specific preferences and discriminate against GppNHp, challenging a general high-affinity GDP/low-affinity GTP model^18^.

While *in vivo* evidence indicates that septin protofilament assembly is rapid, with no detectable monomeric states or intermediates in wild-type cells^39–41^, our pull-down assays required hours of incubation for maximum co-precipitation, prompting us to examine G-interface association kinetics with the NanoBiT constructs. Under pseudo-first-order conditions with millimolar nucleotide, observed rate constants (*k_spec_*) for Cdc10-Cdc3 increased linearly with partner concentration, yielding second-order rate constants of ∼1.5 × 10^−4^ µM^−1^ s^−1^ (GTP) and ∼5.0 × 10^−4^ µM^−1^ s^−1^ (GDP), consistent with a simple bimolecular encounter (Fig. 2g; Supplementary Fig. 3a). By contrast, *k_spec_* for Cdc12-Cdc11 and Cdc12-Shs1 saturated with partner concentration, reaching maximal apparent association rates in the 10^−3^ s^−1^ range, and were better described by a hyperbolic model with non-zero intercepts and half-maximal rates reached at medium-nanomolar (Cdc12-Cdc11) or sub-micromolar (Cdc12-Shs1) concentrations (Fig. 2h,i; Supplementary Fig. 3b,c), indicative of an additional rate-limiting conformational step. Across all pairs, extrapolated zero-concentration rate constants fell in the same ∼10^−4^ s^−1^ range—reflecting *k_off_* for Cdc10-Cdc3 and *k_−2_* for Cdc12-containing pairs—suggesting that a slow reverse step is a conserved feature of septin G-interface formation. Together, these data indicate that slow G-interface assembly is intrinsic to isolated septin pairs, suggesting that rapid assembly *in vivo* may rely on chaperones^40,42^ or other cellular cofactors rather than the intrinsic rate of pairwise G-interface formation.

### G-interface formation enables NC-interface assembly in yeast septins

According to the current model, yeast septin protofilament assembly begins with the formation of a Cdc10-Cdc10 NC-interface dimer, while in parallel Cdc3-Cdc12 NC- and Cdc12-Shs1 G-interface dimers form^39^. Because the G-interfaces encompass the nucleotide-binding pockets, we hypothesized that the nucleotide state would primarily control G-interface stability, while the spatially distant NC-interface assembly would occur independently.

Unexpectedly, full-length Cdc10 containing the α0-helix essential for NC-interface formation^31^, eluted as a monomer in analytical SEC regardless of the presence of nucleotide, like the truncated G-domain construct (Fig. 3a). Cdc10 alone is thus incapable of NC homodimerization. The Cdc3 G-domain together with full-length Cdc10 and G-interface permissive nucleotides yielded a tetramer, whereas with Cdc10 lacking the α0-helix only dimers were formed (Fig. 3a,b; Supplementary Fig. 4e). This suggests that the tetramer corresponds to a Cdc3-Cdc10-Cdc10-Cdc3 complex in which NC-interface assembly is dependent on prior G-interface dimerization. An analogous hierarchy was observed for Cdc3 and Cdc12: no NC dimer was detectable by SEC for constructs containing α0-helices, yet G-interface dimers, tetramers, hexamers, and octamers appeared readily upon addition of G-interface partners under permissive nucleotide conditions, confirmed by SEC and mass photometry (Fig. 3c; Supplementary Fig. 4a–r; Supplementary Table 4). Mass photometry further showed that dissociation of complexes predominantly yields sub-complexes without exposed G-interfaces. Thus, we concluded that G-interface engagement allosterically stabilizes NC contacts.

**Fig. 3.**
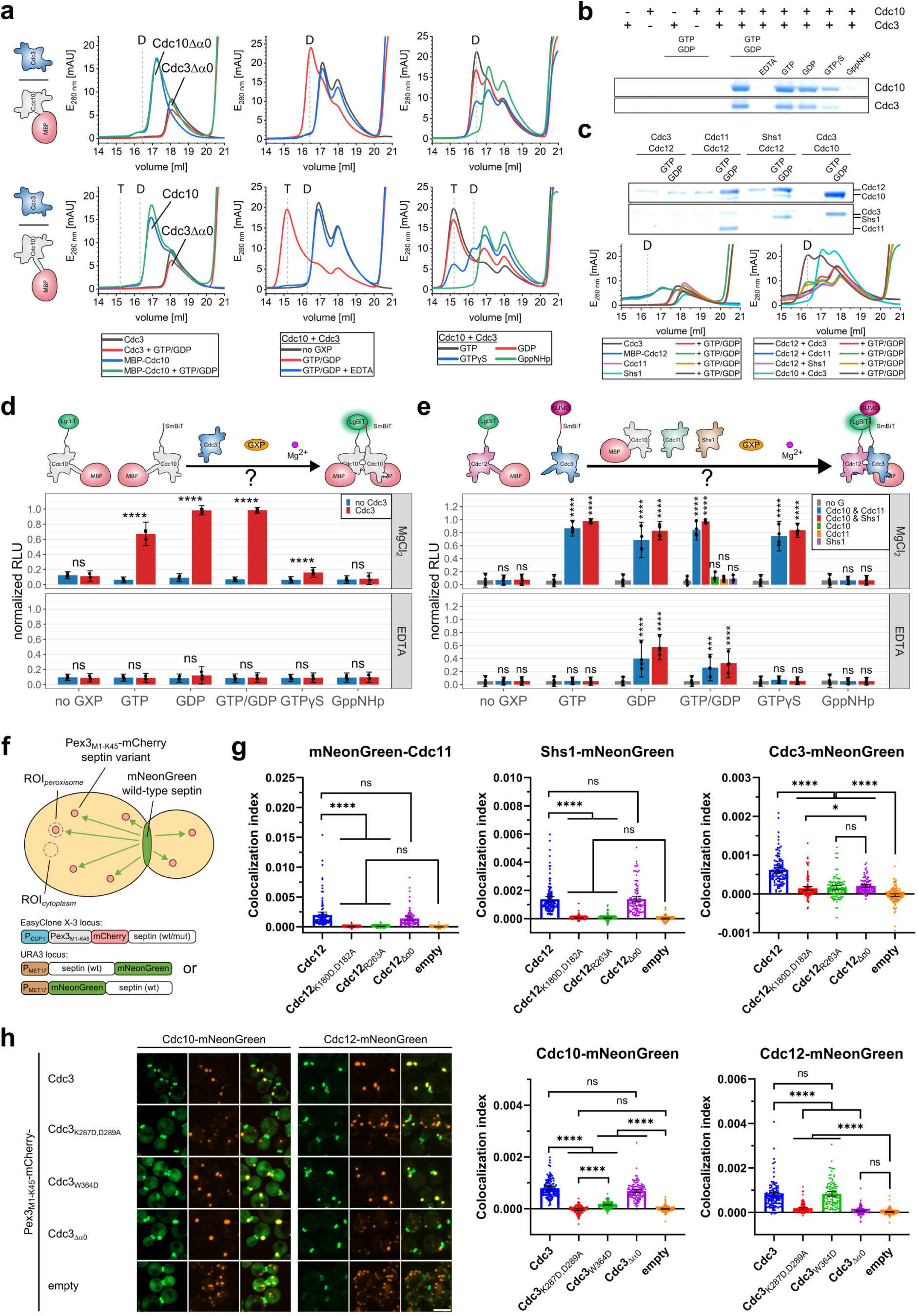
G-interface engagement is required for stable NC-interface assembly of yeast septins. **a**, Analytical SEC of G-domain (top) and full-length (bottom) Cdc10 in the absence or presence of G-domain Cdc3 and the indicated nucleotides after overnight incubation. ‘D’ and ‘T’ denote elution volumes for heterodimeric and tetrameric complexes, respectively. **b**, SDS-PAGE of tetrameric elution fractions from **a**, showing co-elution of full-length Cdc10 and G-domain Cdc3 under G-interface-permissive nucleotide conditions. **c**, Analytical SEC of α0-containing Cdc3 and Cdc12 incubated with each other or with their G-interface partners (Cdc10, Cdc11, or Shs1) in the presence or absence of an equimolar GTP/GDP mix. SDS-PAGE confirms protein identities at the heterodimeric retention volume. Cdc3 and Cdc12 do not co-elute regardless of nucleotide availability, but both co-elute with their G-interface partners in the presence of nucleotide. **d**,**e**, NanoBiT assays of NC-interface formation for Cdc10-Cdc10 (**d**) and Cdc3-Cdc12 (**e**) in the presence or absence of G-interface partners across the indicated nucleotide and Mg^2+^ conditions. Bars show arithmetic means of normalized luminescence ± symmetric 95% CIs. Individual points are averages of 2–4 technical replicates from n = 3 (Cdc10-Cdc10) or n = 4 (Cdc3-Cdc12) independent experiments. A three-way generalized least squares (GLS) model (normalized RLU ∼ G-partner × buffer × nucleotide) revealed significant three-way interactions for both Cdc10-Cdc10 (*χ*^2^(5) ≈ 932.2, *P* ≈ 2.89 × 10^−199^) and Cdc3-Cdc12 (*χ*^2^(10) ≈ 232.6, *P* ≈ 2.44 × 10^−44^) engagement. Significance annotations show post hoc simple-effects comparisons of G-partner versus no G-partner within each nucleotide-buffer combination, followed by Holm correction. **f**, Schematic of the peroxisomal membrane-recruitment assay in yeast. One septin is fused to mNeonGreen, and its binding partner is targeted to peroxisomes via Pex3M1-K45-mCherry, reporting on G- or NC-interface engagement *in vivo*. **g**, Quantification of the peroxisomal membrane-recruitment assay with Cdc12 and its mutants anchored to peroxisomes. The “empty” control consisted of a construct displaying only Pex3M1-K45-mCherry on the peroxisomal surface. Exact sample sizes are provided in Supplementary Table 3. Statistics: nonparametric Kruskal-Wallis test followed by Dunn’s multiple comparison test. **h**, Representative fluorescence images and quantification of the peroxisomal membrane-recruitment assay with Cdc3 on the peroxisomal surface, performed as in **g**. Scale bar, 5 µm.

To corroborate these findings, we leveraged the NanoBiT assay to probe the dependence of NC-interface formation on G-interface pairing across defined nucleotide and Mg^2+^ conditions (Fig. 3d,e). G-interface partners were strictly required for effective NC engagement, and relative signal intensities under different nucleotide and Mg^2+^ conditions followed the trends observed for G-interface formation (Supplementary Table 1; Supplementary Table 2). In EDTA, a significant Cdc3-Cdc12 NC signal was detected in GDP and GTP/GDP, suggesting residual Cdc10-Cdc3 G-interface formation can occur in the absence of Mg^2+^.

To determine whether this assembly hierarchy is preserved *in vivo*, we established a peroxisomal membrane-recruitment assay in yeast. One septin was fused to mNeonGreen while a second was tethered to peroxisomes via Pex3_M1-K45_-mCherry (Fig. 3f). This assay reports on the ability of interface-disrupting mutations to abolish recruitment of G- or NC-interface partners *in vivo*. We focused on the Cdc3-Cdc12 NC-interface, which showed robust mutual recruitment of wild-type proteins, and introduced lethal mutations in both interfaces, identified in a viability screen (Supplementary Fig. 5a,b). Deletion of the α0-helix largely abolished NC-mediated recruitment without affecting G-interface interactions (Fig. 3g,h; Supplementary Fig. 5c). Conversely, G-interface mutations of the G4 motif or the Arg(βb) residue (Cdc3_K287D,D289A_; Cdc12_K180D,D182A_; Cdc12_R263A_) fully eliminating G-interface recruitment also severely impaired NC engagement. Notably, the lethal Cdc3_W364D_ substitution, located in the SUE β-meander, showed significantly reduced but not entirely abolished G-interface recruitment, yet NC engagement remained unaffected, suggesting that the residual G-interface formation in this mutant remained sufficient to support NC-interface assembly.

Taken together, these data establish that G-interface formation is an essential prerequisite for NC-interface assembly, fundamentally revising the previously proposed model in which NC-interfaces initiate yeast septin protofilament assembly^39^.

### Human septins recapitulate the nucleotide-dependent assembly principles observed in yeast

To challenge the generality of our findings, we extended our analysis to human septins. Purified G-domains of SEPT9 and SEPT6 were monomeric. G-domain SEPT2 showed minor homodimerization, while G-domain SEPT7 formed G-interface homodimers that could be dissociated by high MgCl_2_ concentrations^27^. Purified monomeric subunits were nucleotide-free, while untreated SEPT7 homodimers were largely (∼70%) GDP-bound (Fig. 4a).

**Fig. 4.**
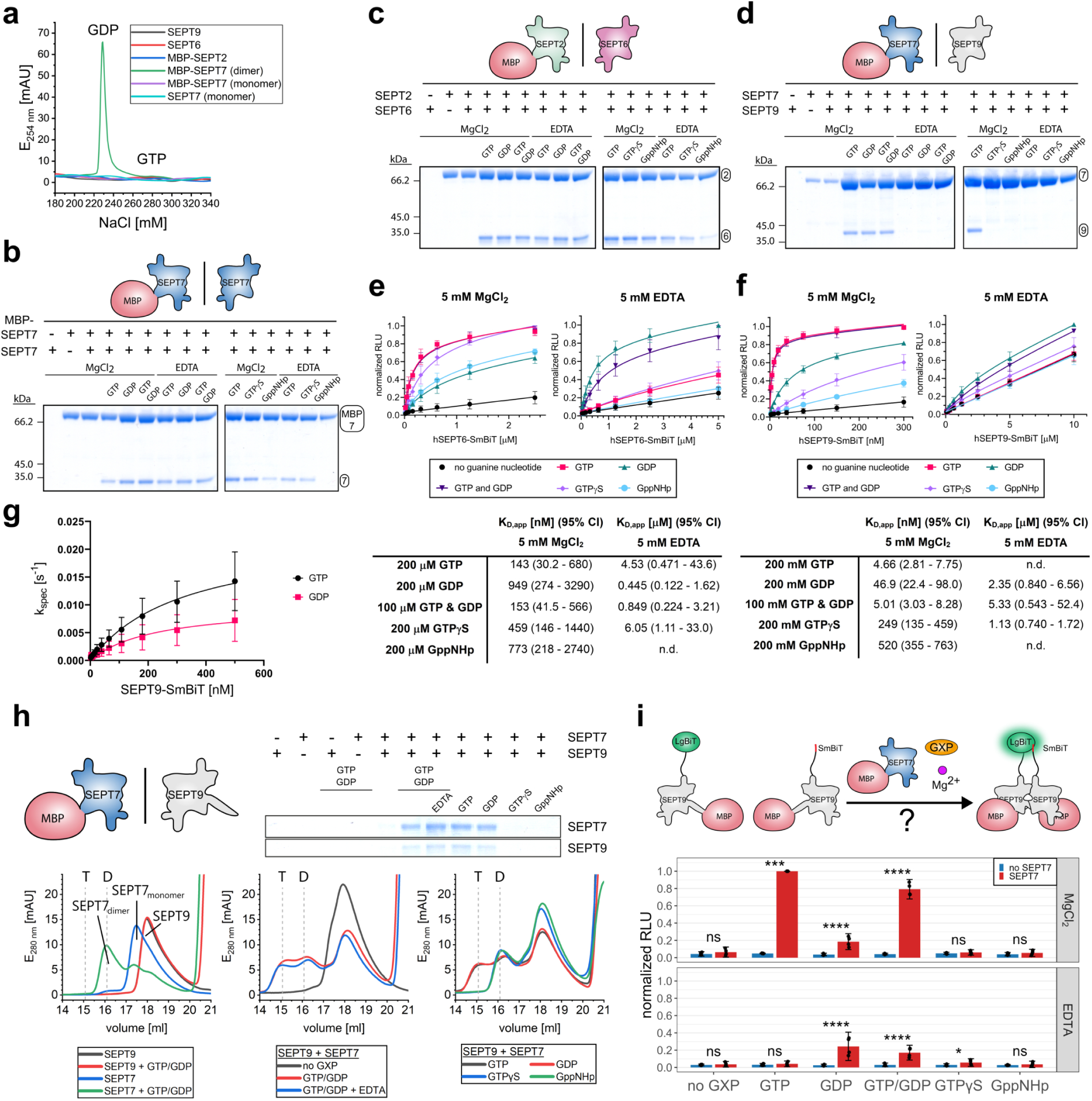
Human septins recapitulate the nucleotide-dependent assembly principles observed in yeast. **a**, Nucleotide-content analysis of purified, heat-denatured monomeric G-domains (SEPT9, SEPT6, SEPT2, SEPT7) and SEPT7 homodimers. Monomeric septins are nucleotide-free, whereas SEPT7 homodimers are predominantly (∼70%) GDP-bound. **b–d**, Reconstitution of G-interface pairs *in vitro* by co-incubation of MBP-tagged SEPT7 or SEPT2 with their binding partners in the presence of the indicated guanine nucleotides and MgCl2 or EDTA, followed by pull-down on amylose resin and SDS-PAGE. **e**,**f**, Equilibrium titrations of SEPT6-SEPT2 (**e**) and SEPT9-SEPT7 (**f**) G-interface formation using the NanoBiT system. Curves show fits to data from n = 3 independent experiments, each averaged over 1–3 technical replicates. Data points are arithmetic means ± s.d. across biological replicates. Tables report geometric means for apparent *KD* values with asymmetric 95% CIs. n.d., not determined. **g**, Association kinetics of G-interface formation measured by the NanoBiT assay in 3 mM GTP or GDP. Observed rate constants (*kspec*) from global fits are plotted against SEPT9-SmBiT concentration and fitted by a hyperbolic model. Data points are arithmetic means ± s.d. from n = 3 independent experiments, each measured in 1–7 technical replicates. Raw luminescence time courses and corresponding global fits are shown in Supplementary Fig. 3d. **h**, Analytical SEC of α0-containing SEPT9 in the presence or absence of G-domain SEPT7 and the indicated nucleotides after overnight incubation. ‘D’ and ‘T’ denote elution volumes for SEPT7 homodimers and SEPT7-SEPT9-SEPT9-SEPT7 hetero-tetramers, respectively. SDS-PAGE confirms co-elution of SEPT9 and SEPT7 in the tetrameric elution fractions under G-interface-permissive conditions. SEPT9 alone elutes as a monomer independent of nucleotide availability. **i**, NanoBiT assay of SEPT9-SEPT9 NC-interface formation in the presence or absence of G-domain SEPT7 across the indicated nucleotide and Mg^2+^ conditions. Bars show arithmetic means of normalized luminescence ± symmetric 95% CIs. Individual points represent independent experiments (n = 4) averaged over 2–3 technical replicates. The three-way GLS model (normalized RLU ∼ G-partner × buffer × nucleotide) revealed a significant three-way interaction (*χ*^2^(5) ≈ 157.5, *P* ≈ 3.36 × 10^−32^). Significance annotations show post hoc simple-effects comparisons of SEPT7 versus no SEPT7 within each nucleotide-buffer combination, followed by Holm correction.

G-interface formation was strictly nucleotide-dependent. In pull-down assays, SEPT7-SEPT7 and SEPT6-SEPT2 assembled with all tested nucleotides in both MgCl_2_ and EDTA, except GppNHp in EDTA, which abolished SEPT7-SEPT7 and reduced SEPT6-SEPT2 assembly (Fig. 4b,c). SEPT9-SEPT7 dimerization was more stringent, requiring MgCl_2_ and not being supported by non-hydrolyzable analogues (Fig. 4d). NanoBiT titrations confirmed these trends quantitatively: SEPT9-SEPT7 assembled with low-nanomolar affinity for GTP and GTP/GDP under MgCl_2_, whereas GDP and GTPγS yielded 10 to 50-fold weaker affinities. GppNHp supported only submicromolar binding, and no detectable interaction was observed in EDTA for GppNHp or GTP (Fig. 4f). SEPT6-SEPT2 displayed submicromolar affinities across all nucleotides in MgCl_2_, while EDTA shifted affinities in GTP and GTPγS to low-micromolar and abolished assembly in GppNHp (Fig. 4e). Together, human septin G-interfaces, like their yeast counterparts, are strictly nucleotide-dependent, tolerate both GTP(γS) and GDP, while showing subunit-specific differences in Mg^2+^-dependence and nucleotide selectivity.

Consistent with slow G-interface formation observed for yeast Cdc12-containing pairs, SEPT9-SEPT7 showed saturating *k_spec_* behavior in the NanoBiT assay, indicative of a rate-limiting conformational step (Fig. 4g; Supplementary Fig. 3d), with maximal apparent association rates of ∼26 × 10^−3^ s^−1^ (GTP) and ∼13 × 10^−3^ s^−1^ (GDP).

SEC and NanoBiT measurements showed that SEPT9 did not homodimerize spontaneously but required both guanine nucleotides and its G-interface partner SEPT7 to support NC-interface formation (Fig. 4h,i; Supplementary Table 6). Non-hydrolyzable GTP analogues failed to promote this interaction, and in the absence of MgCl_2_, only GDP-containing conditions supported substantial assembly. Thus, G-interface engagement and nucleotide binding jointly gate NC-interface assembly, representing a mechanism conserved from yeast to humans.

### An integrated approach of cryo-EM and DEER spectroscopy reveals major nucleotide-induced reorientations in the septin unique element

Next, we attempted to visualize nucleotide-dependent conformational changes in septins to obtain mechanistic insight into how nucleotide binding primes the G-interface. Because the only available monomeric apo structure is that of Cdc11 (PDB 5AR1), we solved the Cdc12-Cdc11 G-interface structure by single-particle cryo-EM. To increase particle size and stabilize assemblies, the Cdc11 G-domain was extended by its α0-helix to promote formation of inside-out Cdc12-Cdc11-Cdc11-Cdc12 protofilaments at low ionic strength, as described previously^43^. SEC-MALS confirmed tetramer formation in low salt (Fig. 5a). Eventually, the complex was reconstructed at 2.52 Å resolution. As observed for mammalian septin filaments^5,30^, the assembly exhibited pronounced bending around the central *C*_2_ axis (Supplementary Fig. 6a,b), resulting in lower local resolution at the NC-interface and only partial density for the Cdc11 α0-helix (Fig. 5b). Nevertheless, the map unambiguously identified GDP in Cdc12 and GTP in Cdc11 (Fig. 5c). The Cdc12-Cdc11 G-interface displays all canonical contacts, including the Arg(βb) hydrogen bond network and His(Tr1)-mediated phosphate coordination of the adjacent nucleotide (Supplementary Fig. 6c). The Cdc11-Cdc11 NC-interface is stabilized by α2/α6 C-terminal salt-bridges, while the α0-helices engage the adjacent subunit via charged interactions, consistent with low-salt assembly (Supplementary Fig. 6d).

**Fig. 5.**
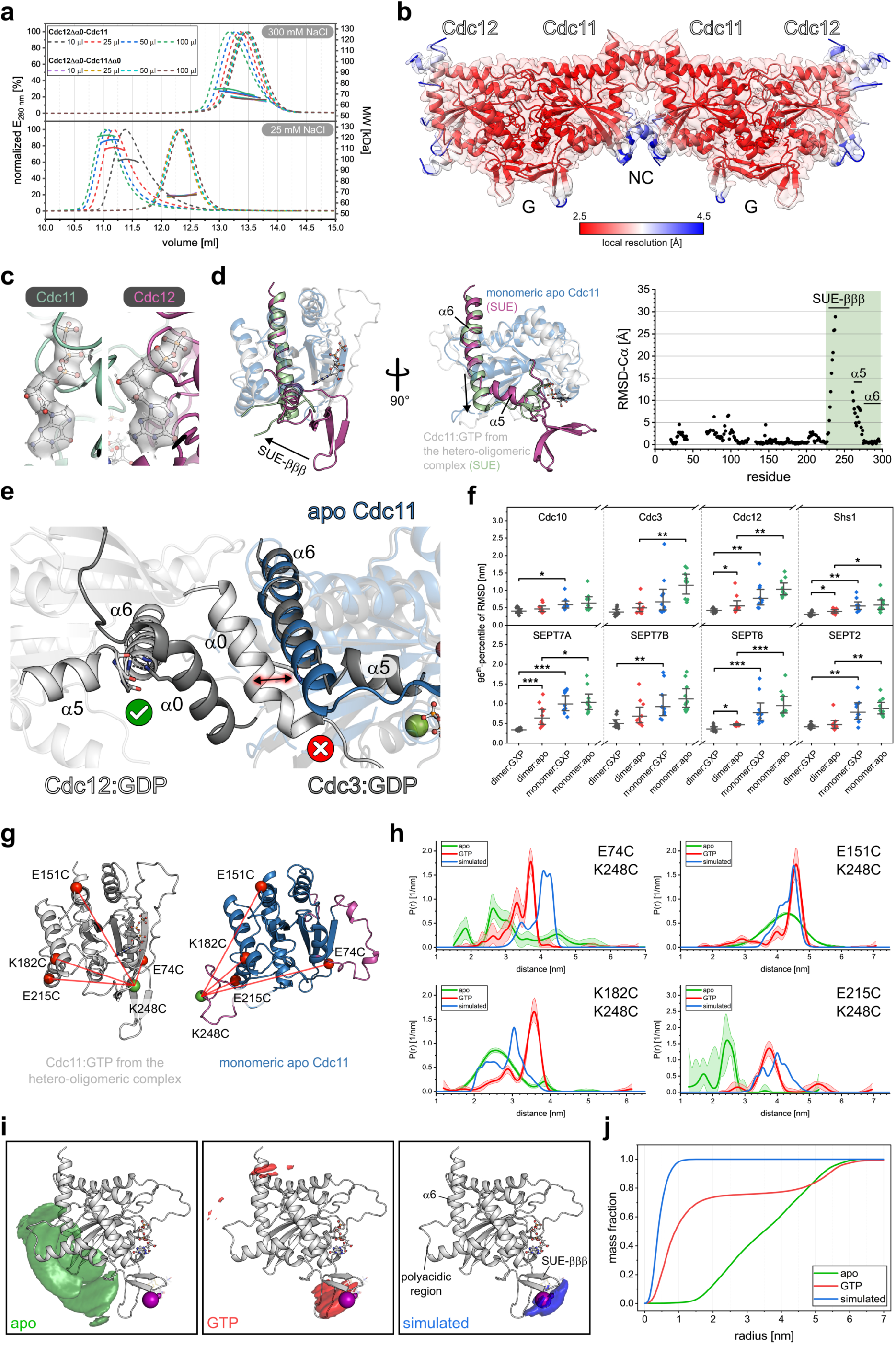
Nucleotide binding licenses the SUE-βββ for G-interface assembly. **a**, SEC-MALS analysis of Cdc12-Cdc11 G-interface dimers confirming formation of tetrameric Cdc12-Cdc11-Cdc11-Cdc12 assemblies at low ionic strength. Dashed lines indicate normalized absorbance at 280 nm and solid lines the calculated molecular weight (MW). Injected volumes were varied between 10–100 µl. Higher local concentrations and low ionic strength favor inside-out rod formation but only in the presence of the Cdc11 α0-helix. **b**, Structural model of the tetrameric complex and local-resolution cryo-EM map. **c**, Local cryo-EM maps identifying GDP in Cdc12 and GTP in Cdc11. **d**, Structural alignment of the Cdc11 cryo-EM model (G-domain) with the apo Cdc11 crystal structure (PDB 5AR1) and corresponding Cα-RMSD plot, revealing pronounced rearrangements in the SUE (light magenta (cryo-EM); pale green (apo)). **e**, Superposition of apo Cdc11 on the Cdc3-Cdc12 NC-interface (PDB 9GD4) predicts steric clash between the elongated α6 and the α0-helix of the neighboring subunit, whose conserved glutamine would otherwise stabilize the α5-α6 elbow in the polymerized state. **f**, SUE-βββ stability from all-atom MD simulations (700 ns, 10 replicas per condition) of yeast and human septins, quantified by the 95^th^-percentile of the backbone RMSD (RMSD95) of the SUE-βββ as a metric of local structural flexibility. Points show RMSD95 from individual trajectories. Horizontal bars and whiskers indicate geometric means and asymmetric 95% CIs. Brackets denote Wilcoxon rank-sum contrasts with Holm correction. The dimer:GXP - monomer:apo contrast was significant in all instances and is omitted for clarity. **g**, Labeling sites for DEER spectroscopy in the apo (blue) and the G-interface-engaged, GTP-bound state of Cdc11 (white). Spheres indicate Cα positions of labeled residues; green marks K248C, and red marks residues that were individually combined with K248C in DEER pairs ([E74C,K248C], [E151C,K248C], [K182C,K248C], [E215C,K248C]). Unresolved regions (including the SUE-βββ) in the apo structure were modeled in SWISS-MODEL^97^ (light magenta). **h**, DEER distance probability distributions of Cdc11 variants spin-labeled at position 248 and four reference positions, shown for monomeric apo and GTP-bound states alongside theoretical distributions from MMM rotamer simulations based on the cryo-EM model (simulated). The distributions *P(r)* are normalized to integrate to 1. Shaded bands around the distributions of [E74C,K248C], [E151C,K248C], and [K182C,K248C] represent 95% confidence bands derived from the ComparativeDeerAnalyzer 2.0 (CDA2.0)^91^ calibration, whereas shaded bands around the distributions of [E215C, K248C] show validation-derived uncertainty bands obtained by varying the background-fit start time as reported by DeerAnalysis2022^92^. **i**, Multilateration-derived probability density maps, shown as isodensity surfaces overlaid on the Cdc11 cryo-EM structure and contoured at 50% of total probability mass. Spheres indicate rotamer-derived N-O midpoint positions whose sizes are scaled by rotamer probability as calculated by MMM. **j**, Cumulative mass-fraction curves derived from integrating the probability density maps for apo, GTP-bound, and simulated ensembles around the N-O midpoint positions.

Comparison of polymerized Cdc11 with the apo crystal structure revealed striking rearrangements within the SUE, spanning the β-meander (SUE-βββ) through helices α5 and α6 into the NC-interface (Fig. 5d). In the filament, residues D263-S274 form a canonical α5-helix followed by a short elbow connecting to α6, creating a pocket in the NC-interface for α0 of the adjacent subunit. In apo Cdc11, the α5-helix is absent, residues D263-L267 are unfolded, and the α6-helix extends N-terminally, eliminating the elbow. The SUE-βββ itself is rotated ∼90° away from the filament axis toward α5′ and only partially resolved, indicative of high flexibility. Superposition of apo Cdc11 onto the Cdc3-Cdc12 NC-interface (PDB 9GD4) predicts steric clash between this elongated α6 and the α0-helix of the neighboring subunit (Fig. 5e). Given that SUE-βββ residues are critical for G-interface stabilization^6,27^ and that α0 stabilizes the NC-interface^31,43,44^, these data suggest that apo Cdc11 adopts an oligomerization-incompetent conformation, with the SUE acting as a nucleotide-state sensor.

To probe the dynamics underlying these conformational states, we performed MD simulations of yeast and human septins in monomeric and G-interface-dimeric states, in apo and nucleotide-bound forms. Quantifying the flexibility of the SUE-βββ over the trajectory, a consistent stability hierarchy was observed across all systems: dimer:GXP > dimer:apo > monomer:GXP > monomer:apo (with GXP indicating bound nucleotide). G-interface engagement emerged as the dominant suppressor of SUE-βββ flexibility, and nucleotide binding exerted a weaker but detectable additional stabilizing effect (Fig. 5f; Supplementary Table 7). Although the 700 ns timescale of the simulations did not capture the full α5 rearrangement, complete Cdc11-like β-meander unfolding was observed in several monomer simulations (Supplementary Fig. 7a,b; Supplementary Movie 1; Supplementary Movie 2). Consistent with our biochemical findings, G-interface formation suppressed nucleotide dynamics, whereas monomeric trajectories exhibited significantly increased instability in five of the eight subunits, culminating in spontaneous GDP ejection in two simulations (Supplementary Fig. 7c; Supplementary Movie 3; Supplementary Movie 4). Together, these results highlight the intrinsic fragility of both the SUE-βββ and nucleotide retention in the absence of stabilizing G-interface contacts.

We performed double electron-electron resonance (DEER) spectroscopy to experimentally validate the *in silico* findings. Engineered Cdc11 variants carrying a spin label in the SUE-βββ (K248) and at structurally invariant reference sites were employed. All constructs retained nucleotide-dependent Cdc12 binding, displayed native-like secondary structure indicated by CD spectroscopy, and remained monomeric after labeling (Supplementary Fig. 8a–c). In the DEER measurements, nucleotide-free samples were compared with samples containing 5 mM GTP, sufficient for >90% occupancy as estimated from stopped-flow and ITC-derived binding constants (Supplementary Fig. 2d; Supplementary Fig. 8d). DEER distance distributions showed clear nucleotide-dependent shifts across all label pairs (Fig. 5h), which we converted into 3D probability density maps of the spin-label ensemble (Fig. 5i). Compared to a simulated ensemble derived from the static cryo-EM model, which localized >98% of probability mass within 1 nm of the expected rotamer positions, the GTP-bound state showed ∼55% within this radius, with a secondary population displaced by ∼5 nm toward the polyacidic/α6 region, indicating a dynamic equilibrium between interface-competent and incompetent conformations (Fig. 5j). In the apo state, probability mass was almost entirely displaced from the expected position, with less than 1% within 1 nm, consistent with a predominantly flexible, interface-incompetent conformation. Thus, nucleotide binding is sufficient to trigger SUE-βββ rearrangements into a metastable, filament-like conformation inaccessible in the apo state.

Together, the cryo-EM, MD, and DEER analyses support a model in which nucleotide binding drives large-scale SUE-βββ reorientation to prime the G-interface for assembly, and subsequent G-interface formation locks this conformation to permit stable NC-interface engagement.

## Discussion

Septin filament assembly has long been recognized as nucleotide-dependent, yet how guanine nucleotides enforce the ordered engagement of septin interfaces has remained unresolved^19,21,33,45^. Here, we show that nucleotide binding triggers a conformational switch within the SUE that primes the G-interface for assembly and allosterically enables subsequent NC-interface formation. Using complementary biochemical, structural, and cellular approaches across septins from yeast and humans, we demonstrate that G-interface formation is strictly nucleotide-dependent and functions as an obligate gatekeeper for NC-interface engagement. The predominantly unidirectional G-to-NC allosteric communication represents a molecular switch that enforces ordered septin protofilament assembly and revises the current model of yeast septin protofilament formation^39^.

An important observation underlying our proposed mechanism is that isolated, monomeric septin G-domains purify in the apo state and bind guanine nucleotides only transiently, with micromolar affinity, as shown by stopped-flow and ITC measurements. Formation of the G-interface fundamentally shifts this equilibrium. Co-expressed G-interface dimers purify with near-stoichiometric nucleotide occupancy, dissociate into monomers upon nucleotide removal, and display markedly reduced GTPase activity, consistent with the very low nucleotide exchange and hydrolysis rates reported for septin complexes across species^18,20,33,36,37,46^. Thus, nucleotide binding and G-interface formation are reciprocally stabilizing, with each process reinforcing the other. The absence of detectable nucleotide binding by isolated Cdc3 and Shs1 in stopped-flow experiments—despite both subunits being nucleotide-bound in complexes (PDB 8SGD, 8FWP, 8PFH, 9GD4)—suggests that efficient nucleotide association by these subunits requires coincident engagement with nucleotide-loaded partners, reinforcing G-interface dimers as the obligate building blocks of septin assemblies.

Quantitative split-luciferase equilibrium titrations reveal that G-interfaces readily tolerate both GTP and GDP as assembly-promoting ligands but mainly discriminate against the non-hydrolyzable analogue GppNHp. The inconsistent support of G-interface formation by GppNHp, in contrast to the partial activity retained with GTPγS, points to a strict dependence on precise γ-phosphate chemistry in the GTP-bound state rather than on GTP hydrolysis—consistent with previous findings^33^. Together, these observations suggest that nucleotide binding functions primarily as a licensing step activating the G-interface rather than as a classical hydrolysis-driven switch.

Furthermore, our findings suggest that septin G-interfaces do not conform to a universal high-affinity GDP/low-affinity GTP paradigm as previously suggested^18^. We observe substantial interface-specific heterogeneity across yeast and human septins, with some interfaces favoring GTP, others GDP, and still others showing minimal discrimination. This diversity is consistent with the mixed nucleotide content reported for purified septin complexes across species^20,30,33,47,48^. In yeast, such interface-specific nucleotide preferences provide a mechanistic basis for how intracellular fluctuations in the GTP:GDP-ratio may influence septin filament composition *in vivo*^49,50^. Notably, the Cdc12-Shs1 G-interface is preferentially stabilized under GDP-rich conditions, whereas the Cdc12-Cdc11 interface exhibits higher affinity in GTP-rich environments. Consistent with this mechanism, experimentally lowering the cellular GTP:GDP-ratio reduces Cdc11 incorporation while favoring Shs1 recruitment into septin rings^39^. These findings suggest that differential nucleotide preferences at the Cdc12 G-interface act as a selective filter that translates global nucleotide availability into alternative protofilament compositions.

Our data also resolve the long-standing question of whether and how G- and NC-interfaces communicate allosterically^5,25,26,35,51^. Contrary to the current model proposing NC-driven nucleation^39^, our peroxisomal colocalization experiments indicate that full-length septins bearing intact NC-interface elements are not competent for effective NC engagement unless stable G-interface formation, primed by guanine nucleotide binding, has occurred. Disruption of nucleotide binding or essential G-interface contacts abolishes NC-mediated recruitment, whereas selective disruption of NC-interface elements leaves G-interface engagement intact, indicating that allosteric control is exerted unidirectionally from the G-to the NC-interface. *In vitro*, across all conditions tested in NanoBiT assays and analytical SEC, NC-interface assembly closely tracks the ability to form G-interfaces. These findings are consistent with earlier co-precipitation experiments of SEPT2, SEPT6, and SEPT7 showing destabilization of NC interactions upon G-interface perturbation^51^. Taken together, we propose a model in which G-interface formation functions as the gatekeeper step in septin protofilament assembly.

Our previous Cdc11 apo crystal structure, together with the cryo-EM, DEER, and MD analyses presented here, provides a structural basis for this ordered pathway. They define a nucleotide-triggered conformational switch within the SUE that coordinates competence for G- and NC-interface formation. In the absence of nucleotide, the Cdc11 SUE-βββ adopts a flexible, off-filament-axis conformation^32^ and helices α5 and α6 merge into a continuous, elongated helix incompatible with NC-interface formation (Fig. 5d). The cryo-EM structure with Cdc11 in the filament context shows that nucleotide binding, stabilized by the Cdc12-Cdc11 G-interface, repositions the β-meander and restores the canonical α5-α6 elbow, thereby generating an NC-competent conformation. Consistent with this, our DEER experiments show that nucleotide binding rigidifies the Cdc11 SUE-βββ, shifting it from a broad, interface-incompetent ensemble to a protofilament-like arrangement. MD simulations reveal that high SUE-βββ flexibility is a conserved feature of monomeric septins across species and that G-interface formation strongly stabilizes the SUE-βββ and bound nucleotides. Based on our experimental data, we propose a stepwise pathway for septin protofilament assembly (Fig. 6). In the apo ground state, the SUE-βββ adopts a highly flexible conformation that renders both G- and NC-interfaces incompetent for engagement. Transient nucleotide binding reshapes the conformational energy landscape and stabilizes G-interface-compatible SUE-βββ conformations. However, this intermediate is short-lived owing to labile nucleotide binding. G-interface formation with a compatible partner captures this conformation, locking in nucleotide binding and licensing subsequent NC-interface engagement, thereby enabling hetero-tetramer formation via α0-helix interactions and finally yielding octameric protofilaments.

**Fig. 6.**
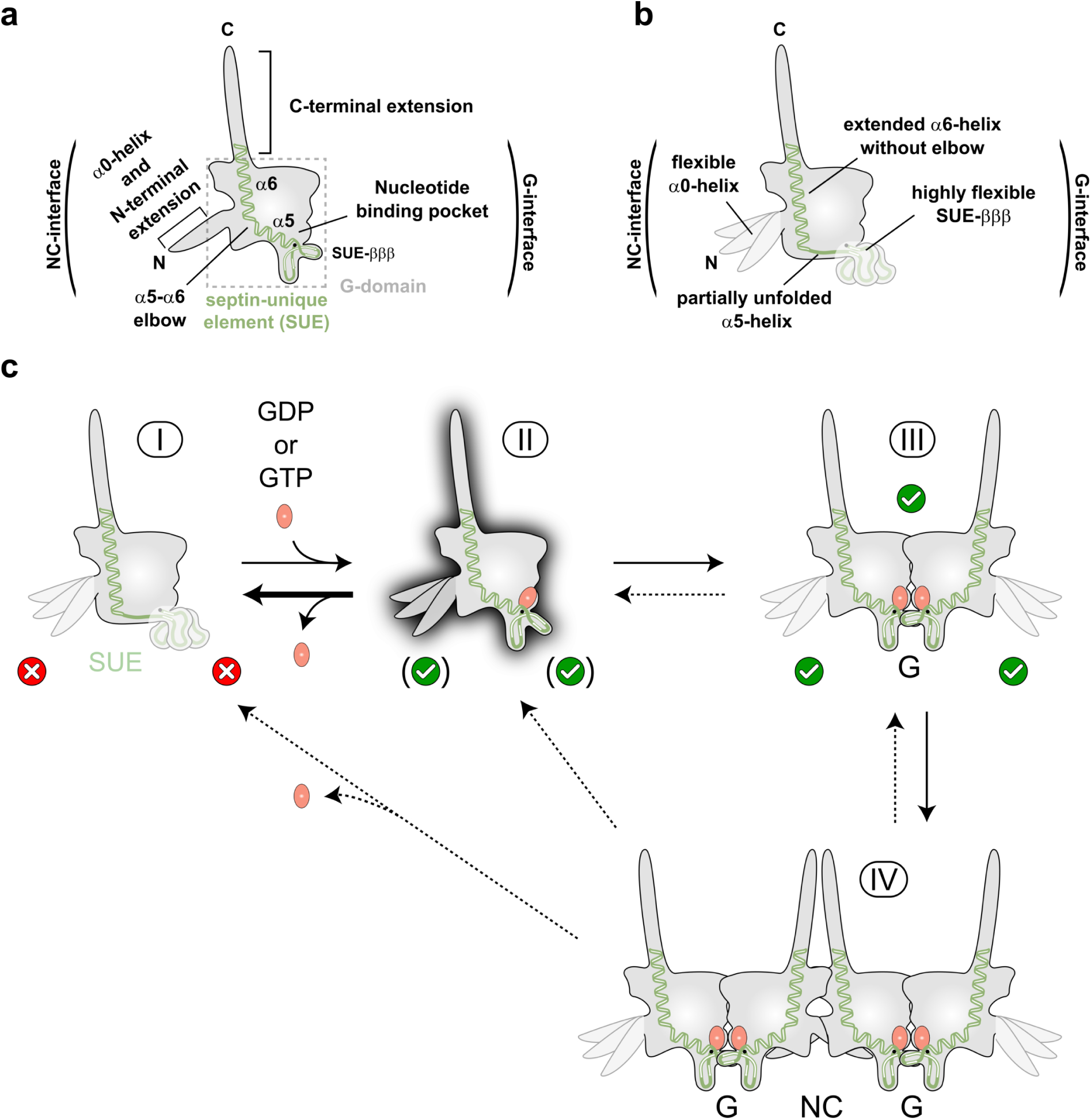
The ordered pathway of septin protofilament assembly. **a**, Cartoon representation of the G- and NC-interface-engaged nucleotide-bound conformation in assembled protofilaments. The α0-helix is rigidified by contacts with the adjacent NC-interface partner, the canonical SUE α5-α6 elbow is formed, and the SUE-βββ is oriented towards the filament axis and stabilized through interactions with the G-interface partner. **b**, Cartoon representation of the monomeric apo ground state with a highly flexible SUE-βββ and a mobile α0-helix. In this state, the α5-α6 elbow is lost, α5 is partially unfolded, and α6 is N-terminally extended into the space required for proper α0-helix positioning. This conformation is incompetent for both G- and NC-interface engagement. **c**, Proposed stepwise pathway of protofilament assembly. Starting from the monomeric apo state (I), transient nucleotide binding reshapes the conformational energy landscape and stabilizes a G-interface-compatible SUE-βββ conformation (II), but this intermediate remains short-lived owing to labile nucleotide binding. Formation of a G-interface dimer with a compatible, nucleotide-loaded partner captures this conformation and locks in nucleotide association (III). This allows subsequent NC-interface formation via α0-helix interactions, yielding hetero-tetramers (IV) and, ultimately, octameric protofilaments.

This mechanism is conserved from yeast to humans and provides a framework for understanding how mutations in the septin nucleotide-binding pocket disrupt filament assembly and cytoskeletal organization, as illustrated by P-Loop and G4 yeast mutants^21^, the SEPT12 D197N male infertility allele^23^, and numerous cancer-associated substitutions predicted to perturb nucleotide binding and interface stability^24^.

An important future goal is to elucidate how nucleotide-primed protofilaments transform into higher-order filaments and rings *in vivo*. In our *in vitro* system, formation of inside-out Cdc12-Cdc11-Cdc11-Cdc12 rods depends on the α0-helix and requires low ionic strength, whereas *in vivo* this process is likely enforced through cooperative inter-protofilament interactions, high local septin concentrations, and post-translational modifications. Many hydrophobic residues that stabilize NC-interfaces in internal subunits are replaced by charged residues in Cdc11^31^, consistent with a tunable balance of electrostatic and hydrophobic interactions modulated by post-translational modifications. Future studies capturing additional nucleotide states and dissecting the roles of membranes, binding partners, modifications, and inter-protofilament contacts will help to fully understand how septin assemblies are organized and regulated in both physiological and pathological contexts.

## Materials and Methods

### Plasmids and strains

All plasmids used in this study are listed in Supplementary Table 8. A list of the constructs used in the different figures is given in Supplementary Table 9. All constructs generated here were verified by sequencing. Point mutations were introduced via splicing by overlap extension PCR (SOE-PCR).

For expression of single septins in *E. coli*, open reading frames were amplified by PCR from existing plasmids and cloned as N-terminal His_6_- or His_6_-MBP-tagged fusions into the pET15b-derived vector pAc^27^ or into the first multiple cloning site of pETDuet-1 (Novagen). For NanoBiT experiments, LgBiT or SmBiT was fused C-terminally to septins via a 26-residue linker with the sequence (GGGGS)_4_-GGGGGS. Because SmBiT fusions expressed poorly in the absence of a solubility tag, septin-SmBiT constructs without an MBP tag were recloned into pE-SUMOpro Kan (Enzo Life Sciences) to introduce a C-terminal SUMO tag separated by an additional (GGGGS)_2_ linker. For SEPT6, SUMO was fused N-terminally directly to the septin without an intervening linker (plasmid #88).

For co-expression of yeast septin G-domain pairs in *E. coli*, compatible G-interface partners were cloned into the two multiple cloning sites of pETDuet-1 to enable bicistronic expression. In these constructs (plasmids #6–#8), Cdc10 or Cdc12 contained an N-terminal His_6_ tag, whereas the binding partner was fused to a C-terminal S tag; for construct #92 used in single-particle cryo-EM, the S tag was omitted by introducing a stop codon. Plasmids and strains used for yeast growth assays (plasmids #21–#65) were generated as described previously^27,31^.

For fluorescence microscopy, the open reading frames of CDC10, CDC3, CDC12, CDC11 and SHS1 were cloned as mNeonGreen fusions into pRS306^52^ under control of the MET17 promoter to generate plasmids #66–#70. For Cdc3 and Shs1, mNeonGreen was fused directly at the C-terminus; for Cdc10 and Cdc12, (GGGGS)_3_ and (GGGGS)_5_ linkers were inserted between the septin and mNeonGreen, respectively. For Cdc11, mNeonGreen was fused at the N-terminus via a (GGGGS)_3_ linker. These plasmids were linearized with StuI and transformed into the haploid yeast strain JD47^53^, resulting in integration at the URA3 locus by homologous recombination and restoration of a functional URA3 allele. For peroxisomal recruitment, the strains were subsequently transformed with an in-house-generated pRS304^52^ derivative (pRS304-X3) containing the homologous regions from the vector pCfB2189^54^ of the EasyClone 2.0 Yeast Toolkit for integration at site 3 on chromosome X. The homologous regions flanking the integration cassette were separated by an SfiI site used for linearization and genomic integration. The multiple cloning site of pRS304-X3 contained a chimeric sequence encoding Pex3_M1-K45_-mCherry for peroxisomal localization^55^ fused N-terminally via a (GGGGS)_3_ linker to variants of Cdc3 or Cdc12, and expression was controlled by the CUP1 promoter.

### Protein expression and purification

All constructs for protein purification were recombinantly expressed in *E. coli* LOBSTR-BL21(DE3)-RIL^56^. Transformed cells were grown overnight in Luria broth supplemented with chloramphenicol and the appropriate antibiotic, selecting for the expression plasmid. Pre-cultures were diluted 1:100 into super broth medium containing only the latter antibiotic for selection of the expression vector. Cells were grown at 37 °C to an OD600 of ∼1.2, induced with 1 mM IPTG, and expressed for ∼20 h at 18 °C. For expression of NanoBiT constructs, pre-cultures were diluted 1:100 into ZYP-5052 autoinduction medium^57^ supplemented with the appropriate antibiotic and grown for 5 h at 37 °C before transfer to 18 °C for ∼20 h. Cells were subsequently harvested by centrifugation, and pellets were stored at −80 °C.

All purifications followed the same workflow of cell lysis, immobilized metal affinity chromatography (IMAC), and preparative size-exclusion chromatography (SEC) described previously^27^ with modifications to the lysis and IMAC buffers depending on the construct. IMAC was generally performed at ambient temperature and SEC at 4 °C. All LgBiT-fused constructs except #86 were purified using an in-house-prepared HisTrap HP column (Cytiva) stripped of nickel ions and recharged with cobalt ions. All other constructs were purified using a 5 ml HisTrap Excel column (Cytiva).

Cell pellets from cultures co-expressing septins (#6–#8, #92) were resuspended in IMAC dimer-A buffer (50 mM Tris pH 8.0, 300 mM NaCl, 5 mM MgCl_2_, 15% (v/v) glycerol) supplemented with 0.1% Tween-20 and cOmplete protease inhibitor cocktail (Roche), and lysed by lysozyme treatment followed by sonication. The clarified lysate was loaded onto a 5 ml HisTrap Excel column (Cytiva), washed with IMAC dimer-A containing 3% IMAC dimer-B buffer (50 mM Tris pH 8.0, 300 mM NaCl, 5 mM MgCl_2_, 15% (v/v) glycerol, 200 mM imidazole), and proteins were eluted with 100% IMAC dimer-B.

Cell pellets from cultures expressing single septin constructs #3–#5, #11–#13, #16, #20, #80–#89 were resuspended in IMAC monomer-A buffer (50 mM Tris pH 8.0, 500 mM NaCl, 5 mM MgCl_2_) supplemented with 0.1% Tween-20 and cOmplete protease inhibitor. IMAC monomer-B buffer (50 mM Tris pH 8.0, 500 mM NaCl, 5 mM MgCl_2_, 200 mM imidazole) was used for washing and elution as described above. For constructs #93–#96, both buffers were supplemented with 5 mM β-mercaptoethanol.

For constructs #82–#86, IMAC monomer-B buffer was supplemented with 1 M MgCl_2_ to dissociate homodimers, as described previously^27^. For constructs displaying lower stability (#1, #2, #9, #10, #14, #17, #18, #90, #91), cell pellets were resuspended in IMAC lowsol-A buffer (50 mM Tris pH 8.0, 500 mM NaCl, 5 mM MgCl_2_, 50 mM L-Arg, 50 mM L-Glu, 15% (v/v) glycerol, 0.1% Tween-20) supplemented with cOmplete protease inhibitor. IMAC lowsol-B buffer (50 mM Tris pH 8.0, 500 mM NaCl, 5 mM MgCl_2_, 50 mM L-Arg, 50 mM L-Glu, 15% (v/v) glycerol, 0.1% Tween-20, 200 mM imidazole) was used for washing and elution. For constructs #15 and #19, both buffers contained 250 mM L-Arg and 250 mM L-Glu.

IMAC product peaks were pooled, supplemented with 5 mM dithiothreitol (DTT) if not eluted in buffer containing 5 mM β-mercaptoethanol, and subjected to preparative SEC on a HiLoad 16/600 Superdex 200 column (120 ml; Cytiva) equilibrated in SEC buffer (25 mM Tris pH 8.0, 300 mM NaCl). For purifications intended for nucleotide-occupancy determination, SEC buffer was supplemented with 5 mM MgCl_2_. Construct #84 was centrifuged immediately before SEC (5 min at 16,100 × g at 4 °C) due to its tendency to precipitate in 1 M MgCl_2_.

Purified proteins were analyzed by SDS-PAGE on 4-12% Bis-Tris gradient gels (Invitrogen) followed by Coomassie Brilliant Blue staining, and oligomeric states were assessed by analytical SEC using a Superose 6 Increase 10/300 GL column (24 ml; Cytiva) equilibrated and operated in SEC buffer containing 5 mM MgCl_2_ at 4 °C (Supplementary Fig. 9; Supplementary Fig. 10). Protein concentrations were determined by Bradford assay (construct #85) or absorbance at 280 nm using a NanoDrop ND-1000 spectrophotometer (Peqlab), with extinction coefficients and molecular weights calculated using the ExPASy ProtParam tool^58^.

### Determination of the nucleotide occupancy

Nucleotide occupancy was determined essentially as described previously^31^, with adjusted extinction coefficients and protein concentrations. Briefly, septin samples were diluted to 30 μM (dimer) or 60 μM (monomer) to yield a constant 60 μM monomer-equivalent concentration and incubated for 5 min at 95 °C to denature the protein. Precipitated protein was removed by centrifugation for 10 min at 16,100 × g and 4 °C, and 110 μl of the supernatant were adjusted with 20 mM Tris pH 8.0 to a final volume of 5.5 ml. Guanine nucleotides were separated by analytical anion-exchange chromatography on a MonoQ HR 5/5 column (1 ml; Cytiva), calibrated with known concentrations of GTP (purity ≥ 99%; Jena Bioscience) and GDP (purity ≥ 96%; Sigma-Aldrich), using a linear NaCl gradient (0-450 mM) over 18 column volumes. Peak areas corresponding to individual nucleotides were integrated in Origin v2025b (OriginLab). The relative nucleotide content per septin subunit was calculated from the determined amount of GTP and GDP, correcting the protein concentration for the absorbance contribution of bound guanine nucleotides at 280 nm, using the appropriate extinction coefficients for the respective oligomeric state of the septin sample and for the guanine nucleotide (7720 M^-1^ cm^-1^)^59^.

To assess the ability of monomeric septins to bind GTP or GDP after purification, proteins (100 μM for constructs #3–#5; 50 μM for constructs #1 and #2) were incubated for 1 h at 20 °C with an equimolar mixture containing a total of 3 mM GTP (purity ≥ 90%; Jena Bioscience) and GDP (purity ≥ 90%; Jena Bioscience) in SEC buffer supplemented with 5 mM MgCl_2_. Subsequently, the proteins were separated from unbound nucleotide using two 5 ml HiTrap columns connected in series (10 ml bed volume) equilibrated and operated with SEC buffer supplemented with 5 mM MgCl_2_. Septin-containing fractions were collected and their concentrations determined by absorbance at 280 nm using a NanoDrop ND-1000 spectrophotometer (Peqlab). Samples were then diluted to 60 μM for constructs #3–#5, or 30 μM for constructs #1 and #2. Subsequent treatment and analysis followed the procedure described above.

### Free phosphate detection (Malachite Green) assay

Malachite green assays were performed using the Malachite Green Phosphate Assay Kit (MAK307; Sigma-Aldrich) according to the manufacturer’s instructions. In brief, 80 µL of purified septin monomer or G-interface dimer (1.25 µM) was mixed with 20 µL GTP (500 µM) (purity ≥ 99%; Jena Bioscience) at 15 min intervals over a total time course of 105 min to monitor GTP hydrolysis. Reactions were performed in SEC buffer supplemented with 5 mM MgCl_2_ or 5 mM EDTA. The temperature was held constant at 20 °C, and reactions were terminated by adding the malachite green Working Reagent prepared from Reagents A and B as specified by the supplier, followed by incubation for 30 min at 20 °C in clear 96-well plates (Sarstedt). Absorbance was recorded at 620 nm on a Paradigm Multi-Mode Detection Platform Microplate Reader (Beckman Coulter). The phosphate standard provided with the kit was used to generate a standard curve and to convert absorbance values to free phosphate concentration. Apparent catalytic rate constants *k_cat,app_* were obtained by linear regression of free phosphate concentration versus time in GraphPad Prism v8.4.3 (Dotmatics). To establish an internal detection threshold for catalytic activity, maltose-binding protein (MBP) was used as a negative control. MBP was expressed in *E. coli* BL21(DE3) from plasmid pMAL-c5X (New England Biolabs) following the same protocol used for septin expression and purified over three consecutive 1 mL MBPTrap HP columns (total bed volume 3 mL; Cytiva) according to the manufacturer’s instructions. The control was assayed in SEC buffer supplemented with 5 mM MgCl_2_. The arithmetic mean and standard deviation (s.d.) of the obtained apparent catalytic constants for MBP were calculated, and the detection threshold was defined as the mean plus two SDs (mean + 2 × s.d.), following standard blank-based detection-limit practice^60^. Each replicate measurement of the other constructs was then classified as 1 (increased activity) if its *k_cat,app_* was greater than or equal to this threshold, or 0 otherwise. For each construct, the fraction of replicates with a value of 1 was computed, and constructs for which ≥ 50% of replicates exceeded the threshold were classified as catalytically active. Constructs and conditions with detectable activity were compared using one-way ANOVA followed by Tukey’s post hoc test.

To assess the substrate-concentration dependence, the same assay was performed over 60 min in 20 min intervals with final GTP concentrations ranging from 20 to 300 µM, and apparent catalytic rate constants versus initial GTP concentration were fitted in GraphPad Prism v8.4.3 using a Michaelis-Menten model. Care was taken not to consume more than 10% of the initially supplied substrate in any condition.

### Stopped-flow measurements

Stopped-flow fluorescence assays were performed on an SX20 stopped-flow spectrometer (Applied Photophysics) with fluorescence detection using a 470 nm LED for excitation and a FBH515-10 band-pass filter (CWL 515 nm, FWHM 10 nm; Thorlabs) for emission. Protein and nucleotide solutions were mixed in a 5:1 volume ratio in the stopped-flow experiment, and all concentrations are reported as final values after mixing. Measurements were carried out at an ambient temperature of ∼20 °C. Protein and nucleotide concentrations were adjusted for each construct to optimize signal-to-noise, and time courses were recorded with construct-specific sampling frequencies and total acquisition times to adequately resolve kinetic phases. Fluorescence traces were normalized for their initial value and plotted as *F(t)/F(0)*.

For association kinetics, constructs #1 and #2 were measured under pseudo-first-order conditions at final protein concentrations between 1.0 and 4.2 µM in the presence of 0.08 µM EDA-GTP-BDP-FL (Jena Bioscience), whereas construct #4 was assayed at final protein concentrations of 21 to 104 µM with 0.8 µM EDA-GTP-BDP-FL. Constructs #3 and #5 were measured at final concentrations of 104 µM protein with 2.5 µM EDA-GTP-BDP-FL or EDA-GDP-BDP-FL (Jena Bioscience). Specificity controls included 520 µM unlabeled GTP in addition to EDA-GTP-BDP-FL. Septin solutions were prepared in SEC buffer, and nucleotide solutions were prepared in SEC buffer supplemented with 30 mM MgCl_2_.

For dissociation kinetics, constructs #1 and #2 were freshly pre-equilibrated with EDA-GTP-BDP-FL at concentrations of 2 µM protein and 0.25 µM nucleotide, and dissociation was initiated by rapid mixing with a chase solution containing 2.5 mM GTP and 2.5 mM GDP. For construct #4, complexes were formed at 75 µM protein and 1.25 µM EDA-GTP-BDP-FL, and dissociation was triggered by mixing with 2.5 mM GTP and 2.5 mM GDP. All solutions used in dissociation experiments were prepared in SEC buffer supplemented with 5 mM MgCl_2_.

Average fluorescence traces of 4–6 technical replicates were normalized to the initial fluorescence and analyzed in GraphPad Prism v8.4.3. Dissociation traces of constructs #1 and #4 were fitted with a one-phase exponential decay model with linear drift,

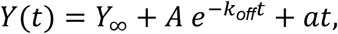

where *t* is time, *Y(t)* is the normalized fluorescence, *Y_∞_* is the fluorescence at long times, *A* is the amplitude, *k_off_* is the dissociation rate constant, and *a* is the linear drift term. For each construct, geometric means and asymmetric 95% confidence intervals (CIs) in log space of *k_off_* across biological replicates (independent protein preparations) were calculated.

Association traces were globally fitted over the protein concentration series of each biological replicate using a one-phase exponential association model with drift,

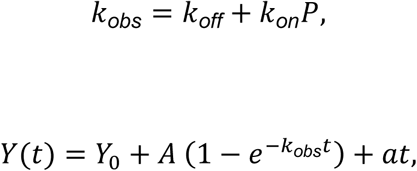

where *P* is the protein concentration, *k_on_* is the association rate constant, *k_obs_* is the observed first-order rate constant, *Y_0_* is the initial fluorescence, and *A* is the total association amplitude. In these fits, *k_off_* was constrained to the geometric mean obtained from dissociation measurements of the same construct. The geometric mean and asymmetric 95% CIs in log space of *k_on_* were then determined across biological replicates. An estimated *K_D_* was calculated as *k_off_/k_on_*, and CIs for *K_D_* were derived from the confidence limits of *k_off_* and *k_on_*, with the upper *K_D_* bound obtained from *k_off,upper_/k_on,lower_*, and the lower *K_D_* bound from *k_off,lower_/k_on,upper_*.

For construct #2, association and dissociation traces displayed biphasic behavior and were analyzed with a global induced-fit model of the form

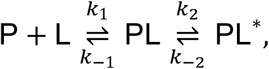

where *k_2_* and *k_−2_* denote the forward and reverse isomerization rates, respectively. Association and dissociation time courses were simultaneously fitted with double-exponential functions in which the observed rate constants are the analytical eigenvalues of the rate matrix for the two-step binding scheme. Because protein was present at least tenfold in excess over the labeled ligand, association was analyzed under an effective pseudo-first-order approximation in protein. In this regime, the free protein concentration *P* remains essentially constant during the experiment, and the two observed rate constants for association are given by

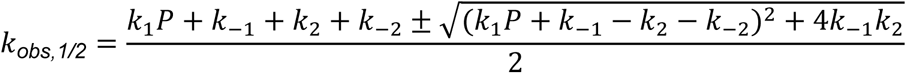

following the general treatment of induced-fit kinetics^61^, and the association time course is described by

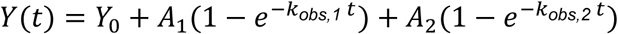

where amplitudes *A_1_* = (*Y_∞_* - *Y_0_*) × *f_fast_* and *A_2_* = (*Y_∞_* - *Y_0_*) × (1 - *f_fast_*) are partitioned by a dataset-specific fractional amplitude parameter *f_fast_*. For dissociation, complexes pre-equilibrated with labeled GTP were rapidly mixed with a large excess of unlabeled GTP. In this configuration, the free concentration of labeled ligand *L* is effectively driven to 0, so that dissociation of labeled nucleotide can be treated as pseudo-first-order. The observed rate constants at *L* = 0 become

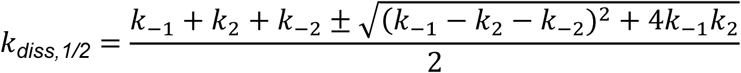

and the dissociation time course is described by

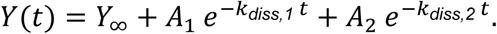

Each of six independent association concentration series was fitted globally together with four dissociation traces, yielding six independent estimates of the microscopic rate constants *k_1_*, *k_−1_*, *k_2_* and *k_−2_*. For each rate constant, the geometric mean and asymmetric 95% CIs in log space were calculated. An overall dissociation constant for the two-step induced-fit scheme^62^ was calculated as (*k_−1_* × *k_-2_*) / (*k_1_* × (*k_2_* + *k_−2_*)). CIs for *K_D_* were derived from the confidence limits of the kinetic constants, with the upper *K_D_* bound obtained from (*k_−1,upper_* × *k_−2,upper_*) / (*k_1,lower_* × (*k_2,lower_* + *k_−2,lower_*)) and the lower *K_D_* bound from (*k_−1,lower_* × *k_−2,lower_*) / (*k_1,upper_* × (*k_2,upper_* + *k_−2,upper_*)).

### Analytical gel filtration experiments

For analytical SEC, 100 µl of the samples were injected onto a Superose 6 Increase 10/300 GL column (24 ml; Cytiva) equilibrated and operated in SEC buffer containing 5 mM MgCl_2_ at 4 °C. Elution was monitored by UV absorbance at 280 nm, and peak positions and shapes were used to assess oligomeric state and complex formation under the different nucleotide conditions. The column was calibrated with the Bio-Rad Gel Filtration Standard #1511901, and the resulting standard curve was used to estimate apparent molecular masses of the eluting species. Because septin complexes are rod-shaped, size-exclusion calibration with globular protein standards becomes increasingly inaccurate with increasing rod length. Therefore, mass photometry was used to confirm the molecular weight of complexes larger than dimers after analytical SEC.

To assess G-interface dimer integrity under nucleotide-depleted conditions, 15 µM of purified complexes #6–#8 were incubated for 1 h at 30 °C in the presence or absence of 150 µM GTP and GDP. An additional sample contained the nucleotide mixture and 0.6 U of shrimp alkaline phosphatase (M0371; New England Biolabs).

Analytical gel filtration was also used to analyze septin complex formation under defined nucleotide conditions. Reaction mixtures (200 µl total volume) were prepared in SEC buffer supplemented with 5 mM MgCl_2_, except where indicated. For 1:1 mixtures, proteins were typically mixed at 10 µM each; for 1:1:1:1 mixtures, each protein was used at 7.5 µM. An exception was the SEPT7-SEPT9 combination (plasmids #83 and #89), where 7 µM SEPT7 were mixed with 35 µM SEPT9 to compete with SEPT7 homodimerization. Guanine nucleotides were added at a total concentration of 200 µM for 1:1 mixtures, or 300 µM for 1:1:1:1 mixtures. The following nucleotide conditions were tested: no nucleotide, GTP, GDP, equimolar GTP and GDP, equimolar GTP and GDP in the presence of 5 mM EDTA instead of MgCl_2_, GTPγS (Jena Bioscience), and GppNHp (Jena Bioscience). All samples were incubated overnight at 4 °C before analysis.

### Pulldown experiments

Pulldown assays were performed using amylose resin (New England Biolabs) to assess complex formation between tagged and untagged septin constructs under defined nucleotide conditions. Purified septin samples were prepared in SEC buffer. For each condition, 500 µl reactions were prepared containing 2 µM of the MBP-tagged septin construct and 10 µM of the interaction partner. Binding controls were included in which each protein was incubated alone in SEC buffer. Where indicated, the reactions were supplemented with either 5 mM MgCl_2_ or 5 mM EDTA, and with the following nucleotide conditions: no nucleotide, 50 µM GTP and 50 µM GDP, 100 µM GTP, 100 µM GDP, 100 µM GTPγS, or 100 µM GppNHp. Samples were incubated for 16 h at 4 °C.

For the pulldown, 50 µl amylose resin (New England Biolabs) were pre-equilibrated in SEC buffer supplemented with either 5 mM MgCl_2_ or 5 mM EDTA and incubated with the mixtures for 30 min at 4 °C on a rotating wheel. Subsequently, beads were washed three times with SEC buffer containing either 5 mM MgCl_2_ or 5 mM EDTA. Bound proteins were eluted by the addition of elution buffer (SEC buffer supplemented with 10 mM maltose). Beads were pelleted by centrifugation, and the supernatant was mixed with Laemmli buffer and subsequently analyzed by SDS-PAGE.

### NanoBiT experiments

Assays were performed on a Paradigm multimode detection platform (Beckman Coulter) in white 96-well polypropylene plates (Nunc MicroWell, catalog no. 267350; Thermo Fisher Scientific). Luminescence was measured using the Lum96 module (150 ms integration time) and recorded as relative light units (RLU). The high brightness of NanoLuc enabled the use of low septin-LgBiT concentrations (100 pM or 5 ng/ml for construct #85), which preserved protein stability and minimized GTP turnover during incubation by these subunits. For assays using yeast septins, samples were supplemented with 0.1% BSA (w/v), whereas assays using human septins contained 0.5% BSA (w/v) to reduce unspecific interactions and to increase protein stability.

In equilibrium titration experiments, LgBiT fusions were incubated with increasing concentrations of cognate G-interface partners fused to SmBiT in 100 µl total volume containing 200 µM guanine nucleotide and either 5 mM MgCl_2_ or 5 mM EDTA. Nucleotide conditions included GTP, GDP, an equimolar GTP and GDP mixture (100 µM/100 µM), and the non-hydrolyzable analogs GTPγS or GppNHp. LgBiT and SmBiT fusion proteins together with the indicated nucleotides were pre-incubated overnight at 20 °C. A nucleotide-free sample served to estimate nonspecific interactions between solubility tags and NanoBiT fragments. Immediately before measurement, fluorofurimazine (Selleckchem) was added to 10 µM from a 500 µM stock solution prepared in SEC buffer containing 5% DMSO.

Luminescence was normalized to the maximum signal per plate. To enable direct comparison across conditions, all nucleotide (GTP, GDP, GTP/GDP, GTPγS, GppNHp) and magnesium (MgCl_2_, EDTA) conditions were assayed together on the same plate. Data were analyzed in GraphPad Prism v8.4.3 using a one-site total binding model that incorporated specific and nonspecific components. Nonspecific binding was first estimated from nucleotide-free controls by linear fitting (yielding background and slope parameters), which were then fixed during fitting of total binding data, allowing only specific binding parameters to vary. Apparent *K_D_* values and asymmetric 95% CIs were obtained by fitting individual biological replicates and computing geometric means and asymmetric 95% CIs in log space, followed by back-transformation.

To monitor NC-interface formation, assays used fixed concentrations of 100 pM septin-LgBiT, 1 µM septin-SmBiT, and 5 µM of each G-interface partner (10 µM when both subunits shared the same G partner). All other procedures, including incubation, nucleotide conditions, measurement, and normalization, followed those described for measurements investigating G-interface formation. Normalized luminescence was analyzed in R v4.5.2 using the *tidyverse*, *nlme*, *car*, and *emmeans* packages. For each NanoBiT dataset, responses were modeled with a three-way generalized least squares (GLS) model (normalized RLU ∼ G-partner × buffer × nucleotide) incorporating a varPower variance structure to accommodate mean-variance relationships, and Type III likelihood-ratio tests were used for omnibus effects. Significant interactions were followed by simple-effects analyses on estimated marginal means to test three a priori predictions: (i) different nucleotides elicit distinct responses when G-partner is present; (ii) nucleotide-dependent differences disappear in the absence of G-partner, indicating a requirement for G-interface formation; and (iii) Mg^2+^ modulates these effects. Within each buffer (MgCl_2_ or EDTA), nucleotide effects were examined separately for G-partner present versus absent, and G-partner effects were compared within each nucleotide-buffer combination. Pairwise nucleotide contrasts were corrected using Tukey adjustment, whereas contrasts comparing G-partner versus no G-partner were Holm-adjusted. Additionally, for the Cdc3-Cdc12 NC-interface, a small set of preplanned single-protein contrasts (Cdc10, Cdc11, or Shs1 versus no G-partner at MgCl_2_ + GTP/GDP) was tested using Holm-adjusted comparisons of estimated marginal means.

For measuring septin interaction kinetics, septin-LgBiT fusion protein was diluted to 100 pM (or 5 ng/ml for construct #85) in SEC buffer supplemented with 5 mM MgCl_2_ and 3 mM nucleotide (GTP or GDP), and incubated for 15 min at ambient temperature to allow nucleotide binding. An identical sample without nucleotide was prepared in parallel. Septin-SmBiT fusion proteins were prepared at variable concentrations from a concentrated stock by 3:5 serial dilution to yield 12 different SmBiT concentrations; the absolute concentration range depended on the construct and nucleotide state.

Immediately before the measurement, the LgBiT solutions were supplemented with 20 µM fluorofurimazine substrate from a 10 mM stock solution prepared in 100% DMSO. Reactions were initiated by rapid mixing of equal volumes (50 µl) of the septin-LgBiT and septin-SmBiT solutions, and luminescence was recorded over time at ambient temperature on the same instrument and with the same settings as described above. The time resolution and total recording duration were adjusted for each construct and nucleotide condition to capture the association. Luminescence time courses were analyzed globally using a custom Python workflow (lmfit v1.3.4) that implements a one-phase association model with explicit specific and nonspecific components for each SmBiT concentration *i*. For each concentration, apo (no nucleotide) traces were fit to a single-exponential association function to estimate nonspecific binding contribution

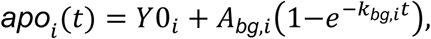

while specific traces (3 mM GTP or GDP) were fit to

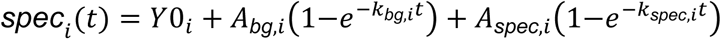

where *t* denotes time, *Y0_i_* is the extrapolated signal at *t* = 0, *A_bg,i_* and *k_bg,i_* describe nonspecific binding, and *A_spec,i_* and *k_spec,i_* describe specific complex formation. For a given concentration *i*, the kinetic parameters *Y0_i_*, *A_bg,i_*, *k_bg,i_* were shared between apo and specific traces, while *A_spec,i_* and *k_spec,i_* were fitted exclusively to the specific traces. All model parameters were optimized simultaneously by nonlinear least-squares fitting. The obtained *k_spec,i_* values were subsequently plotted against the corresponding septin-SmBiT concentrations in GraphPad Prism v8.4.3 and either fitted by linear regression (Cdc10-Cdc3) or by a hyperbolic model

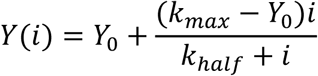

for Cdc12-Cdc11, Cdc12-Shs1 and SEPT9-SEPT7. In this model, *Y(i)* corresponds to *k_spec_* at septin-SmBiT concentration *i*, *Y_0_* is the y-intercept, *k_max_* is the fitted maximum corresponding to *k_spec,∞_*, and *k_half_* the septin-SmBiT concentration at which *Y* = (*Y_0_* + *k_max_*) / 2.

### Yeast viability spot assay

Assays were performed using the haploid yeast strain JD47^53^ in the CDC10, CDC3, CDC12, CDC11, and SHS1 deletion background essentially as described previously^27,31^. Each deletion strain carried a wild-type copy of the respective septin on a URA3-marked centromeric plasmid. Test constructs (wild-type or mutant septins) were introduced on centromeric plasmids under control of the PMET17 promoter. Overnight cultures grown in YPD were serially diluted (tenfold) and spotted onto plates containing 5-fluoroorotic acid (5-FOA) to select against the URA3 plasmid, and onto control plates. Plates were incubated at 30 °C or 37 °C. Relative growth on 5-FOA plates served as the readout for functional complementation by the investigated construct.

### Mass photometry

Septin complexes were assembled in SEC buffer supplemented with 5 mM MgCl_2_, 1 mM GTP, and 1 mM GDP as described above. Assembled complexes were separated by analytical SEC and supplemented with an equimolar mixture of 500 µM GTP and 500 µM GDP for storage. Protein concentrations were determined by Bradford assay.

Mass photometry measurements were performed on a TwoMP instrument (Refeyn Ltd.) to assess mass distributions and oligomeric states. Sample well cassettes (RD501078; Refeyn Ltd.) were mounted on clean microscope coverslips (MP-CON-41001; Refeyn Ltd.). For each measurement, 16–19 µL of mass photometry buffer (25 mM Tris pH 8.0, 300 mM NaCl, 5 mM MgCl_2_, 500 µM GTP, 500 µM GDP) was added to stabilize autofocus, followed by 1–4 µL of protein sample (final concentration 20–25 nM in 20 µL). Movies (60 s) were recorded using standard acquisition settings in AcquireMP (Refeyn Ltd.). The instrument was calibrated with a MassFerence P1 calibrant (Refeyn Ltd.).

Data were analyzed in Origin v2025b using Gaussian peak fitting (GaussAmp global fit function). Histogram bin width was set to 5 kDa. The number of fitted curves corresponded to all theoretically possible oligomeric intermediates, assuming molecular weights of ∼80 kDa for MBP-tagged subunits and ∼40 kDa for untagged subunits. Expected peak positions (e.g., 40-240 kDa for a tetrameric 1:1:1:1 complex containing two MBP-tagged and two untagged subunits) were constrained within ±10 kDa of the theoretical values, with a shared peak width of 5-25 kDa.

### Mass spectrometry

Septin complexes assembled and separated as described above were precipitated employing methanol/chloroform extraction following well-established protocols^63^. Each pellet was dissolved in 15 µl ammonium bicarbonate. Samples were reduced with 5 mM DTT (AppliChem) for 20 min at ambient temperature, alkylated with iodoacetamide (Sigma-Aldrich) for 20 min at 37 °C, and digested overnight at 37 °C with trypsin at a 1:50 enzyme-to-protein ratio.

Digests (15 µl) were analyzed on an LTQ Orbitrap Elite mass spectrometer (Thermo Fisher Scientific) online coupled to an UltiMate 3000 RSLCnano system (Thermo Fisher Scientific) as described previously^64^, with the following modifications: The column was equilibrated for 5 min in 5% solvent B (solvents A: 0.1% formic acid; solvent B: 86% acetonitrile, 0.1% formic acid). This was followed by different elution steps, in which the percentage of B was first raised from 5 to 15% in 5 min, followed by an increase from 15 to 40% B in 105 min. The 20 most intense ions from the survey scan were selected for CID fragmentation. Singly charged ions were rejected, and m/z of fragmented ions were excluded from fragmentation for 60 s. MS2 spectra were acquired in the linear ion trap at rapid scan speed.

Database search was performed using MaxQuant v.2.4.13.0^65^. For peptide identification, the built-in Andromeda search engine^66^ was employed to correlate MS/MS spectra with the respective protein sequences and the MaxQuant contaminant database. Carbamidomethylation of cysteine was set as a fixed modification, and oxidation of methionine and protein N-terminal acetylation as variable modifications. iBAQ quantification with log fit and charge normalization were enabled. False discovery rates for peptide and protein identifications were controlled at 1%. Quantification was based on iBAQ values.

### Fluorescence microscopy

Fluorescence microscopy was performed on an Axio Observer Z.1 spinning-disk confocal microscope (Zeiss) equipped with an Evolve512 EMCCD camera (Photometrics), a Plan-Apochromat 100×/1.4 NA oil DIC objective, and 488-, 561-nm diode lasers (Zeiss). Images were acquired using ZEN 2 software (Zeiss).

Cells were grown overnight in appropriate selection medium supplemented with 70 µM methionine and 50 µM CuSO_4_, diluted into 3 ml fresh medium, and cultured for 2 h at 30 °C to mid-log phase. Immediately before imaging, 1.5 ml of culture were harvested, resuspended in 100 µl sterile water, and 3.5 µl of the suspension were mounted on a glass slide under a coverslip. Z-stacks of 10 optical sections separated by 0.26 µm were collected in both the mNeonGreen and mCherry channels using the 100× objective.

Image analysis was carried out in Fiji^67^ (ImageJ v2.16.0/1.54p). Maximum-intensity projections were generated for each channel, and only G2/M cells were included for quantification. Peroxisomal signal was quantified by drawing an elliptical region of interest (ROI) around an mCherry-labeled peroxisome to capture the brightest pixels in the projection; a second ROI of identical size and shape was placed in an adjacent cytoplasmic region lacking visible mCherry enrichment, and avoiding the bud-neck region with mNeonGreen-labeled septins. For each peroxisome-cytoplasm pair, the maximum intensity within the ROIs was quantified in both channels and subsequently processed according to the following formula:

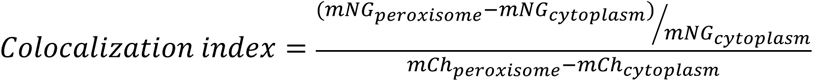

Peroxisomal intensities were background-corrected by subtracting the corresponding cytoplasmic maximum, and mNeonGreen (mNG) recruitment to peroxisomes was normalized to septin expression (cytoplasmic mNG signal) and peroxisome abundance (background-corrected mCherry (mCh) signal) to yield a colocalization index for each peroxisome. Data were analyzed in GraphPad Prism v8.4.3 using a nonparametric Kruskal-Wallis test followed by Dunn’s multiple comparison test.

### SEC-MALS measurements

SEC-MALS experiments were carried out on a Waters Alliance iS HPLC system controlled via HPLC CONNECT v4.0.4.115 and coupled to a DAWN multi-angle light-scattering detector (Waters | Wyatt Technology) and an Optilab differential refractive index detector (Waters | Wyatt Technology). Proteins were separated at ambient temperature on a Superdex 200 Increase 10/300 GL column (24 ml; Cytiva) operated at 0.75 ml min^−1^ in either low-salt buffer (25 mM Tris pH 8.0, 25 mM NaCl, 0.02% (w/v) NaN_3_) or standard-salt buffer (25 mM Tris pH 8.0, 300 mM NaCl, 0.02% (w/v) NaN_3_). UV absorbance at 280 nm and the dRI signal were recorded in parallel, and protein concentration calculated from the dRI trace was combined with light-scattering data to obtain absolute molar masses in ASTRA v8.3.1.21, using a dn/dc value of 0.185 ml g^−1^ for all proteins. Purified complexes at 1.5–2.2 mg ml^−1^ were injected at 10, 25, 50, or 100 µl per run to examine potential concentration-dependent changes in oligomeric state.

### Cryo-EM sample preparation and data collection

Cdc12-Cdc11 septin dimers (construct #92) were buffer-exchanged into low-salt buffer using 10 kDa molecular weight cutoff centrifugal filter units (Cytiva) by repeated concentration and dilution with low-salt buffer. The solution was incubated overnight at 4 °C to promote inside-out tetramer formation. Samples were clarified by centrifugation (10,000 × g, 10 min, 4 °C) to remove aggregates. Holey gold grids (UltrAuFoil R1.2/1.3) were glow-discharged for 60 s. Aliquots (3–4 µl) of septin samples (∼10 mg ml^−1^ for optimal grids) supplemented with 0.025% (w/v) CHAPSO to reduce preferred orientation were applied under controlled conditions (4 °C, 100% humidity) using a Vitrobot Mark IV (Thermo Fisher Scientific). Excess liquid was blotted for 6 s (blot force 5) before plunge-freezing in liquid ethane. Grids were clipped into autogrid cartridges, screened for ice thickness and particle distribution, and high-quality grids (medium-thin vitreous ice, homogeneous particle spread on UltrAuFoil supports) were stored in liquid nitrogen until data collection.

The Titan Krios G4 cryo-TEM (Thermo Fisher Scientific) was operated in nanoprobe EFTEM mode at an accelerating voltage of 300 kV and a magnification of 215,000×, corresponding to a calibrated physical pixel size of 0.572 Å at the specimen level. Images were recorded using a Falcon 4i direct electron detector (Thermo Fisher Scientific) in electron-counting mode with a 10 eV energy-filter slit, using EPU software (Thermo Fisher Scientific) in Faster acquisition (AFIS) mode. Dose-fractionated movies (8,195) with a total exposure of 40 e^−^ Å^−2^ were recorded with an exposure time of 1.76 s per movie and a focus step of 0.1 µm over a defocus range from −0.5 to −2.2 µm. Data were acquired using an aberration-free image shift to target five image areas per hole, selecting regions with suitable ice thickness and particle density while monitoring drift and image quality in real time.

### Single particle cryo-EM image processing and model building

The processing of movies was conducted in cryoSPARC v4.7.0 using patch motion correction and patch CTF estimation^68^. Initially, particles were selected using a blob picker, then extracted with a twofold Fourier downsampling factor, and subsequently subjected to 2D classification. In the initial iteration of the reconstruction process, a 3.4 Å map was obtained from a set of 84,000 particles, after a non-uniform refinement^69^. This map was then utilized as input for template-based particle picking, and the resulting hits were combined with the initial blob-picked particles. The presence of duplicates was checked, and the particles were extracted using a factor of 4 Fourier downsampling, followed by 2D classification. The sorting of particles was achieved through the implementation of heterogeneous refinement, with obvious bad particles and artefacts serving as exemplars of poor outcomes. This process resulted in the identification of 421,937 particles. The most optimal particle ensemble was then refined to a 2.74 Å reconstruction from 351,177 particles through implementation of a multifaceted approach. This approach encompassed the division of the dataset into 69 exposure groups, the application of reference-based motion correction, and subsequent non-uniform refinement. The map exhibits an absence of distinct density in the NC-interface. Movement along the complex axis has been shown to lead to conformational changes (Supplementary Fig. 6), as demonstrated in a variability analysis which employs superposed states^70^. This analysis comprises 20 states and a filter resolution of 5 Å. Within the analysis, representative states (1, 10, and 20) along the first two principal eigenvectors were reconstructed in cryoSPARC, and the Cdc12-Cdc11-Cdc11-Cdc12 model was subjected to a single round of real-space refinement in Coot against each map^71^. The implementation of focused 3D classification, accompanied by the utilization of a mask surrounding the NC-interface, has resulted in the identification of a sub-population of 70,904 particles, which exhibit uniform conformation of the α0-helix. The final particle sets were introduced to *C*_2_ via the Volume Alignment Tool and subjected to nonuniform refinement with minimization over a per-particle scale, optimized per-particle defocus, and per-group CTF parameters switched on, subjected to local refinement. The initial reference was low-pass filtered at 3.5 Å. During the refinement process, per-particle scale factors were subjected to optimization, while translational and rotational searches were confined to 5 Å and 10°, respectively. At each iteration, both rotations and shifts were subjected to recentering. The resulting reconstruction achieved a resolution of 2.52 Å, as determined by the gold-standard FSC criterion of 0.143. The full processing scheme is displayed in. Supplementary Fig. 11. The AlphaFold-predicted model of Cdc12-Cdc11-Cdc11-Cdc12 was subsequently integrated into the density in UCSF ChimeraX, manually built in Coot, and refined in real space using PHENIX^70–74^. The water molecules were constructed following the sharpened map output of cryoSPARC. The quality of the models was assessed using MolProbity. The collection and refinement statistics of the data are summarized in Supplementary Table 10. The cryo-EM map around different structural elements in Cdc12 and Cdc11 is shown in Supplementary Fig. 12

### Molecular Dynamics (MD) simulations

All-atom MD simulations were performed using energy-minimized septin dimer structures prepared in our previous work^27,75^. Briefly, simulations with human septins employed the G-interface dimers SEPT6-SEPT2 (PDB 6UPA) and SEPT7-SEPT7 (PDB 6N0B), truncated to their G-domains, with unresolved regions modeled in SWISS-MODEL, termini capped, protonation states assigned at pH 7, and hydrogen-bond networks optimized. Structures were relaxed in Maestro with the OPLS4 force field^76^ before conversion for CHARMM36m compatibility using the CHARMM-GUI webserver^77,78^. For simulations with yeast septins, the Cdc10-Cdc3, and Cdc12-Shs1 G-interface dimers were extracted from PDB 9GD4. The models were truncated to their G-domains and completed by homology modeling of missing segments in MODELLER^79^; energy minimization and protonation/hydrogen-bond optimization followed the same Maestro and CHARMM-GUI pipeline as outlined above. For the present study, only fully nucleotide-loaded dimers (GDP/GDP:Mg/GTP:Mg as in the crystal structures) and their corresponding apo dimers (all nucleotides removed) were simulated; monomers were simulated upon removal of the atomic coordinates belonging to the binding partner.

All simulations were run in GROMACS v2022.2^80^ using the CHARMM36m force field^78^ and the CHARMM-modified TIP3P water model^75,81,82^ on the JUSTUS2 high-performance cluster at Ulm University. Each construct was placed in a dodecahedral water box with at least 1.5 nm between protein and box edge, charge neutralized, and supplemented with 150 mM NaCl for human septins and 300 mM KCl for yeast septins. Bonds to hydrogens were constrained with the LINCS algorithm^83,84^, a 2 fs time step was used, and long-range electrostatics were treated with particle-mesh Ewald^85,86^ (real-space cutoff 1.2 nm), and Lennard-Jones interactions were smoothly switched off between 1.0 and 1.2 nm. After steepest-descent energy minimization, systems were equilibrated for 50 ps in the NVT ensemble (velocity-rescaling thermostat^87^; 310 K for human, 300 K for yeast) followed by 500 ps in the NPT ensemble (stochastic cell-rescaling barostat^88^; 1 bar) with positional restraints of 1000 kJ mol^−1^ nm^−2^ on all heavy atoms in both equilibration steps. Production runs were performed in the NPT ensemble using the Parrinello-Rahman barostat^89^ at the respective target temperature without restraints. For each system, ten independent 700 ns trajectories with different Maxwell-Boltzmann velocity seeds were generated. For the dimeric systems Cdc10-Cdc3 and Cdc12-Shs1, the five 700 ns trajectories from our previous study^27^ were reused and combined with five newly generated 700 ns replicas per system, yielding ten independent trajectories per nucleotide state for analysis.

To quantify local structural flexibility of the SUE-βββ or of nucleotides in the binding pockets, simulation frames were aligned to the initial conformation by superposition of all secondary structure elements excluding the α5′-helix and the SUE-βββ, and the backbone RMSD of SUE-βββ residues or the RMSD of all non-hydrogen atoms in the nucleotide was calculated per trajectory in GROMACS v2022.2^80^. The 95^th^-percentile RMSD per trajectory (RMSD95) was used as the primary response variable, and all statistical analyses were performed in R v4.5.2. For SUE-βββ flexibility, five pre-specified contrasts were evaluated per subunit using two-sided Wilcoxon rank-sum tests with Hodges-Lehmann estimators and asymmetric 95% CIs: dimer:GXP vs. dimer:apo, monomer:GXP vs. monomer:apo, dimer:GXP vs. monomer:GXP, dimer:apo vs. monomer:apo, and dimer:GXP vs. monomer:apo. *P*-values were adjusted within each subunit by the Holm method. Directional consistency of these effects across the eight subunits was assessed using one-sided sign tests on the per-subunit Wilcoxon estimates and one-sided Wilcoxon signed-rank tests on the per-subunit effect sizes (testing whether the median stabilizing effect is less than zero). For quantification of nucleotide flexibility, a single dimer vs. monomer contrast was evaluated per subunit with *P*-values adjusted across all eight subunits by the Benjamini-Hochberg method.

### Circular dichroism (CD) measurements

Samples used in Double Electron-Electron Resonance (DEER) spectroscopy were analyzed with CD spectroscopy to validate structural integrity. Purified constructs were stored in SEC buffer supplemented with 5 mM DTT and freshly desalted into CD buffer (300 mM NaF, 10 mM sodium phosphate pH 7.5) using a NAP-5 column (Cytiva). CD spectra were recorded at 20 °C on a Jasco J-810 spectropolarimeter using 0.1 mg/ml protein in a 1 mm path-length quartz cuvette (SUPRASIL 110-QS; Hellma Analytics). CD, high-tension, and absorbance signals were collected over a wavelength range of 260-185 nm with a data pitch of 0.5 nm, a bandwidth of 1.0 nm, and a digital integration time of 1 s. Spectra were acquired in continuous scanning mode at 50 nm/min, starting immediately after sample loading. For each sample, five scans were averaged, and corresponding buffer spectra were recorded and subtracted before further analysis.

### Isothermal titration calorimetry (ITC)

Measurements were performed on a MicroCal PEAQ-ITC instrument (Malvern Panalytical) at 20 °C. Purified Cdc11 (construct #4) was loaded into the sample cell at 100 µM in SEC buffer supplemented with 5 mM MgCl_2_, and GTP (purity ≥ 95%; Sigma-Aldrich) at a concentration of 2 mM prepared in the identical buffer was titrated from the syringe in 19 injections (first injection 0.4 µl, subsequent injections 2.0 µl) at a reference power of 10 µcal s^-1^; the first injection was excluded from analysis. Heats of dilution from GTP-into-buffer control titrations were subtracted from the raw data, and the corrected isotherms were fitted in the PEAQ-ITC analysis software v1.41 to a single-site binding model with the stoichiometry fixed at n = 1.

### Spin labeling

Spin labeling was carried out using MTSSL (AdipoGen Life Sciences) on septin constructs #93–#96. For constructs #93, #95, and #96, samples were used directly after preparative SEC in SEC buffer. For construct #94, preparative SEC was performed in SEC buffer supplemented with 5 mM DTT; immediately before spin labeling, this sample was desalted into SEC buffer using a PD-10 column (Cytiva). Proteins were incubated with a 20-fold molar excess of MTSSL overnight at 4 °C. For construct #94, the unreacted spin label was removed by a second preparative SEC step in SEC buffer supplemented with 5 mM MgCl_2_, and the peak corresponding to the monomeric species was collected, concentrated using 10 kDa cut-off centrifugal filters (Amicon), and adjusted to DEER buffer (300 mM NaCl, 25 mM Tris pH 8.0, 5 mM MgCl_2_, 30% glycerol). The remaining constructs were directly desalted into DEER buffer using two 5 ml HiTrap columns (Cytiva) connected in series (10 ml bed volume), equilibrated and operated with DEER buffer. Labeled samples were then split into two equal fractions, one supplemented with 5 mM GTP and the other diluted with an equivalent volume of DEER buffer.

### Determination of spin labeling efficiency

Spin-labeling efficiencies were determined from room-temperature continuous-wave EPR spectra recorded at X-band on a Bruker EMXnano benchtop spectrometer using 0.316 mW microwave power and 0.2 mT modulation amplitude at 100 kHz. Samples were measured in quartz glass tubes with 1 mm inner diameter. For constructs #93, #95, and #96, spectra were recorded using a single scan, and spin concentrations were determined by comparing the double integrals of the protein spectra with the double integral of a 200 µM aqueous TEMPONE standard (Sigma-Aldrich). For construct #94, three-scan spectra were recorded, and spin concentrations were determined by absolute spin counting using the quantitative EPR routines in Xenon software (Bruker). Labeling efficiencies for all constructs were calculated relative to protein concentrations determined by UV absorbance at 280 nm using a NanoDrop ND-1000 spectrophotometer (Peqlab). The same samples were also used for pulsed EPR experiments. Calculated labeling efficiencies are summarized in Supplementary Table 11.

### Pulsed EPR spectroscopy and four-pulse DEER

Pulsed EPR measurements were performed in Q-band at 80 K on an ELEXSYS E580 spectrometer (Bruker) using a dielectric ring Q-band resonator (model EN5107D2; Bruker). The cavity was cooled by a gas-flow cryostat with liquid nitrogen (CF935; Oxford Instruments), and the temperature was regulated by a PID controller (ITC4; Oxford Instruments). Microwave pulses at Q-band frequency (*ν_MW_* ≈ 34 GHz) were generated using a Super QFTu-EPR bridge (Bruker). The microwave pulses were amplified with a 50 W solid-state amplifier (AMGHz; Bruker). The following parameters were used for constructs #93, #95, and #96. The pulse lengths were set to *π*/2 = 18 ns and *π* = 36 ns at 0 dB attenuation. Echo-detected field-swept spectra (EDFS) were recorded in a magnetic field range from 1180–1220 mT by applying a standard Hahn-echo sequence with a pulse separation time *τ* = 200 ns. For the four-pulse DEER experiments, the pump pulse (*π_pump_* = 36 ns) was applied at the maximum of the nitroxide spectrum, and the observer pulses were high-field-shifted by ∼1.8 mT, corresponding to *Δν* ≈ 47–57 MHz (*ν_pump_ = ν_obs_ + Δν*) across acquisitions. For construct #94, the resonator was fully overcoupled; observer pulses were generated using the MPFU channels with pulse lengths of *π*/2 = *π* = 32 ns, pump pulses were formed using an incoherent ELDOR unit (E580-400U; Bruker) with *π_pump_* = 14 ns, and the frequency offset was set to *Δν* = +80 MHz.

A standard *π*/2 − *τ_1_* − *π* − *t* − *π_pump_* − *(τ_1_ + τ_2_* − *t)* − *π* − *τ_2_* sequence was used to acquire DEER time traces. The pump pulse position was incremented in steps of *Δt* = 8–16 ns, and the echo integration gate was positioned symmetrically around the refocused primary echo using quadrature detection. A two-step phase cycle [+(+x)−(−x)] applied on the first observer *π*/2 pulse was used to eliminate trigger offsets. For constructs #93, #95, and #96, *τ_1_* was set to 200 ns, and *τ_2_* was set according to the desired dipolar trace length and signal-to-noise ratio (SNR). The shot repetition time was set to 3 ms. For construct #94, *τ_1_* was selected based on the decay of the refocused primary echo for the chosen *τ_2_* value^90^, yielding *τ_1_* = 2.4 µs and *τ_2_* = 3.8 µs for both nucleotide states; *τ_2_* was chosen to observe at least two full dipolar oscillations while balancing background separation against the modulation depth-to-noise ratio, and the shot repetition time was set to 1.5 ms. The total measurement time for each sample ranged from 4 to 12 h, depending on the length of the time trace and SNR. Per-sample acquisition parameters are summarized in Supplementary Table 12.

### DEER data analysis

DEER time traces were analyzed using ComparativeDeerAnalyzer 2.0 (CDA2.0)^91^ within DeerAnalysis2022^92^ to obtain non-parametric distance distributions *P(r)* for constructs #93–#95; background-corrected DEER form factors and fits are shown in Supplementary Fig. 13a–c. In CDA2.0, these distributions are accompanied by calibrated 95% confidence bands. For construct #96, DEERNet^93^ background reconstruction was not used because the automatically generated backgrounds were considered unreliable. Instead, the distance distributions were obtained by Tikhonov regularization in DeerAnalysis2022^92^, with the regularization parameter *α* selected by the L-curve corner criterion; the corresponding form factors and fits are shown in Supplementary Fig. 13d. The robustness of the distribution was assessed by varying the starting time of the background fit and evaluating whether the main features of *P(r)* were qualitatively preserved; uncertainty bands are derived from the resulting spread in *P(r)*. Analysis parameters and resulting distance distribution statistics for all constructs are summarized in Supplementary Table 13. Reference distance distributions were simulated from the cryo-EM structure using the spin-label site-scan and labeling functionality in MMM v2024_1^94^, introducing MTSSL side-chain rotamers at the experimental sites at ambient temperature (298 K).

Multilateration was performed in MMMx v2023_2^95^ using the full CDA2.0 consensus distance distributions for constructs #93–#95 and the Tikhonov-regularized distance distribution for construct #96 as input constraints to obtain three-dimensional localization volumes. Cumulative mass-fraction curves were calculated using a custom Python script (NumPy v2.1.2, mrcfile v1.5.4, BioPython v1.84) by integrating the probability density map within unions of spheres centered on nitroxide N-O midpoints from the MMM rotamer library, considering only rotamers with non-zero occupancy. Sphere radii were incremented in 0.5 Å steps, and for each radius the fraction of total map density contained within the spheres was computed, yielding a cumulative mass-fraction versus distance curve for the apo, GTP-bound, and simulated ensembles.

### Figure preparation

Schematic representations of septin subunits and protofilaments were generated using Inkscape v1.4.3. Cartoon elements depicting the characteristic septin subunit shape were adapted from Grupp & Gronemeyer, *Biological Chemistry* 404, 1–13 (2023), with permission from De Gruyter. These elements appear across multiple figures as a consistent visual system representing individual septin subunits in monomeric, dimeric, tetrameric, hexameric, and octameric configurations with modified coloring and layout. All graphs were assembled using GraphPad Prism v8.4.3, Origin v2025b, and R v4.5.2. Panels showing protein structures were prepared in PyMOL v3.1.6.1^96^ (Schrödinger) and ChimeraX v1.11.1^74^.

### Statistics

Unless stated otherwise, data are presented as arithmetic means ± s.d., arithmetic means with symmetric 95% confidence intervals, or geometric means with asymmetric 95% confidence intervals calculated in log space and back-transformed (for quantities expected to follow log-normal distributions), as specified in the figure legends. Statistical analyses were performed in R v4.5.2 using the *nlme*, *emmeans*, *car*, and *tidyverse* packages, or in GraphPad Prism v8.4.3. Details of individual statistical models, including GLS model specifications, test choices, and multiple-comparison correction methods, are described in the relevant Methods subsections. Significance annotations on all figures follow the convention: \*\*\*\**P* < 0.0001, \*\*\**P* < 0.001, \*\**P* < 0.01, \**P* < 0.05, ns *P* > 0.05.

### Disclosure for the use of AI tools

Custom Python and R scripts used for the quantification of NanoBiT kinetics, integration of DEER probability density maps, and statistical analysis of NanoBiT assays and MD simulations were developed by the authors, with assistance from the large language model tools Perplexity (Perplexity AI), Grok (xAI), and Claude (Anthropic). All scripts were independently checked, modified, tested, and validated by the authors, who take full responsibility for the code and analyses. The authors used the AI tools Perplexity, Grok, and Claude for AI-assisted language copy editing to improve readability, grammar, and spelling. The authors reviewed and take full responsibility for all content.

## Acknowledgements

Electron microscopy data were collected at the Cryo-EM Facility of the University of Freiburg (RRID: SCR_025860). The Titan Krios G4 cryo-TEM used for imaging was funded by Deutsche Forschungsgemeinschaft (project no. 506518771) and is operated within the Microscopy and Image Analysis Platform (MIAP), University of Freiburg. The authors acknowledge support by the state of Baden-Württemberg through bwHPC, the German Research Foundation (DFG) through grant INST 40/575-1 FUGG (JUSTUS 2 high-performance computing cluster), and the staff of the University of Ulm computer center. The authors are grateful to Reinhild Rösler (ULMTeC Core Facility Mass Spectrometry & Proteomics of the Medical Faculty at Ulm University) for technical assistance and for drafting the mass spectrometry Materials and Methods section. The authors thank the MagRes Center of the University of Freiburg for the use of their EPR spectrometers. The authors are grateful to Ashley Redman for his help in recording some of the DEER time curves. Monique Furlan is acknowledged for helpful discussions on setting up the peroxisomal colocalization experiment. The authors thank Joscha Borho (Institute of Experimental and Clinical Pharmacology, Toxicology and Pharmacology of Natural Products, Ulm University) for technical support regarding the isothermal titration calorimetry (ITC) instrument.

## Author Contributions

B.G. conceptualized and designed the study, performed all experiments unless otherwise stated, formally analyzed the data, wrote the manuscript draft, and prepared the figures. B.J., S.S., and J.Re. performed cryo-EM. B.J. processed and formally analyzed the cryo-EM data, built the structural model, and prepared the corresponding supplementary figure. A.G. and J.S. performed pulsed EPR measurements. A.G., J.S., E.S., and B.G. formally analyzed the corresponding data. B.J. and A.G. drafted method-specific Materials and Methods subsections. J.Re. purified the protein complexes for cryo-EM and SEC-MALS experiments. J.F.G. and B.G. cloned and purified human septin constructs and performed the corresponding pulldown experiments. C.S. performed SEC-MALS measurements and formally analyzed the corresponding data. T.W. assisted with mass photometry measurements and formal analysis of the corresponding data. T.V. assisted with stopped-flow measurements and formal analysis of the corresponding data. A.G., J.S., and J.Ru. performed cw-EPR measurements. A.G., J.Ru., J.S., and E.S. formally analyzed the corresponding data. E.S. supervised pulsed EPR and cw-EPR experiments. S.G. supervised cryo-EM data processing and model building. N.J acquired funding. T.G. and N.J. provided overall supervision of the project, contributed to refining the study design, and to writing and editing of the manuscript. T.G. handled correspondence with the journal. All authors reviewed the manuscript and participated in data interpretation.

## Supplementary Figures

**Supplementary Fig. 1.**
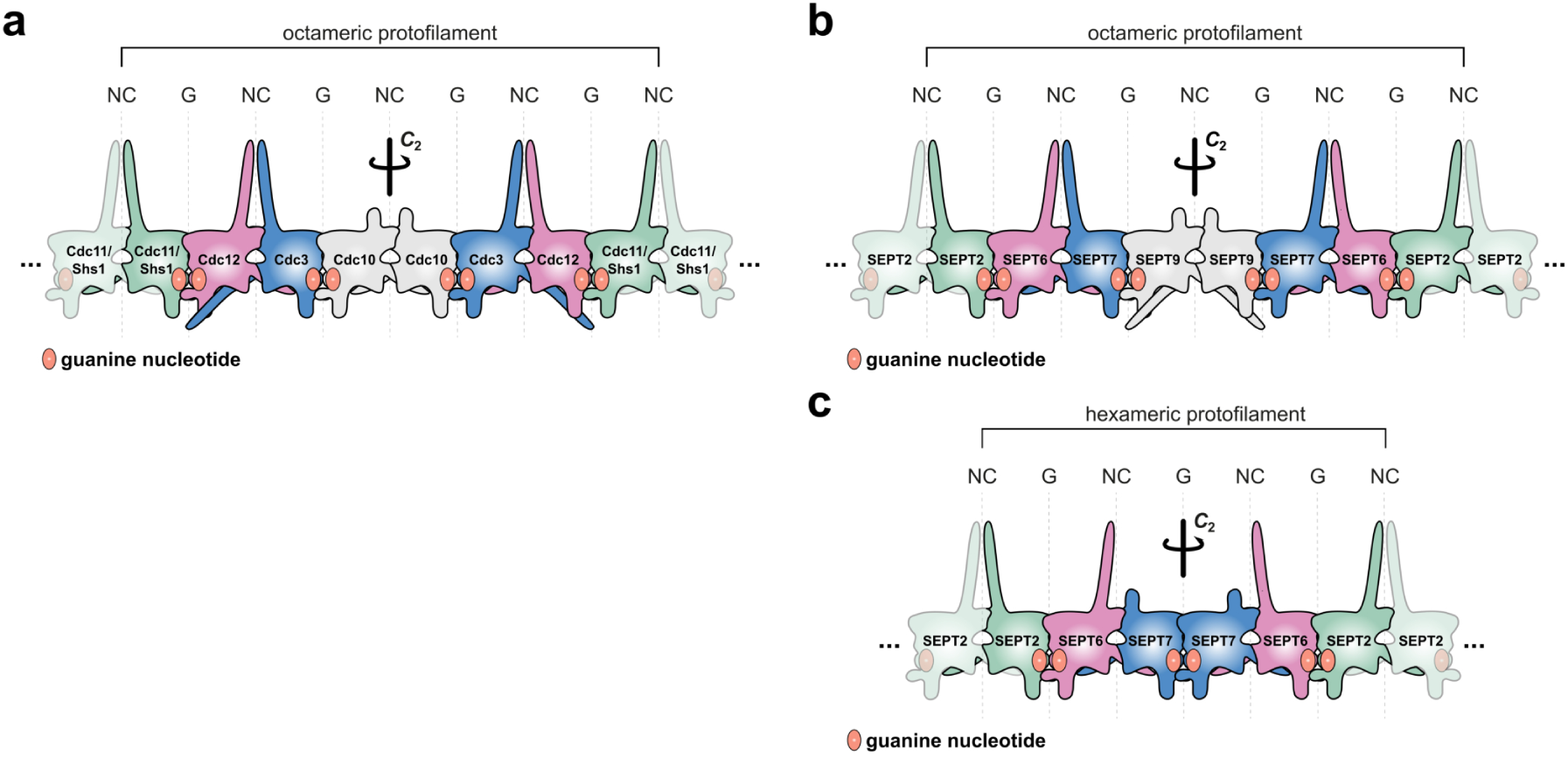
Septin protofilaments in budding yeast and mammals. **a**, Canonical octameric septin protofilaments in *S. cerevisiae*, in which the terminal position can be occupied by either Cdc11 or Shs1. **b**, Canonical mammalian octameric protofilament. **c**, Mammalian hexameric protofilament formed in the absence of SEPT9. Cartoons adapted from ^4^, © De Gruyter Brill, used with permission.

**Supplementary Fig. 2.**
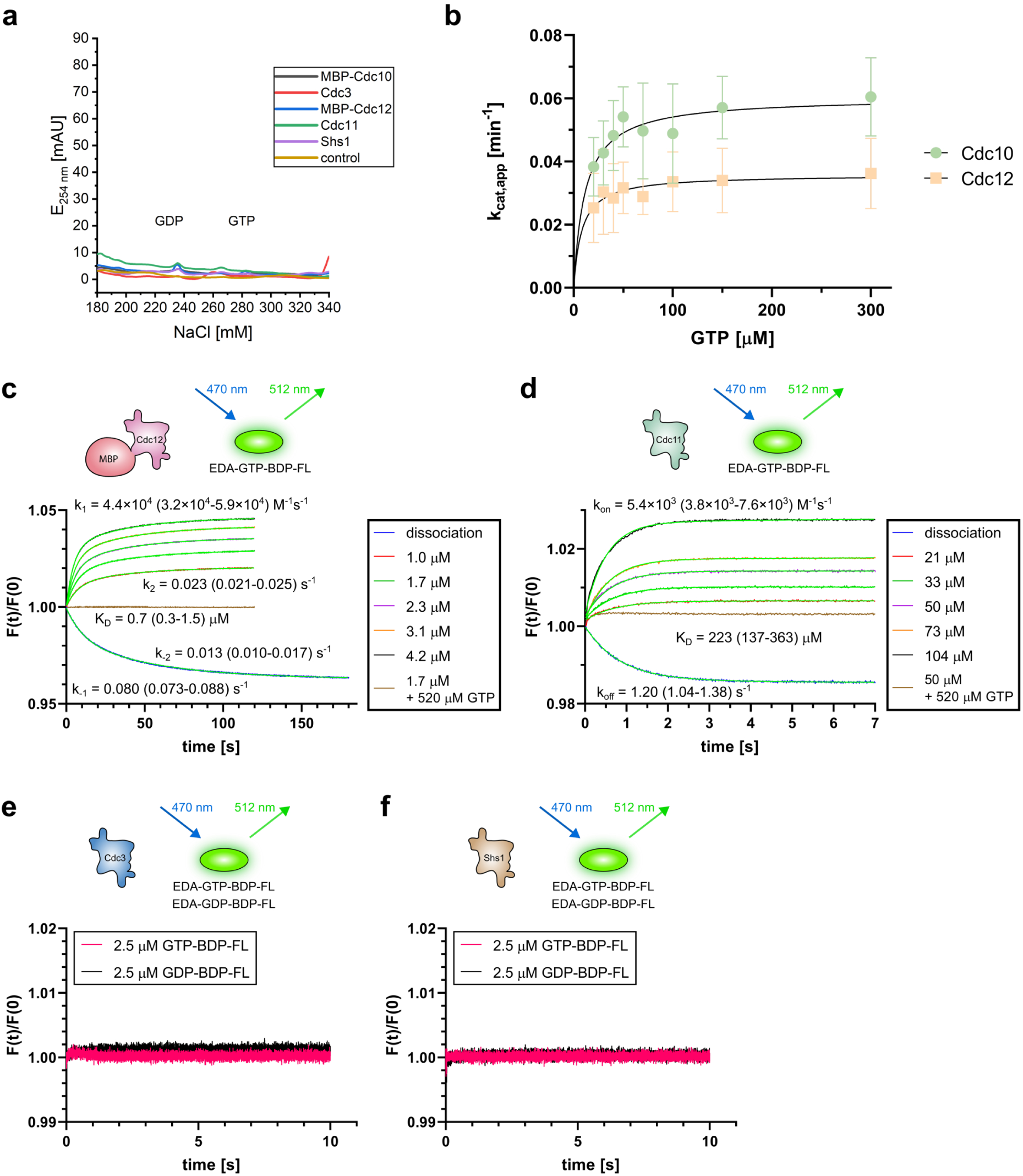
Nucleotide binding and hydrolysis by monomeric yeast septin G-domains. **a**, Nucleotide-content analysis of purified, heat-denatured monomeric G-domains after incubation with 3 mM GTP and GDP followed by desalting. No stably bound nucleotide was detected. A protein-free desalting control is shown for comparison. **b**, GTP hydrolysis by monomeric Cdc10 and Cdc12 quantified by malachite green assays over a range of initial GTP concentrations (20–300 µM). Data points show arithmetic means ± s.d. of three independent experiments, each averaged from 1–4 technical replicates. **c**, Representative stopped-flow fluorescence traces of EDA-GTP-BDP-FL association and dissociation with monomeric Cdc12. Global non-linear fits are shown in neon green. Kinetic parameters (*k1*, *k−1*, *k2*, *k−2*, *KD*) are geometric means with asymmetric 95% CIs from six independent association experiments (association was measured at 1.0–4.2 µM Cdc12 (four preparations) or 1.0–3.1 µM (two preparations)) and four independent dissociation experiments. **d**, As in **c** for Cdc11 with kinetic constants from four independent experiments. **e**,**f**, Stopped-flow fluorescence traces for Cdc3 (**e**) and Shs1 (**f**) revealing no detectable binding to EDA-GTP-BDP-FL or EDA-GDP-BDP-FL. However, we cannot exclude the possibility that these subunits are incompatible with the fluorescent nucleotide analogs.

**Supplementary Fig. 3.**
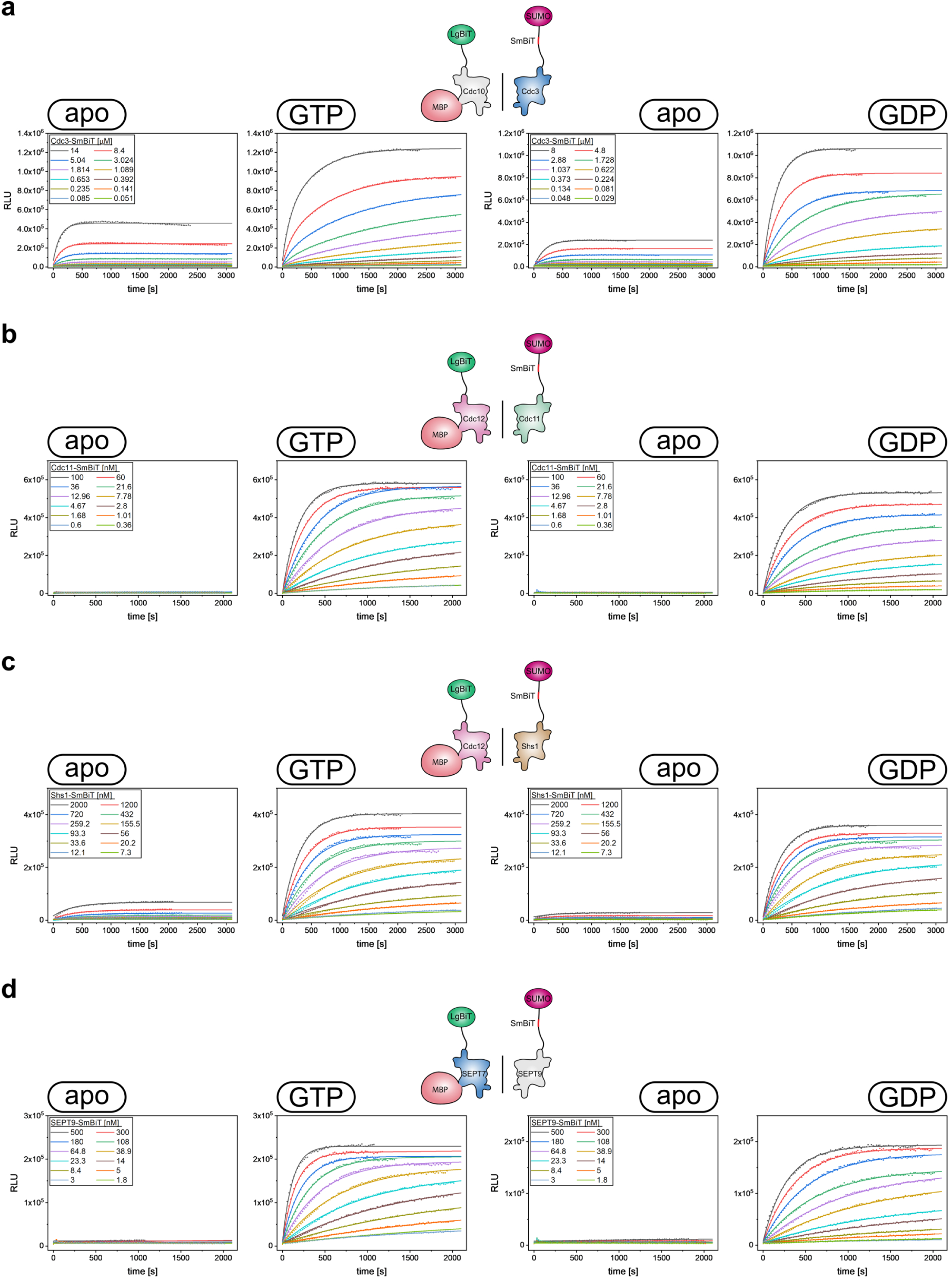
Global kinetic analysis of septin G-interface association measured by the NanoBiT assay. **a–d**, Shown are representative raw luminescence time courses (dots) and corresponding global fits (solid lines) for Cdc10-Cdc3 (**a**), Cdc12-Cdc11 (**b**), Cdc12-Shs1 (**c**), and SEPT9-SEPT7 (**d**) G-interface formation in the presence of 3 mM GTP or GDP. For each SmBiT concentration, apo traces (no nucleotide) and specific traces (GTP or GDP) were fitted jointly by an association model comprising explicit nonspecific and specific components, with shared kinetic parameters per concentration.

**Supplementary Fig. 4.**
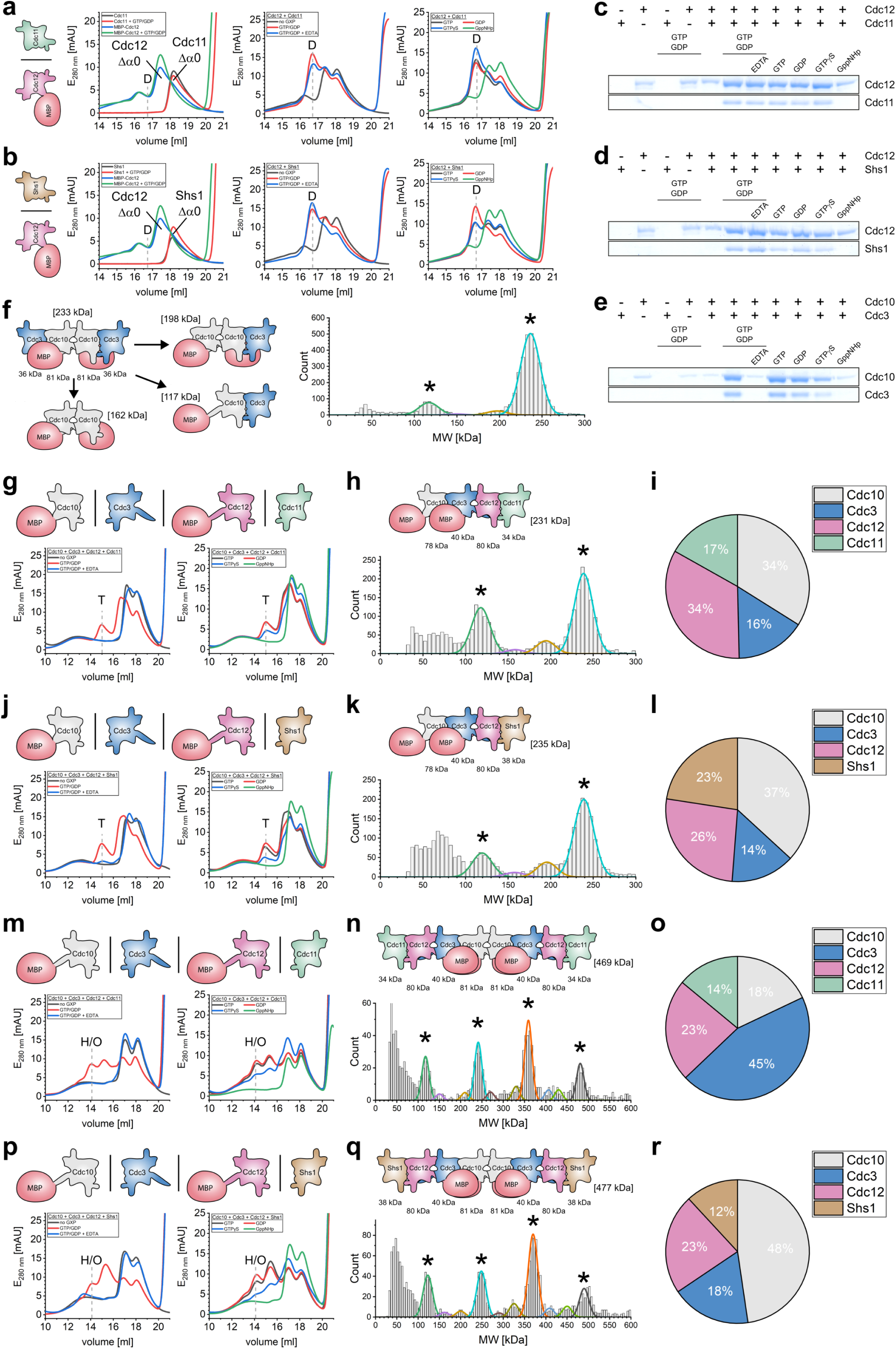
Incremental assembly of septin protofilaments *in vitro*. **a**,**b**, Analytical SEC of G-domain Cdc12 in the absence or presence of G-domain Cdc11 (**a**) or Shs1 (**b**) and the indicated nucleotides after overnight incubation. ‘D’ denotes the heterodimeric elution volume. **c–e**, SDS-PAGE of dimeric SEC fractions from **a**, **b**, and Fig. 3a, respectively. SEC runs mirror the nucleotide dependencies for G-interface formation observed in pull-down experiments (Fig. 2a–c) and confirm the dimeric character of the complexes. **f**, Mass photometry of tetrameric complexes formed by full-length Cdc10 and G-domain Cdc3 from Fig. 3a. Asterisks mark peaks attributed to fragments lacking exposed G-interfaces. **g**,**j**,**m**,**p**, Analytical SEC of higher-order assemblies reconstituted from G-domain (**g**,**j**) or full-length (**m**,**p**) Cdc10 together with Cdc3, Cdc12, and Cdc11 (**g**,**m**) or Shs1 (**j**,**p**). ‘T’ and ‘H/O’ indicate tetrameric and hexameric/octameric elution volumes as assigned by mass photometry. **h**,**k**,**n**,**q**, Mass photometry of complexes from **g**, **j**, **m**, and **p**, respectively. **i**,**l**,**o**,**r**, Mass spectrometric analysis of samples from **g**, **j**, **m**, and **p**, respectively, identifying all complex components in the purified fractions. Because band resolution and relative staining on SDS-PAGE do not allow unambiguous assignment of all subunits in these higher-order assemblies, mass spectrometry was used to verify the presence of Cdc10, Cdc3, Cdc12, and Cdc11 or Shs1 at comparable signal intensities, consistent with successful complex formation.

**Supplementary Fig. 5.**
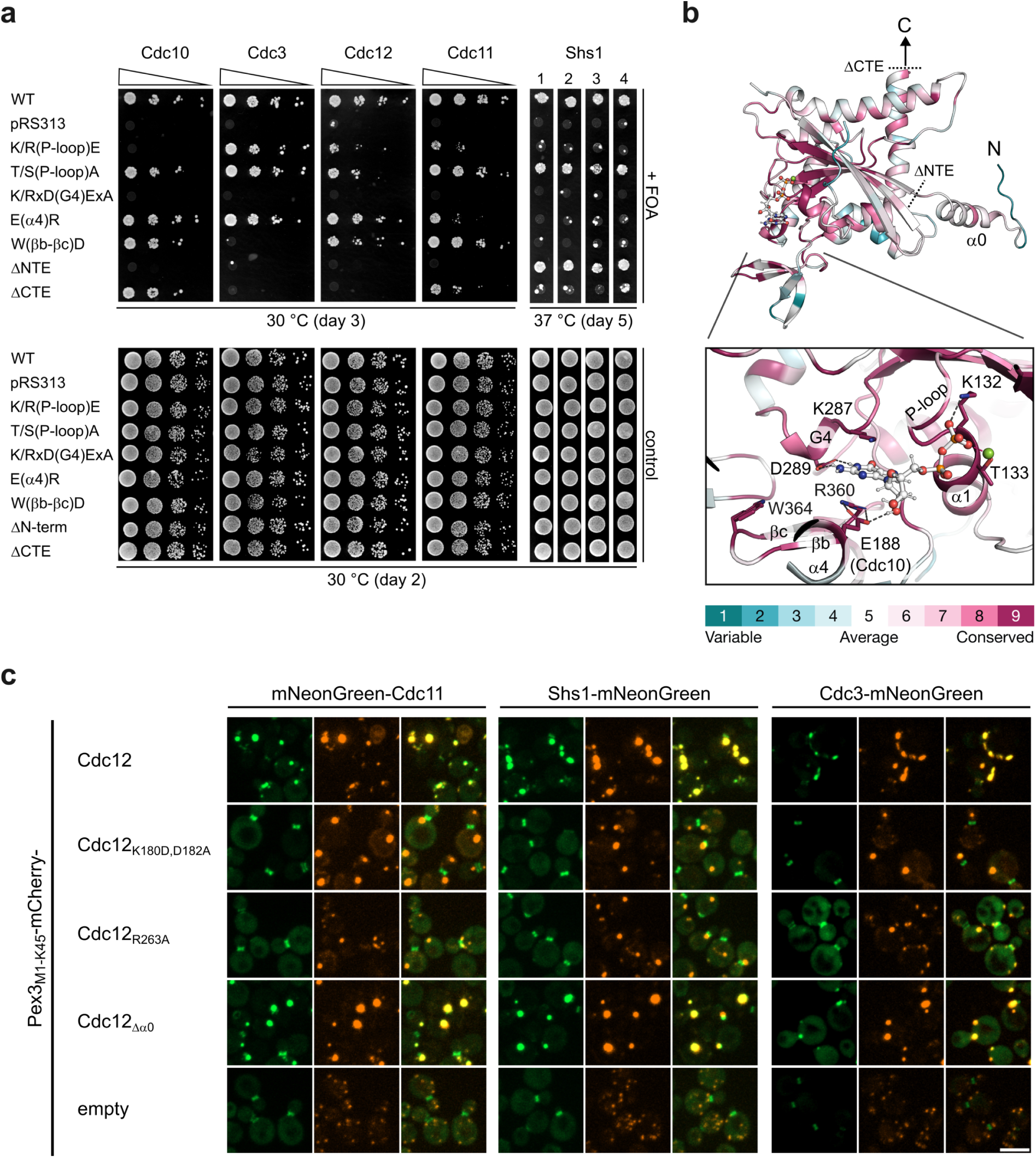
*In vivo* hierarchy of G- and NC-interface assembly. **a**, Viability assay screening G- and NC-interface mutations in yeast septins. Lethality indicates filament assembly defects^27,31,43^. Mutations were designed to impair nucleotide binding (K/R(P-loop)E, K/RxD(G4)ExA), magnesium coordination (T/S(P-loop)A), G-interface contacts (E(α4)R, W(βb-βc)D), or NC-interface integrity (ΔNTE, ΔCTE). A detailed mapping of the mutated sites to the amino acids in the different subunits is shown in Supplementary Table 5. The empty vector (pRS313) functioned as a negative control. Serial dilutions were spotted onto selective media and grown at 30 °C. *shs1Δ* is temperature-sensitive^27^ and was incubated at 37 °C without serial dilution; there four different clones are shown. **b**, Structure of Cdc3 (PDB 9GD4) showing the mutated residues and truncations in the context of the conserved septin GTPase fold and interfaces. The color-coded sequence conservation was calculated using Consurf^98^ as described previously^27^. **c**, Representative fluorescence images of the peroxisomal membrane-recruitment assay with Pex3M1-K45-mCherry fusions to Cdc12 and its mutants quantified in Fig. 3g. Scale bar, 5 µm.

**Supplementary Fig. 6.**
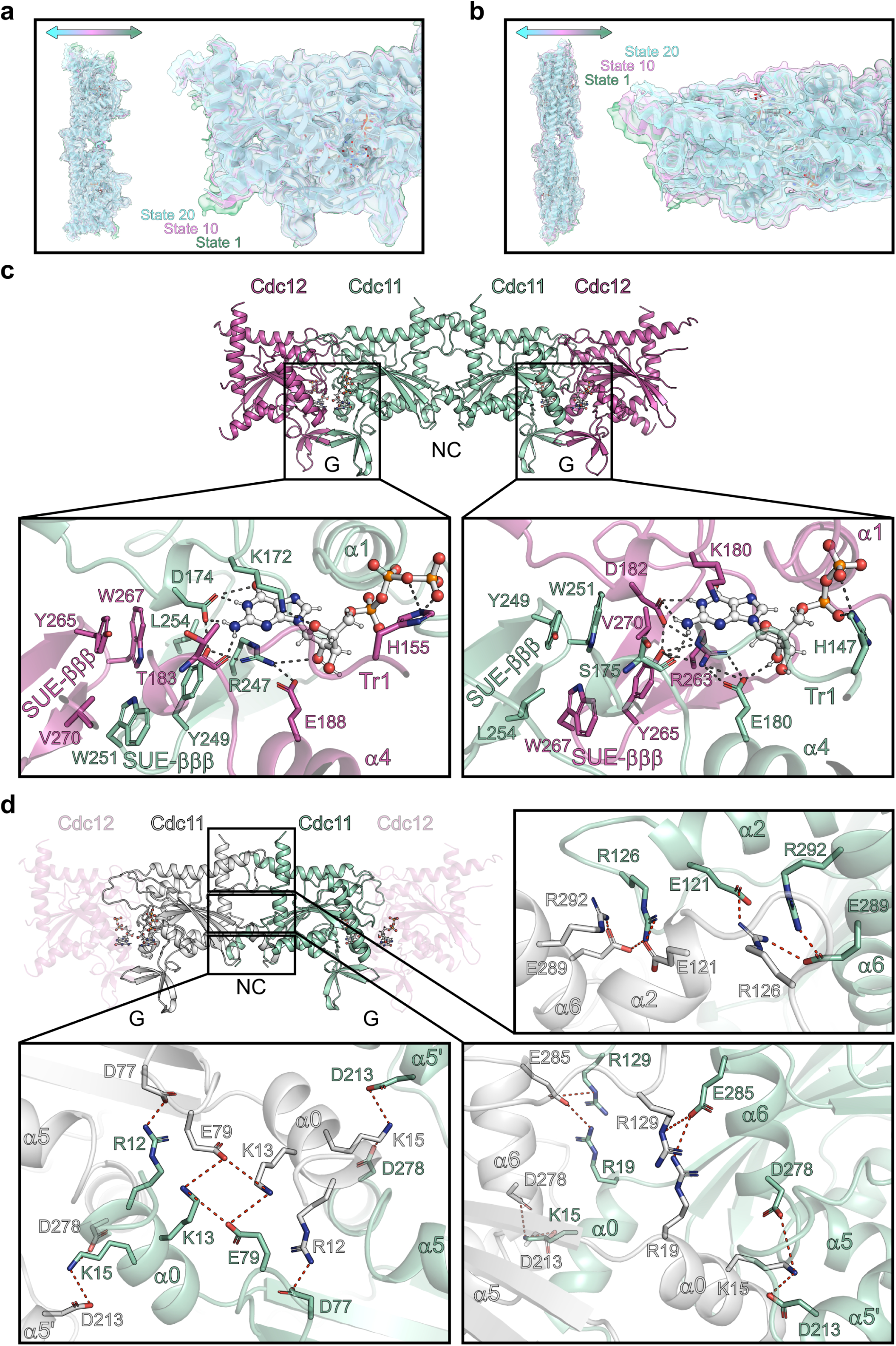
Conformational flexibility and interface architecture of the Cdc12-Cdc11-Cdc11-Cdc12 septin complex. **a**,**b**, Superposition of three representative states (states 1, 10 and 20) from a 20-state cryoSPARC 3D variability analysis, shown along the first (**a**) and second (**b**) principal eigenvectors, illustrating the range of conformational motion captured in the dataset. **c**, Details of the Cdc12-Cdc11 G-interface showing canonical contacts around the nucleotide binding pocket, including the Arg(βb) hydrogen bond network, His(Tr1)-mediated phosphate coordination of the adjacent nucleotide, and packing of hydrophobic residues of two adjacent SUE-βββ elements. Hydrogen bonds are shown as dashed lines in dark grey. **d**, The Cdc11-Cdc11 NC-interface is stabilized primarily through charged interactions (red dashes) at three interfacial sites. In the lower part of the interface, the α0-helices engage with each other (bottom, left), the central part α0-helices interact with the α5’- and α6-helices (bottom, right), and at the top, charged residues of the α2- and α6-helices form a conserved salt-bridge network similar to other septin complexes (top, right)^31,34,99^.

**Supplementary Fig. 7.**
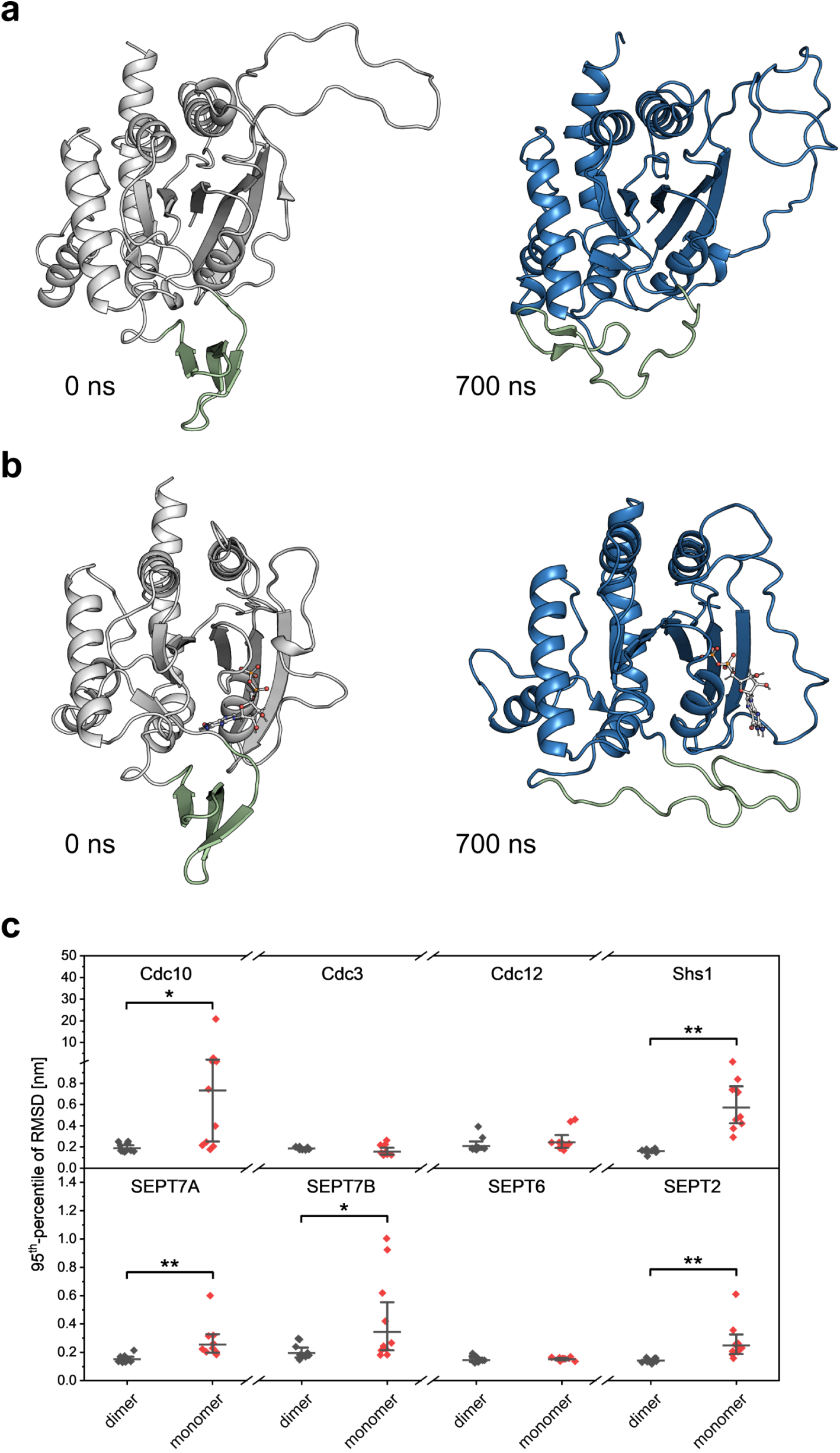
G-interface formation stabilizes the SUE-βββ and nucleotide binding in MD simulations. **a**, Trajectory of SUE-βββ unfolding in monomeric Cdc3:apo. Shown are the starting structure (white) and the conformation after 700 ns (blue). The SUE-βββ is highlighted in pale green. Extensive local unfolding and outward displacement of the SUE-βββ is observed, approaching a Cdc11-like unfolded state by the end of the trajectory. **b**, As in **a**, for monomeric GDP-bound SEPT7, showing major SUE-βββ unfolding accompanied by partial nucleotide dissociation from the active site. **c**, Stabilization of bound guanine nucleotides by G-interface dimerization in MD simulations, quantified by RMSD95 of nucleotide non-hydrogen atoms (700 ns, 10 replicas per condition). Points show RMSD95 from individual trajectories. Horizontal bars and whiskers indicate geometric means and asymmetric 95% CIs. Brackets denote Wilcoxon rank-sum contrasts (dimer vs. monomer) with Benjamini-Hochberg correction.

**Supplementary Fig. 8.**
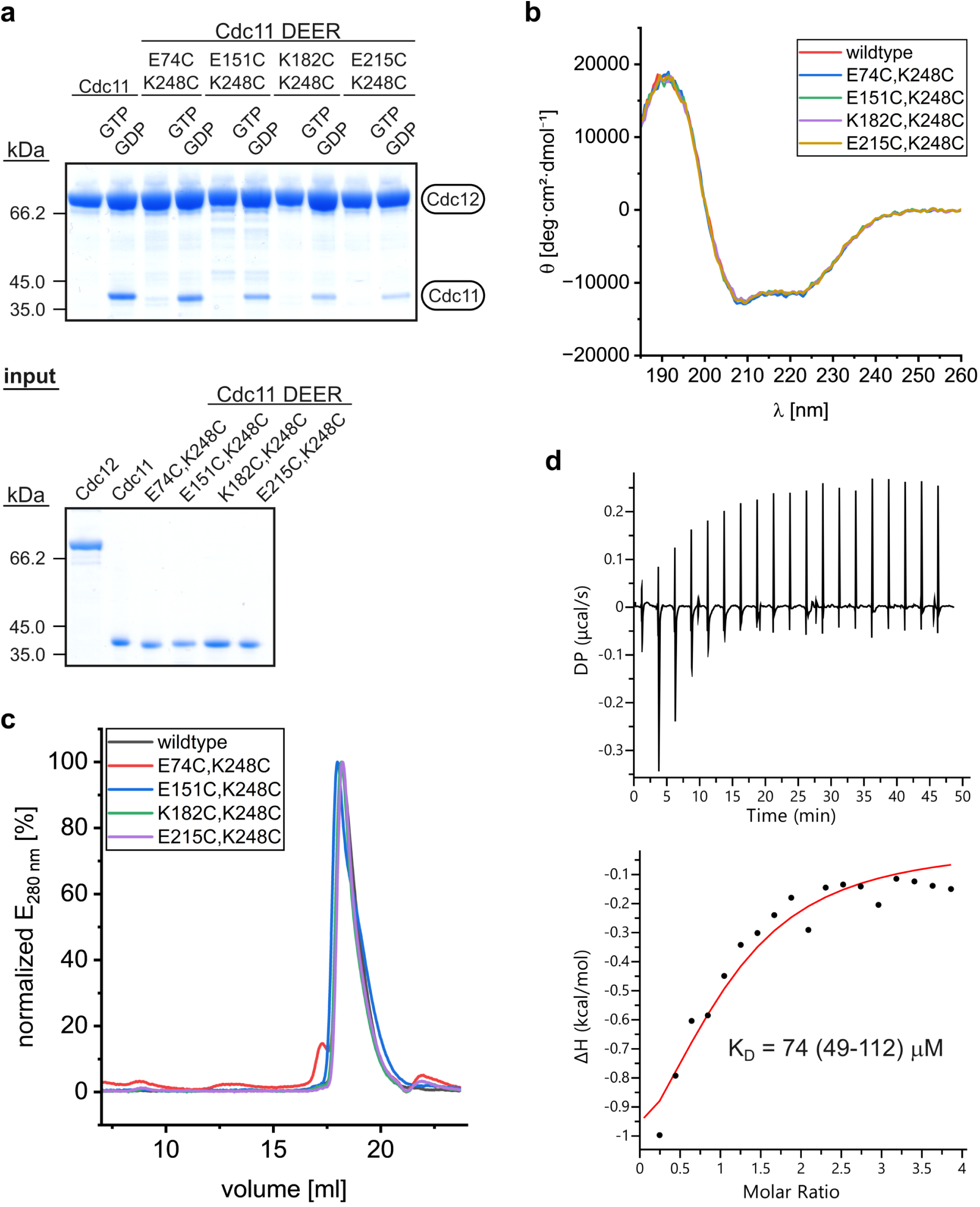
Validation of Cdc11 constructs for DEER spectroscopy. **a**, Pull-down assay testing nucleotide-dependent Cdc12-Cdc11 G-interface assembly. G-domain Cdc12 and G-domain Cdc11 carrying spin-label mutations were incubated with amylose resin in the absence or presence of an equimolar GTP/GDP mixture, and bound proteins were analyzed by SDS-PAGE. All Cdc11 constructs showed nucleotide-dependent co-precipitation with Cdc12, confirming that the engineered mutations preserve G-interface formation. **b**, Far-UV CD spectra of the Cdc11 DEER constructs, demonstrating native-like secondary structure in all variants. **c**, Analytical SEC chromatograms of spin-labeled Cdc11 constructs compared to the unlabeled wild-type, confirming that spin labeling does not perturb the monomeric state. **d**, Isothermal titration calorimetry of GTP binding to G-domain Cdc11. Representative thermogram (top) and integrated heats fitted to a single-site binding model (bottom). *KD* is the geometric mean of three independent protein preparations with asymmetric 95% CIs.

**Supplementary Fig. 9.**
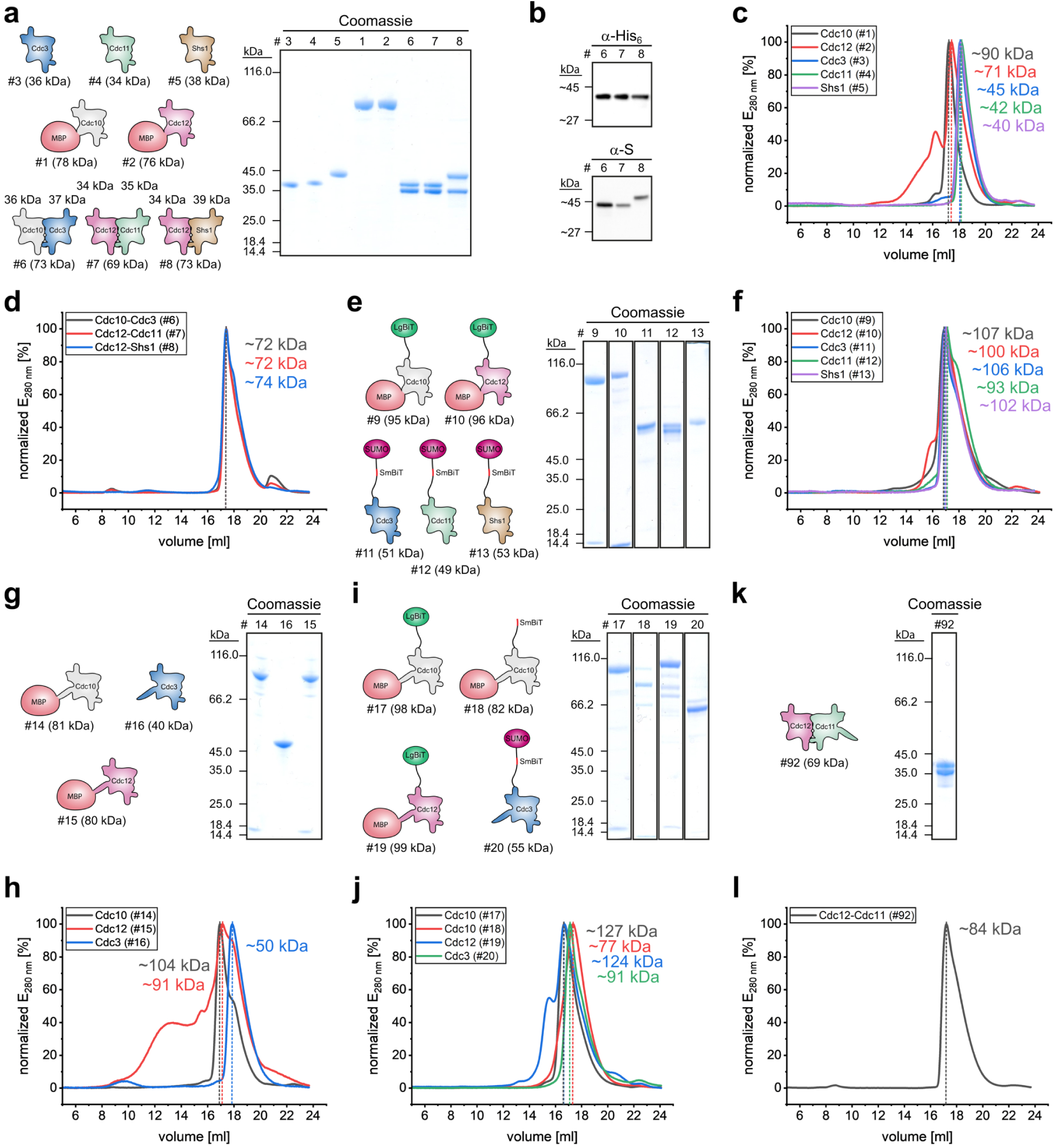
Biochemical characterization of purified yeast septin constructs. **a**,**e**,**g**,**i**,**k**, Schematic representations and SDS-PAGE analysis of purified septin constructs. Theoretical molecular weights are indicated. **b**, Western blot analysis using anti-His6 and anti-S tag antibodies, confirming identity and integrity of purified G-interface complexes. **c**,**d**,**f**,**h**,**j**,**l**, Analytical SEC of purified constructs. Estimated molecular weights are indicated. Co-expressed G-domains elute as dimers. Individual septins elute as monomers, and SUMO-tagged constructs elute at elevated apparent molecular weights, likely reflecting contributions of the SUMO tag.

**Supplementary Fig. 10.**
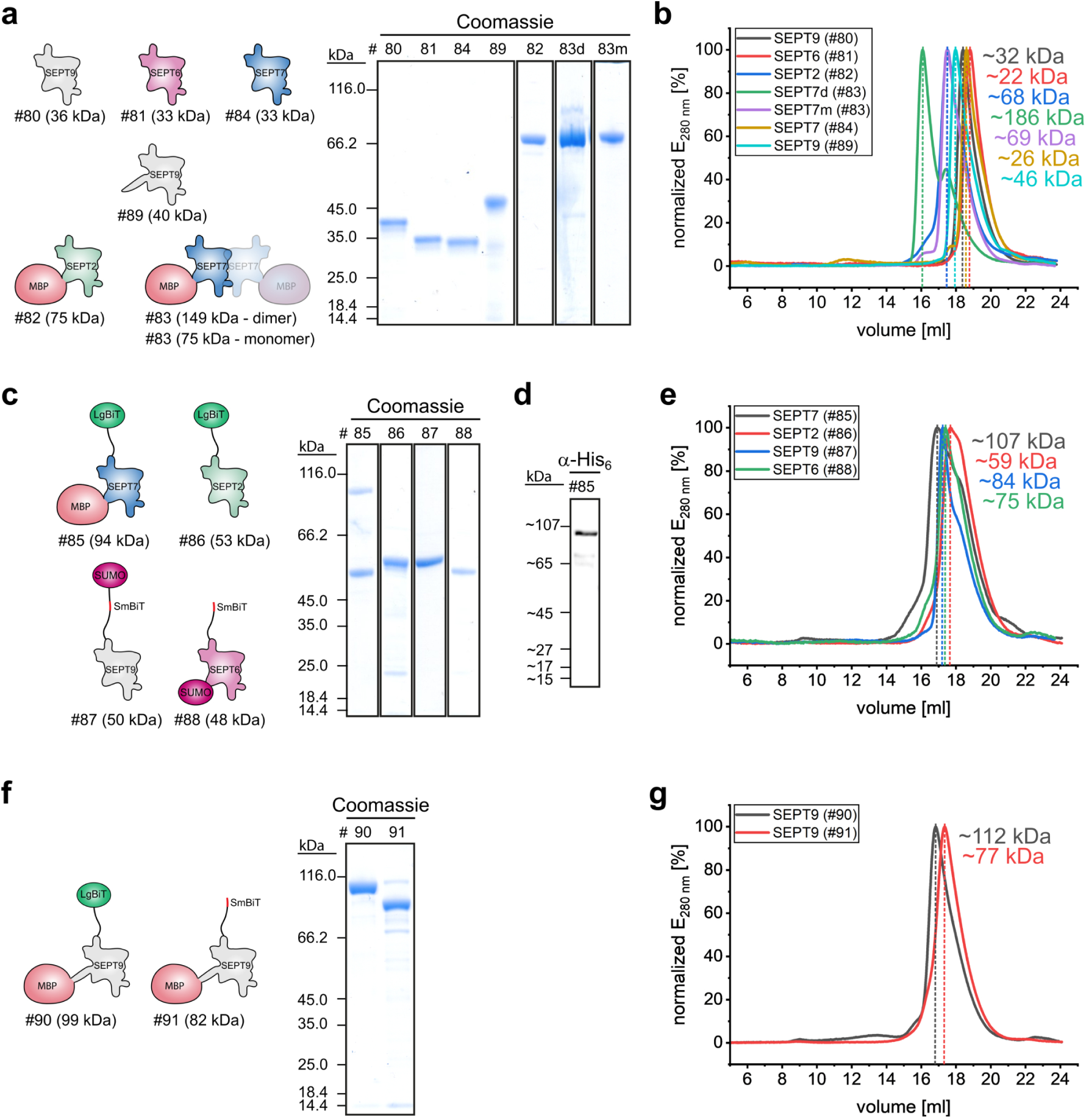
Biochemical characterization of purified human septin constructs. **a**,**c**,**f**, Schematic representations and SDS-PAGE analysis of purified septin constructs. Theoretical molecular weights are indicated. **b**,**e**,**g**, Analytical SEC of purified constructs. Estimated molecular weights are indicated. The elevated apparent molecular weights of SUMO-tagged constructs likely reflect contributions of the SUMO tag. In **a** and **b** #83d and #83m indicate dimeric fractions of SEPT7 and monomeric fractions obtained after treatment with 1 M MgCl2, respectively. **d**, Western blot analysis using anti-His6, confirming identity of construct #85.

**Supplementary Fig. 11.**
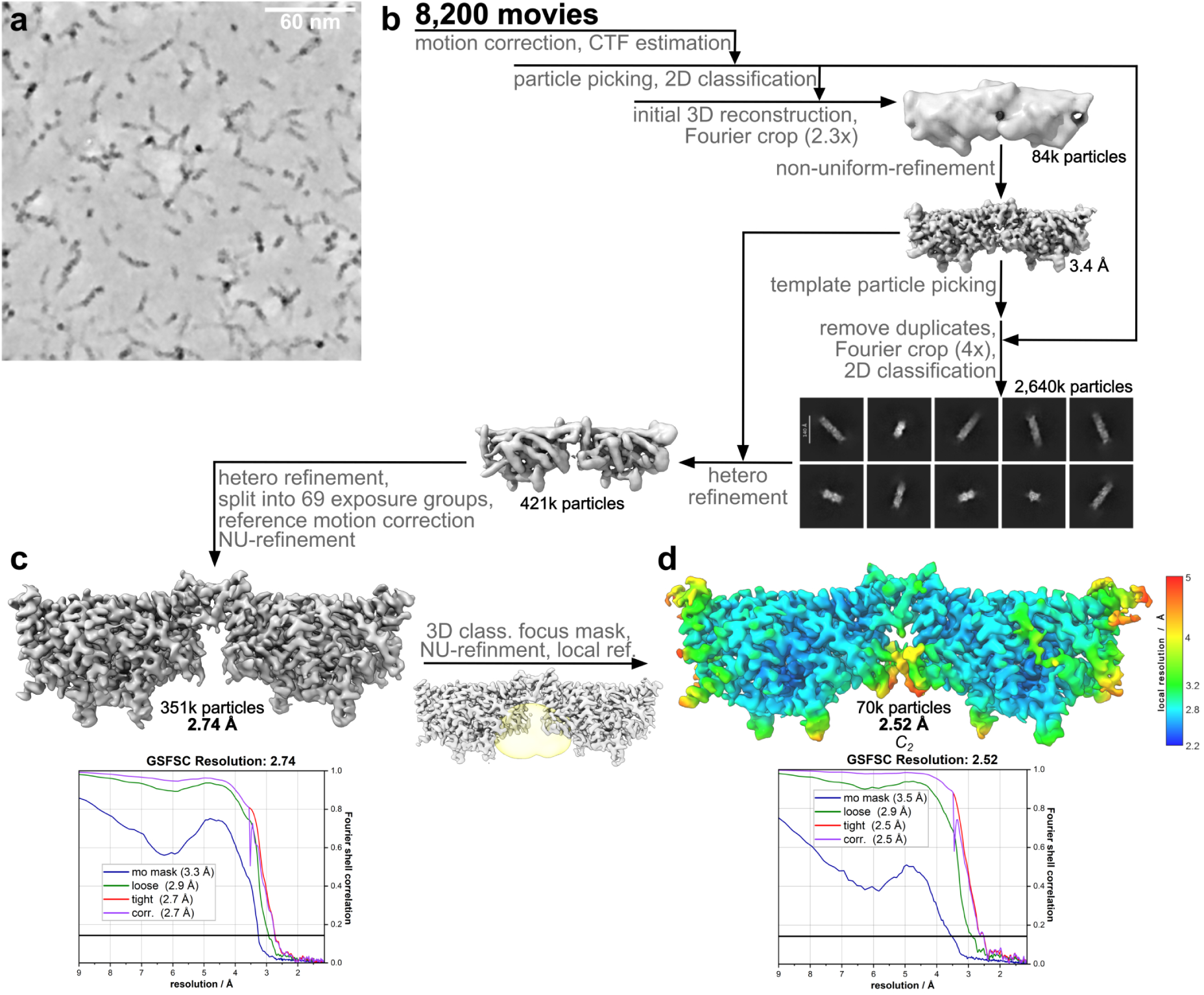
Cryo-EM analysis of the inside-out septin assembly Cdc12-Cdc11-Cdc11-Cdc12. **a**, Representative denoised micrograph from the dataset of 8,200 movies. **b**, Cryo-EM data-processing workflow showing iterative particle picking and refinement, yielding 421,937 particles. **c**, Initial reconstruction obtained from 351,177 particles without imposed symmetry, at 2.74 Å resolution. **d**, After 3D classification with a focus mask around the NC-interface, a *C*2-symmetric reconstruction was obtained from 70,904 particles at 2.52 Å resolution. Local-resolution coloring highlights the lower-resolution NC-interface. Gold-standard FSC (GSFSC) curves are shown below **c** and **d** (threshold, 0.143).

**Supplementary Fig. 12.**
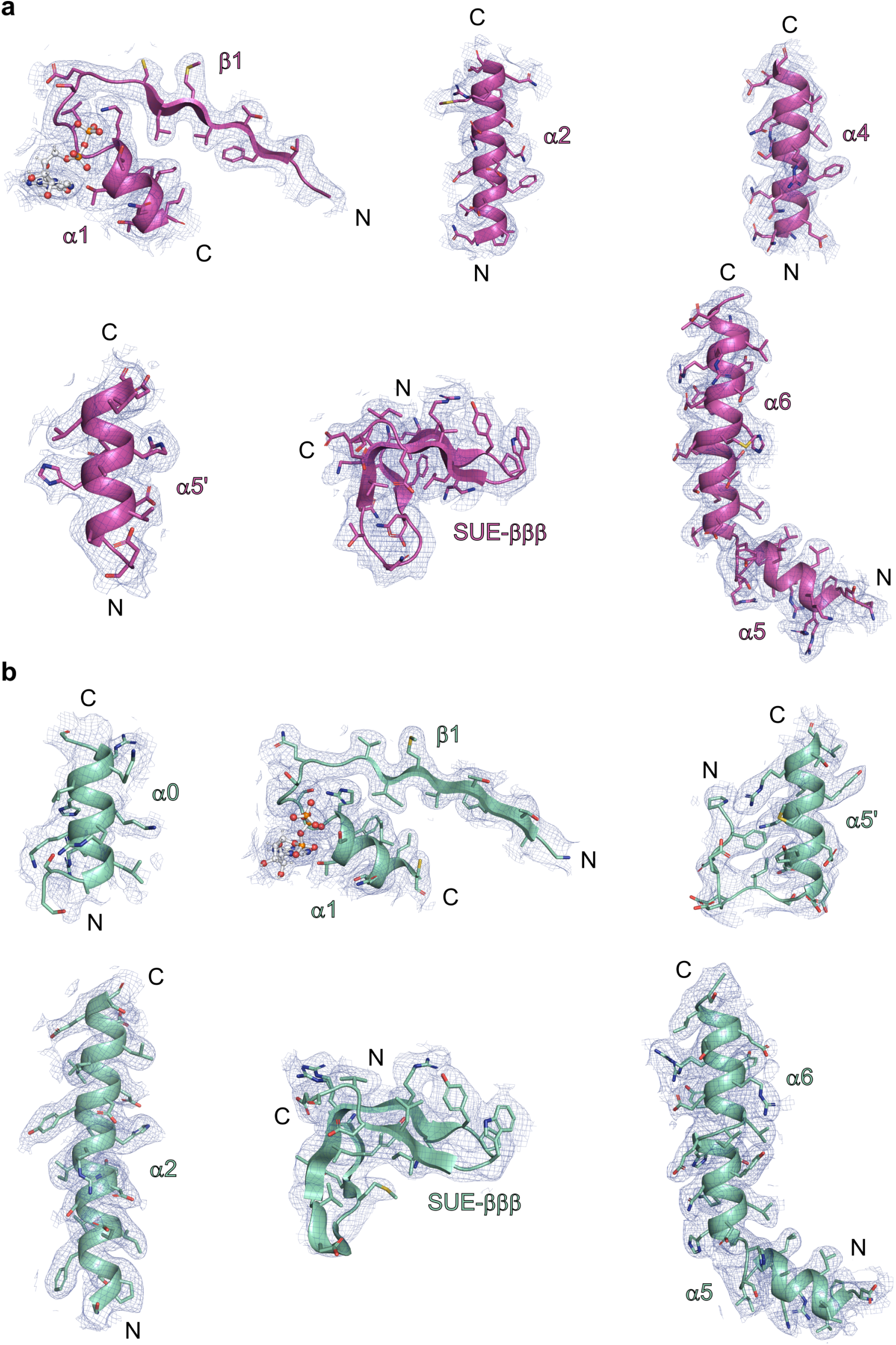
Cryo-EM map validation. The 2.52 Å resolution cryo-EM map is displayed as a blue mesh contoured at 6.0 σ, carved to 2.5 Å around displayed atoms from Cdc12 (**a**) and Cdc11 (**b**). Sigma values calculated from PyMOL’s internal map normalization (mean = 0, s.d. = 1).

**Supplementary Fig. 13.**
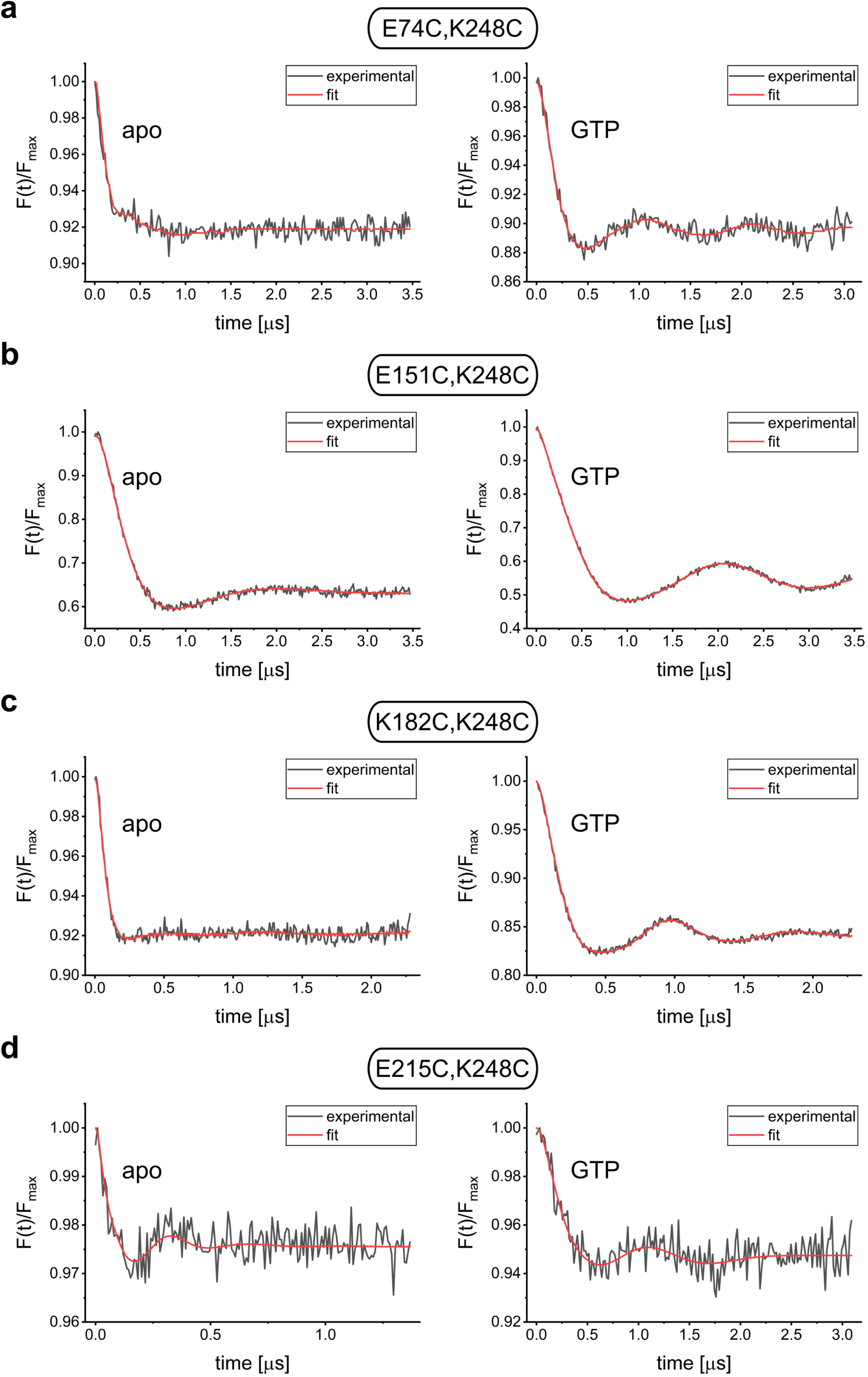
Background-corrected DEER form factors of Cdc11 variants. **a–d**, Experimental DEER form factors *F(t)/Fmax* (black) and corresponding fits (red) for spin-labeled Cdc11 variants carrying the indicated cysteine pairs (**a**, E74C,K248C; **b**, E151C,K248C; **c**, K182C,K248C; **d**, E215C,K248C) in the apo and GTP-bound states. Traces show background-divided DEER time-domain signals, normalized to the maximum of the background-corrected experimental form factor. For E74C,K248C, E151C,K248C, and K182C,K248C (**a–c**), fits and distance distributions were obtained using ComparativeDeerAnalyzer 2.0^91^ consensus analysis combining DEERNet and Tikhonov-based DeerLab solutions, whereas the E215C,K248C data (**d**) were analyzed by Tikhonov regularization using the L-curve corner criterion as implemented in DeerAnalysis2022^92^.

## Supplementary Tables

**Supplementary Table 1.** Pairwise comparisons of nucleotide effects on Cdc10-Cdc10 NC-interface NanoBiT luminescence in the presence or absence of Cdc3. Tukey-adjusted pairwise contrasts from the three-way GLS model (normalized RLU ∼ Cdc3 × buffer × nucleotide) are shown for conditions without (top) and with (bottom) the G-interface partner Cdc3. Estimates represent differences in estimated marginal means of normalized relative light units; SE, pooled standard error of the contrast; *df*, residual degrees of freedom; *t* ratio, test statistic; adjusted *P*, Tukey HSD-corrected *P*-value. Adjustments were applied separately within each buffer × Cdc3 combination. Positive estimates indicate a higher signal for the first term in the contrast. Both NC-interface formation above baseline (noGXP) and nucleotide-specific differences in signal magnitude are observed exclusively in the presence of Cdc3 and MgCl2, confirming that G-interface engagement is a prerequisite for nucleotide-dependent NC-interface assembly.

| Contrast | G | Buffer | Estimate | SE | df | t ratio | adjusted P |
| --- | --- | --- | --- | --- | --- | --- | --- |
| noGXP - GTP | - | MgCl<sub>2</sub> | 0.062 | 0.019 | 42.5 | 3.17 | 0.0314 |
| noGXP - GDP | - | MgCl<sub>2</sub> | 0.033 | 0.020 | 46.9 | 1.64 | 0.5782 |
| noGXP - (GTP/GDP) | - | MgCl<sub>2</sub> | 0.054 | 0.020 | 43.9 | 2.77 | 0.0824 |
| noGXP - GTPyS | - | MgCl<sub>2</sub> | 0.061 | 0.019 | 42.5 | 3.16 | 0.0321 |
| noGXP - GppNHp | - | MgCl<sub>2</sub> | 0.057 | 0.020 | 43.4 | 2.91 | 0.0594 |
| GTP - GDP | - | MgCl<sub>2</sub> | -0.028 | 0.019 | 33.8 | -1.53 | 0.6477 |
| GTP - (GTP/GDP) | - | MgCl<sub>2</sub> | -0.007 | 0.018 | 27.4 | -0.40 | 0.9985 |
| GTP - GTPyS | - | MgCl<sub>2</sub> | 0.000 | 0.018 | 25.3 | -0.01 | 1.0000 |
| GTP - GppNHp | - | MgCl<sub>2</sub> | -0.005 | 0.018 | 26.7 | -0.26 | 0.9998 |
| GDP - (GTP/GDP) | - | MgCl<sub>2</sub> | 0.021 | 0.019 | 35.9 | 1.13 | 0.8670 |
| GDP - GTPyS | - | MgCl<sub>2</sub> | 0.028 | 0.019 | 33.8 | 1.52 | 0.6527 |
| GDP - GppNHp | - | MgCl<sub>2</sub> | 0.024 | 0.019 | 35.1 | 1.27 | 0.7996 |
| (GTP/GDP) - GTPyS | - | MgCl<sub>2</sub> | 0.007 | 0.018 | 27.5 | 0.40 | 0.9986 |
| (GTP/GDP) - GppNHp | - | MgCl<sub>2</sub> | 0.003 | 0.018 | 28.8 | 0.14 | 1.0000 |
| GTPyS - GppNHp | - | MgCl<sub>2</sub> | -0.005 | 0.018 | 26.7 | -0.25 | 0.9998 |
| noGXP - GTP | - | EDTA | 0.004 | 0.020 | 42.6 | 0.18 | 1.0000 |
| noGXP - GDP | - | EDTA | 0.011 | 0.019 | 41.0 | 0.55 | 0.9934 |
| noGXP - (GTP/GDP) | - | EDTA | 0.009 | 0.019 | 41.3 | 0.47 | 0.9969 |
| noGXP - GTPyS | - | EDTA | 0.007 | 0.019 | 41.8 | 0.36 | 0.9992 |
| noGXP - GppNHp | - | EDTA | 0.003 | 0.020 | 42.8 | 0.14 | 1.0000 |
| GTP - GDP | - | EDTA | 0.007 | 0.019 | 40.1 | 0.37 | 0.9990 |
| GTP - (GTP/GDP) | - | EDTA | 0.006 | 0.019 | 40.5 | 0.29 | 0.9997 |
| GTP - GTPyS | - | EDTA | 0.003 | 0.019 | 41.0 | 0.18 | 1.0000 |
| GTP - GppNHp | - | EDTA | -0.001 | 0.020 | 42.0 | -0.04 | 1.0000 |
| GDP - (GTP/GDP) | - | EDTA | -0.002 | 0.019 | 38.7 | -0.08 | 1.0000 |
| GDP - GTPyS | - | EDTA | -0.004 | 0.019 | 39.2 | -0.19 | 1.0000 |
| GDP - GppNHp | - | EDTA | -0.008 | 0.019 | 40.3 | -0.42 | 0.9982 |
| (GTP/GDP) - GTPyS | - | EDTA | -0.002 | 0.019 | 39.7 | -0.11 | 1.0000 |
| (GTP/GDP) - GppNHp | - | EDTA | -0.006 | 0.019 | 40.7 | -0.33 | 0.9994 |
| GTPyS - GppNHp | - | EDTA | -0.004 | 0.019 | 41.2 | -0.22 | 0.9999 |
| noGXP - GTP | Cdc3 | MgCl<sub>2</sub> | -0.559 | 0.027 | 11.53 | -20.45 | 0.0000 |
| noGXP - GDP | Cdc3 | MgCl<sub>2</sub> | -0.871 | 0.029 | 7.63 | -29.61 | 0.0000 |
| noGXP - (GTP/GDP) | Cdc3 | MgCl<sub>2</sub> | -0.873 | 0.029 | 7.62 | -29.67 | 0.0000 |
| noGXP - GTPyS | Cdc3 | MgCl<sub>2</sub> | -0.048 | 0.022 | 46.26 | -2.23 | 0.2463 |
| noGXP - GppNHp | Cdc3 | MgCl<sub>2</sub> | 0.035 | 0.020 | 43.11 | 1.80 | 0.4734 |
| GTP - GDP | Cdc3 | MgCl<sub>2</sub> | -0.312 | 0.034 | 5.36 | -9.05 | 0.0014 |
| GTP - (GTP/GDP) | Cdc3 | MgCl<sub>2</sub> | -0.314 | 0.035 | 5.36 | -9.10 | 0.0014 |
| GTP - GTPyS | Cdc3 | MgCl<sub>2</sub> | 0.511 | 0.028 | 10.91 | 18.18 | 0.0000 |
| GTP - GppNHp | Cdc3 | MgCl<sub>2</sub> | 0.594 | 0.027 | 11.88 | 22.32 | 0.0000 |
| GDP - (GTP/GDP) | Cdc3 | MgCl<sub>2</sub> | -0.002 | 0.036 | 4.60 | -0.05 | 1.0000 |
| GDP - GTPyS | Cdc3 | MgCl<sub>2</sub> | 0.823 | 0.030 | 7.46 | 27.31 | 0.0000 |
| GDP - GppNHp | Cdc3 | MgCl<sub>2</sub> | 0.906 | 0.029 | 7.67 | 31.51 | 0.0000 |
| (GTP/GDP) - GTPyS | Cdc3 | MgCl<sub>2</sub> | 0.825 | 0.030 | 7.44 | 27.36 | 0.0000 |
| (GTP/GDP) - GppNHp | Cdc3 | MgCl<sub>2</sub> | 0.908 | 0.029 | 7.65 | 31.56 | 0.0000 |
| GTPyS - GppNHp | Cdc3 | MgCl<sub>2</sub> | 0.083 | 0.021 | 48.00 | 4.03 | 0.0026 |
| noGXP - GTP | Cdc3 | EDTA | 0.000 | 0.019 | 40.34 | 0.03 | 1.0000 |
| noGXP - GDP | Cdc3 | EDTA | -0.031 | 0.020 | 46.23 | -1.55 | 0.6341 |
| noGXP - (GTP/GDP) | Cdc3 | EDTA | -0.004 | 0.019 | 41.34 | -0.19 | 1.0000 |
| noGXP - GTPyS | Cdc3 | EDTA | -0.001 | 0.019 | 40.60 | -0.03 | 1.0000 |
| noGXP - GppNHp | Cdc3 | EDTA | -0.007 | 0.020 | 42.09 | -0.36 | 0.9991 |
| GTP - GDP | Cdc3 | EDTA | -0.032 | 0.020 | 46.17 | -1.58 | 0.6181 |
| GTP - (GTP/GDP) | Cdc3 | EDTA | -0.004 | 0.019 | 41.23 | -0.22 | 0.9999 |
| GTP - GTPyS | Cdc3 | EDTA | -0.001 | 0.019 | 40.48 | -0.06 | 1.0000 |
| GTP - GppNHp | Cdc3 | EDTA | -0.008 | 0.020 | 41.98 | -0.39 | 0.9988 |
| GDP - (GTP/GDP) | Cdc3 | EDTA | 0.027 | 0.020 | 46.65 | 1.36 | 0.7518 |
| GDP - GTPyS | Cdc3 | EDTA | 0.031 | 0.020 | 46.30 | 1.52 | 0.6536 |
| GDP - GppNHp | Cdc3 | EDTA | 0.024 | 0.020 | 46.98 | 1.19 | 0.8411 |
| (GTP/GDP) - GTPyS | Cdc3 | EDTA | 0.003 | 0.019 | 41.48 | 0.16 | 1.0000 |
| (GTP/GDP) - GppNHp | Cdc3 | EDTA | -0.003 | 0.020 | 42.88 | -0.17 | 1.0000 |
| GTPyS - GppNHp | Cdc3 | EDTA | -0.007 | 0.020 | 42.22 | -0.33 | 0.9994 |

**Supplementary Table 2.** Pairwise comparisons of nucleotide effects on Cdc3-Cdc12 NC-interface NanoBiT luminescence in the presence or absence of G-interface partners. Tukey-adjusted pairwise contrasts from the three-way GLS model (normalized RLU ∼ G-interface partner × buffer × nucleotide) are shown for conditions without (top) and with (bottom) the G-interface partners Cdc10 and Cdc11 or Cdc10 and Shs1. Estimates represent differences in estimated marginal means of normalized relative light units; SE, pooled standard error of the contrast; *df*, residual degrees of freedom; *t* ratio, test statistic; adjusted *P*, Tukey HSD-corrected *P*-value. Adjustments were applied separately within each buffer × G-interface partner combination. Positive estimates indicate a higher signal for the first term in the contrast. Both NC-interface formation above baseline (noGXP) and nucleotide-specific differences in signal magnitude are observed exclusively in the presence of G-interface partners, confirming that G-interface engagement is a prerequisite for nucleotide-dependent NC-interface assembly. GDP and GTP/GDP retain partial activity under EDTA, suggesting residual Cdc10-Cdc3 G-interface formation can occur in the absence of Mg^2+^.

| Contrast | G | Buffer | Estimate | SE | df | t ratio | adjusted P |
| --- | --- | --- | --- | --- | --- | --- | --- |
| noGXP - GTP | - | MgCl<sub>2</sub> | 0.007 | 0.038 | 74.8 | 0.17 | 1.0000 |
| noGXP - GDP | - | MgCl<sub>2</sub> | 0.002 | 0.039 | 76.9 | 0.06 | 1.0000 |
| noGXP - (GTP/GDP) | - | MgCl<sub>2</sub> | 0.007 | 0.038 | 74.7 | 0.18 | 1.0000 |
| noGXP - GTPyS | - | MgCl<sub>2</sub> | 0.008 | 0.038 | 74.3 | 0.20 | 1.0000 |
| noGXP - GppNHp | - | MgCl<sub>2</sub> | 0.003 | 0.039 | 76.7 | 0.07 | 1.0000 |
| GTP - GDP | - | MgCl<sub>2</sub> | -0.004 | 0.038 | 73.6 | -0.11 | 1.0000 |
| GTP - (GTP/GDP) | - | MgCl<sub>2</sub> | 0.000 | 0.037 | 71.3 | 0.00 | 1.0000 |
| GTP - GTPyS | - | MgCl<sub>2</sub> | 0.001 | 0.037 | 70.9 | 0.02 | 1.0000 |
| GTP - GppNHp | - | MgCl<sub>2</sub> | -0.004 | 0.038 | 73.4 | -0.11 | 1.0000 |
| GDP - (GTP/GDP) | - | MgCl<sub>2</sub> | 0.004 | 0.038 | 73.5 | 0.12 | 1.0000 |
| GDP - GTPyS | - | MgCl<sub>2</sub> | 0.005 | 0.038 | 73.1 | 0.14 | 1.0000 |
| GDP - GppNHp | - | MgCl<sub>2</sub> | 0.000 | 0.038 | 75.6 | 0.01 | 1.0000 |
| (GTP/GDP) - GTPyS | - | MgCl<sub>2</sub> | 0.001 | 0.037 | 70.8 | 0.02 | 1.0000 |
| (GTP/GDP) - GppNHp | - | MgCl<sub>2</sub> | -0.004 | 0.038 | 73.3 | -0.11 | 1.0000 |
| GTPyS - GppNHp | - | MgCl<sub>2</sub> | -0.005 | 0.038 | 72.9 | -0.13 | 1.0000 |
| noGXP - GTP | - | EDTA | 0.000 | 0.035 | 60.2 | -0.01 | 1.0000 |
| noGXP - GDP | - | EDTA | -0.001 | 0.035 | 60.4 | -0.02 | 1.0000 |
| noGXP - (GTP/GDP) | - | EDTA | 0.000 | 0.035 | 60.0 | 0.00 | 1.0000 |
| noGXP - GTPyS | - | EDTA | 0.000 | 0.035 | 60.2 | -0.01 | 1.0000 |
| noGXP - GppNHp | - | EDTA | -0.007 | 0.036 | 64.1 | -0.21 | 0.9999 |
| GTP - GDP | - | EDTA | 0.000 | 0.035 | 60.5 | -0.01 | 1.0000 |
| GTP - (GTP/GDP) | - | EDTA | 0.000 | 0.035 | 60.2 | 0.01 | 1.0000 |
| GTP - GTPyS | - | EDTA | 0.000 | 0.035 | 60.3 | 0.00 | 1.0000 |
| GTP - GppNHp | - | EDTA | -0.007 | 0.036 | 64.3 | -0.20 | 1.0000 |
| GDP - (GTP/GDP) | - | EDTA | 0.001 | 0.035 | 60.4 | 0.02 | 1.0000 |
| GDP - GTPyS | - | EDTA | 0.000 | 0.035 | 60.5 | 0.01 | 1.0000 |
| GDP - GppNHp | - | EDTA | -0.007 | 0.036 | 64.5 | -0.19 | 1.0000 |
| (GTP/GDP) - GTPyS | - | EDTA | 0.000 | 0.035 | 60.1 | -0.01 | 1.0000 |
| (GTP/GDP) - GppNHp | - | EDTA | -0.007 | 0.036 | 64.1 | -0.21 | 0.9999 |
| GTPyS - GppNHp | - | EDTA | -0.007 | 0.036 | 64.3 | -0.20 | 1.0000 |
| noGXP - GTP | Cdc10 & Cdc11 | MgCl<sub>2</sub> | -0.798 | 0.071 | 34.6 | -11.22 | 0.0000 |
| noGXP - GDP | Cdc10 & Cdc11 | MgCl<sub>2</sub> | -0.617 | 0.066 | 43.6 | -9.28 | 0.0000 |
| noGXP - (GTP/GDP) | Cdc10 & Cdc11 | MgCl<sub>2</sub> | -0.775 | 0.071 | 35.5 | -10.98 | 0.0000 |
| noGXP - GTPyS | Cdc10 & Cdc11 | MgCl<sub>2</sub> | -0.677 | 0.068 | 40.1 | -9.94 | 0.0000 |
| noGXP - GppNHp | Cdc10 & Cdc11 | MgCl<sub>2</sub> | 0.005 | 0.039 | 77.1 | 0.12 | 1.0000 |
| GTP - GDP | Cdc10 & Cdc11 | MgCl<sub>2</sub> | 0.181 | 0.089 | 27.3 | 2.03 | 0.3514 |
| GTP - (GTP/GDP) | Cdc10 & Cdc11 | MgCl<sub>2</sub> | 0.023 | 0.092 | 25.3 | 0.25 | 0.9998 |
| GTP - GTPyS | Cdc10 & Cdc11 | MgCl<sub>2</sub> | 0.121 | 0.090 | 26.5 | 1.34 | 0.7584 |
| GTP - GppNHp | Cdc10 & Cdc11 | MgCl<sub>2</sub> | 0.803 | 0.071 | 34.5 | 11.32 | 0.0000 |
| GDP - (GTP/GDP) | Cdc10 & Cdc11 | MgCl<sub>2</sub> | -0.158 | 0.089 | 27.7 | -1.78 | 0.4939 |
| GDP - GTPyS | Cdc10 & Cdc11 | MgCl<sub>2</sub> | -0.060 | 0.087 | 29.4 | -0.69 | 0.9819 |
| GDP - GppNHp | Cdc10 & Cdc11 | MgCl<sub>2</sub> | 0.622 | 0.066 | 43.4 | 9.39 | 0.0000 |
| (GTP/GDP) - GTPyS | Cdc10 & Cdc11 | MgCl<sub>2</sub> | 0.098 | 0.090 | 26.9 | 1.09 | 0.8799 |
| (GTP/GDP) - GppNHp | Cdc10 & Cdc11 | MgCl<sub>2</sub> | 0.779 | 0.070 | 35.4 | 11.08 | 0.0000 |
| GTPyS - GppNHp | Cdc10 & Cdc11 | MgCl<sub>2</sub> | 0.681 | 0.068 | 40.0 | 10.04 | 0.0000 |
| noGXP - GTP | Cdc10 & Cdc11 | EDTA | -0.002 | 0.036 | 64.1 | -0.06 | 1.0000 |
| noGXP - GDP | Cdc10 & Cdc11 | EDTA | -0.345 | 0.056 | 73.8 | -6.14 | 0.0000 |
| noGXP - (GTP/GDP) | Cdc10 & Cdc11 | EDTA | -0.207 | 0.050 | 100.2 | -4.13 | 0.0011 |
| noGXP - GTPyS | Cdc10 & Cdc11 | EDTA | -0.019 | 0.038 | 73.1 | -0.49 | 0.9962 |
| noGXP - GppNHp | Cdc10 & Cdc11 | EDTA | 0.004 | 0.035 | 60.9 | 0.11 | 1.0000 |
| GTP - GDP | Cdc10 & Cdc11 | EDTA | -0.343 | 0.056 | 73.8 | -6.09 | 0.0000 |
| GTP - (GTP/GDP) | Cdc10 & Cdc11 | EDTA | -0.205 | 0.050 | 100.1 | -4.07 | 0.0013 |
| GTP - GTPyS | Cdc10 & Cdc11 | EDTA | -0.017 | 0.038 | 74.1 | -0.44 | 0.9979 |
| GTP - GppNHp | Cdc10 & Cdc11 | EDTA | 0.006 | 0.036 | 62.1 | 0.17 | 1.0000 |
| GDP - (GTP/GDP) | Cdc10 & Cdc11 | EDTA | 0.138 | 0.066 | 59.9 | 2.08 | 0.3120 |
| GDP - GTPyS | Cdc10 & Cdc11 | EDTA | 0.326 | 0.057 | 73.7 | 5.68 | 0.0000 |
| GDP - GppNHp | Cdc10 & Cdc11 | EDTA | 0.349 | 0.056 | 73.7 | 6.24 | 0.0000 |
| (GTP/GDP) - GTPyS | Cdc10 & Cdc11 | EDTA | 0.188 | 0.052 | 99.0 | 3.65 | 0.0055 |
| (GTP/GDP) - GppNHp | Cdc10 & Cdc11 | EDTA | 0.211 | 0.050 | 100.4 | 4.23 | 0.0007 |
| GTPyS - GppNHp | Cdc10 & Cdc11 | EDTA | 0.022 | 0.037 | 71.3 | 0.60 | 0.9906 |
| noGXP - GTP | Cdc10 & Shs1 | MgCl<sub>2</sub> | -0.903 | 0.074 | 31.0 | -12.20 | 0.0000 |
| *noGXP - GDP* | Cdc10 & Shs1 | MgCl<sub>2</sub> | -0.753 | 0.070 | 36.4 | -10.69 | 0.0000 |
| *noGXP - (GTP/GDP)* | Cdc10 & Shs1 | MgCl<sub>2</sub> | -0.899 | 0.074 | 31.2 | -12.16 | 0.0000 |
| *noGXP - GTPyS* | Cdc10 & Shs1 | MgCl<sub>2</sub> | -0.760 | 0.071 | 36.1 | -10.75 | 0.0000 |
| *noGXP - GppNHp* | Cdc10 & Shs1 | MgCl<sub>2</sub> | 0.009 | 0.039 | 80.8 | 0.23 | 0.9999 |
| *GTP - GDP* | Cdc10 & Shs1 | MgCl<sub>2</sub> | 0.149 | 0.094 | 24.2 | 1.59 | 0.6118 |
| *GTP - (GTP/GDP)* | Cdc10 & Shs1 | MgCl<sub>2</sub> | 0.004 | 0.097 | 22.8 | 0.04 | 1.0000 |
| *GTP - GTPyS* | Cdc10 & Shs1 | MgCl<sub>2</sub> | 0.143 | 0.094 | 24.1 | 1.52 | 0.6548 |
| *GTP - GppNHp* | Cdc10 & Shs1 | MgCl<sub>2</sub> | 0.912 | 0.074 | 30.8 | 12.40 | 0.0000 |
| *GDP - (GTP/GDP)* | Cdc10 & Shs1 | MgCl<sub>2</sub> | -0.145 | 0.094 | 24.2 | -1.55 | 0.6377 |
| *GDP - GTPyS* | Cdc10 & Shs1 | MgCl<sub>2</sub> | -0.006 | 0.091 | 25.9 | -0.07 | 1.0000 |
| *GDP - GppNHp* | Cdc10 & Shs1 | MgCl<sub>2</sub> | 0.762 | 0.070 | 36.1 | 10.88 | 0.0000 |
| *(GTP/GDP) - GTPyS* | Cdc10 & Shs1 | MgCl<sub>2</sub> | 0.139 | 0.094 | 24.2 | 1.48 | 0.6802 |
| *(GTP/GDP) - GppNHp* | Cdc10 & Shs1 | MgCl<sub>2</sub> | 0.908 | 0.073 | 30.9 | 12.36 | 0.0000 |
| *GTPyS - GppNHp* | Cdc10 & Shs1 | MgCl<sub>2</sub> | 0.769 | 0.070 | 35.8 | 10.95 | 0.0000 |
| *noGXP - GTP* | Cdc10 & Shs1 | EDTA | 0.003 | 0.035 | 60.8 | 0.08 | 1.0000 |
| *noGXP - GDP* | Cdc10 & Shs1 | EDTA | -0.521 | 0.062 | 51.4 | -8.38 | 0.0000 |
| *noGXP - (GTP/GDP)* | Cdc10 & Shs1 | EDTA | -0.275 | 0.053 | 86.9 | -5.16 | 0.0000 |
| *noGXP - GTPyS* | Cdc10 & Shs1 | EDTA | -0.003 | 0.036 | 63.9 | -0.07 | 1.0000 |
| *noGXP - GppNHp* | Cdc10 & Shs1 | EDTA | 0.003 | 0.035 | 60.8 | 0.09 | 1.0000 |
| *GTP - GDP* | Cdc10 & Shs1 | EDTA | -0.524 | 0.062 | 51.3 | -8.45 | 0.0000 |
| *GTP - (GTP/GDP)* | Cdc10 & Shs1 | EDTA | -0.278 | 0.053 | 86.9 | -5.24 | 0.0000 |
| *GTP - GTPyS* | Cdc10 & Shs1 | EDTA | -0.006 | 0.036 | 62.3 | -0.16 | 1.0000 |
| *GTP - GppNHp* | Cdc10 & Shs1 | EDTA | 0.000 | 0.035 | 59.1 | 0.00 | 1.0000 |
| *GDP - (GTP/GDP)* | Cdc10 & Shs1 | EDTA | 0.246 | 0.074 | 44.0 | 3.34 | 0.0198 |
| *GDP - GTPyS* | Cdc10 & Shs1 | EDTA | 0.519 | 0.062 | 51.6 | 8.31 | 0.0000 |
| *GDP - GppNHp* | Cdc10 & Shs1 | EDTA | 0.525 | 0.062 | 51.3 | 8.46 | 0.0000 |
| *(GTP/GDP) - GTPyS* | Cdc10 & Shs1 | EDTA | 0.272 | 0.053 | 86.8 | 5.09 | 0.0000 |
| *(GTP/GDP) - GppNHp* | Cdc10 & Shs1 | EDTA | 0.278 | 0.053 | 86.9 | 5.24 | 0.0000 |
| *GTPyS - GppNHp* | Cdc10 & Shs1 | EDTA | 0.006 | 0.036 | 62.3 | 0.16 | 1.0000 |

**Supplementary Table 3.** Sample sizes for quantification of peroxisomal recruitment of mNeonGreen-fused septins to Pex3M1-K4 5-mCherry-Cdc3/Cdc12 (wild-type and mutants) anchors (related to Fig. 3g,h). The “-” control displays only Pex3M1-K4 5-mCherry on peroxisomes.

| Anchor construct (Pex3<sub>M1-K45</sub>-mCherry fused to) | mNeonGreen-fused binding partner | n (number of cells quantified / independent experiments) |
| --- | --- | --- |
| *Cdc3* | Cdc10 | 117 / 2 |
| *Cdc3<sub>K287D,D289A</sub>* | Cdc10 | 104 / 2 |
| *Cdc3<sub>W364D</sub>* | Cdc10 | 111 / 2 |
| *Cdc3<sub>Δα0</sub>* | Cdc10 | 119 / 2 |
| - | Cdc10 | 116 / 2 |
| *Cdc3* | Cdc12 | 110 / 2 |
| *Cdc3<sub>K287D,D289A</sub>* | Cdc12 | 109 / 2 |
| *Cdc3<sub>W364D</sub>* | Cdc12 | 109 / 2 |
| *Cdc3<sub>Δα0</sub>* | Cdc12 | 112 / 2 |
| - | Cdc12 | 119 / 2 |
| *Cdc12* | Cdc3 | 111 / 2 |
| *Cdc12<sub>K180D,D182A</sub>* | Cdc3 | 120 / 2 |
| *Cdc12<sub>R263A</sub>* | Cdc3 | 107 / 2 |
| *Cdc12<sub>Δα0</sub>* | Cdc3 | 101 / 2 |
| - | Cdc3 | 132 / 2 |
| *Cdc12* | Cdc11 | 106 / 2 |
| *Cdc12<sub>K180D,D182A</sub>* | Cdc11 | 112 / 2 |
| *Cdc12<sub>R263A</sub>* | Cdc11 | 106 / 2 |
| *Cdc12<sub>Δα0</sub>* | Cdc11 | 101 / 2 |
| - | Cdc11 | 116 / 2 |
| *Cdc12* | Shs1 | 127 / 2 |
| *Cdc12<sub>K180D,D182A</sub>* | Shs1 | 111 / 2 |
| *Cdc12<sub>R263A</sub>* | Shs1 | 104 / 2 |
| *Cdc12<sub>Δα0</sub>* | Shs1 | 106 / 2 |
| - | Shs1 | 113 / 2 |

**Supplementary Table 4.** Combinatorial degeneracy of contiguous fragments from linear septin rods. The table lists all possible contiguous fragments generated by breaks of linear D-C-B-A-A-B-C-D octamers and B-A-A-B and A-B-C-D tetramers, assuming dissociation occurs at any interface but only between adjacent subunits and that distinct fragment orientations in the original rod (e.g., A-B versus B-A) are counted as separate microstates. For each approximate fragment molecular weight, the degeneracy (number of distinct contiguous sequences with that mass), the contributing fragments, and the number of exposed G-interfaces per fragment are shown. A-B and C-D are treated as engaged interfaces; all other G-interfaces are considered exposed. In this model, A maps to variants of His6-MBP-Cdc10 (∼80 kDa), B to variants of His6-Cdc3 (∼40 kDa), C to His6-MBP-Cdc12G32-G314 (∼80 kDa), and D to His6-Cdc11G20-G298 or His6-Shs1G21-S339 (∼40 kDa).

| Approximate fragment molecular weight [kDa] | Degeneracy | Contributing fragments | Exposed G-interfaces |
| --- | --- | --- | --- |
| *Octamers type D-C-B-A-A-B-C-D* |  |  |  |
| 480 | 1 | D-C-B-A-A-B-C-D | 0 |
| 440 | 2 | C-B-A-A-B-C-D; D-C-B-A-A-B-C | 1 |
| 400 | 1 | C-B-A-A-B-C | 2 |
| 360 | 2 | B-A-A-B-C-D; D-C-B-A-A-B | 0 |
| 320 | 4 | A-A-B-C-D; B-A-A-B-C; C-B-A-A-B; D-C-B-A-A | 1 |
| 280 | 2 | A-A-B-C; C-B-A-A | 2 |
| 240 | 3 | A-B-C-D; B-A-A-B; D-C-B-A | 0 |
| 200 | 4 | A-A-B; A-B-C; B-A-A; C-B-A | 1 |
| 160 | 3 | A-A; B-C-D; D-C-B | 2; 1; 1 |
| 120 | 6 | A-B; B-A; B-C; C-B; C-D; D-C | 0; 0; 2; 2; 0; 0 |
| *Tetramers type B-A-A-B* |  |  |  |
| 240 | 1 | B-A-A-B | 0 |
| 200 | 2 | A-A-B; B-A-A | 1 |
| 160 | 1 | A-A | 2 |
| 120 | 2 | A-B; B-A | 0 |
| *Tetramers type A-B-C-D* |  |  |  |
| 240 | 1 | A-B-C-D | 0 |
| 200 | 1 | A-B-C | 1 |
| 160 | 1 | B-C-D | 1 |
| 120 | 3 | A-B; B-C; C-D | 0; 2; 0 |

**Supplementary Table 5.** Mapping of septin mutations analyzed in the viability assay. The table lists all point mutations as well as N- and C-terminal truncations introduced in Cdc10, Cdc3, Cdc12, Cdc11, and Shs1 that were tested in the viability assay in Supplementary Fig. 5a. Mutations target conserved residues and regions involved in nucleotide-binding (K/R(P-loop)E, K/RxD(G4)ExA), magnesium-coordination (T/S(P-loop)A), G-interface stabilization (E(α4)R, W(βb-βc)D), and NC-interface formation (ΔNTE, ΔCTE). For truncation mutants, the remaining amino acid range of each protein (ΔNTE, ΔCTE) is indicated.

| Mutation | Cdc10 | Cdc3 | Cdc12 | Cdc11 | Shs1 |
| --- | --- | --- | --- | --- | --- |
| K/R(P-loop)E | K45E | K132E | K47E | R35E | K36E |
| T/S(P-loop)A | S46A | T133A | T48A | S36A | T37A |
| K/RxD(G4)ExA | K180D, D182A | K287D, D289A | K180D, D182A | K172D, D174A | R218D, D220A |
| E(α4)R | E188R | E295R | E188R | E180R | E226R |
| W(βb-βc)D | W255D | W364D | W267D | W251D | W292D |
| ΔNTE | G30-end | G117-end | G32-end | G20-end | G21-end |
| ΔCTE | M1-L302 | M1-A410 | M1-G314 | M1-G298 | M1-S339 |

**Supplementary Table 6.** Pairwise comparisons of nucleotide effects on SEPT9-SEPT9 NC-interface NanoBiT luminescence in the presence or absence of SEPT7. Tukey-adjusted pairwise contrasts from the three-way GLS model (normalized RLU ∼ SEPT7 × buffer × nucleotide) are shown for conditions without (top) and with (bottom) the G-interface partner SEPT7. Estimates represent differences in estimated marginal means of normalized relative light units; SE, pooled standard error of the contrast; *df*, residual degrees of freedom; *t* ratio, test statistic; adjusted *P*, Tukey HSD-corrected *P*-value. Adjustments were applied separately within each buffer × SEPT7 combination. Positive estimates indicate a higher signal for the first term in the contrast. Both NC-interface formation above baseline (noGXP) and nucleotide-specific differences in signal magnitude are observed exclusively in the presence of SEPT7, confirming that G-interface engagement is a prerequisite for nucleotide-dependent NC-interface assembly. Unlike for yeast Cdc10-Cdc10, GDP and GTP/GDP retain partial activity under EDTA conditions, suggesting that human SEPT9-SEPT7 G-interface formation is less strictly Mg^2+^-dependent.

| Contrast | G | Buffer | Estimate | SE | df | t ratio | adjusted P |
| --- | --- | --- | --- | --- | --- | --- | --- |
| noGXP - GTP | - | MgCl<sub>2</sub> | -0.005 | 0.012 | 61.5 | -0.44 | 0.9979 |
| noGXP - GDP | - | MgCl<sub>2</sub> | 0.007 | 0.011 | 53.0 | 0.60 | 0.9905 |
| noGXP - (GTP/GDP) | - | MgCl<sub>2</sub> | 0.002 | 0.012 | 56.3 | 0.16 | 1.0000 |
| noGXP - GTPyS | - | MgCl<sub>2</sub> | -0.007 | 0.013 | 62.3 | -0.53 | 0.9948 |
| noGXP - GppNHp | - | MgCl<sub>2</sub> | 0.003 | 0.012 | 55.4 | 0.27 | 0.9998 |
| GTP - GDP | - | MgCl<sub>2</sub> | 0.012 | 0.012 | 57.9 | 1.03 | 0.9054 |
| GTP - (GTP/GDP) | - | MgCl<sub>2</sub> | 0.007 | 0.012 | 60.5 | 0.59 | 0.9911 |
| GTP - GTPyS | - | MgCl<sub>2</sub> | -0.001 | 0.013 | 65.1 | -0.09 | 1.0000 |
| GTP - GppNHp | - | MgCl<sub>2</sub> | 0.009 | 0.012 | 59.8 | 0.70 | 0.9809 |
| GDP - (GTP/GDP) | - | MgCl<sub>2</sub> | -0.005 | 0.011 | 51.3 | -0.44 | 0.9977 |
| GDP - GTPyS | - | MgCl<sub>2</sub> | -0.013 | 0.012 | 59.0 | -1.12 | 0.8709 |
| GDP - GppNHp | - | MgCl<sub>2</sub> | -0.004 | 0.011 | 50.2 | -0.33 | 0.9994 |
| (GTP/GDP) - GTPyS | - | MgCl<sub>2</sub> | -0.008 | 0.012 | 61.4 | -0.68 | 0.9830 |
| (GTP/GDP) - GppNHp | - | MgCl<sub>2</sub> | 0.001 | 0.011 | 53.9 | 0.11 | 1.0000 |
| GTPyS - GppNHp | - | MgCl<sub>2</sub> | 0.010 | 0.012 | 60.7 | 0.79 | 0.9677 |
| noGXP - GTP | - | EDTA | -0.002 | 0.009 | 36.3 | -0.19 | 1.0000 |
| noGXP - GDP | - | EDTA | 0.004 | 0.008 | 32.1 | 0.46 | 0.9971 |
| noGXP - (GTP/GDP) | - | EDTA | 0.000 | 0.009 | 35.2 | -0.05 | 1.0000 |
| noGXP - GTPyS | - | EDTA | 0.000 | 0.009 | 34.7 | 0.03 | 1.0000 |
| noGXP - GppNHp | - | EDTA | 0.002 | 0.009 | 33.3 | 0.24 | 0.9999 |
| GTP - GDP | - | EDTA | 0.006 | 0.009 | 33.7 | 0.65 | 0.9858 |
| GTP - (GTP/GDP) | - | EDTA | 0.001 | 0.009 | 36.7 | 0.15 | 1.0000 |
| GTP - GTPyS | - | EDTA | 0.002 | 0.009 | 36.1 | 0.22 | 0.9999 |
| GTP - GppNHp | - | EDTA | 0.004 | 0.009 | 34.8 | 0.43 | 0.9979 |
| GDP - (GTP/GDP) | - | EDTA | -0.004 | 0.008 | 32.5 | -0.51 | 0.9955 |
| GDP - GTPyS | - | EDTA | -0.004 | 0.008 | 31.9 | -0.43 | 0.9979 |
| GDP - GppNHp | - | EDTA | -0.002 | 0.008 | 30.3 | -0.22 | 0.9999 |
| (GTP/GDP) - GTPyS | - | EDTA | 0.001 | 0.009 | 35.0 | 0.08 | 1.0000 |
| (GTP/GDP) - GppNHp | - | EDTA | 0.002 | 0.009 | 33.7 | 0.29 | 0.9997 |
| GTPyS - GppNHp | - | EDTA | 0.002 | 0.009 | 33.1 | 0.21 | 0.9999 |
| noGXP - GTP | SEPT7 | MgCl<sub>2</sub> | -0.935 | 0.093 | 4.9 | -10.00 | 0.0013 |
| noGXP - GDP | SEPT7 | MgCl<sub>2</sub> | -0.121 | 0.028 | 28.7 | -4.34 | 0.0020 |
| noGXP - (GTP/GDP) | SEPT7 | MgCl<sub>2</sub> | -0.727 | 0.078 | 5.8 | -9.28 | 0.0008 |
| noGXP - GTPyS | SEPT7 | MgCl<sub>2</sub> | 0.004 | 0.016 | 72.0 | 0.28 | 0.9998 |
| noGXP - GppNHp | SEPT7 | MgCl<sub>2</sub> | 0.010 | 0.015 | 71.7 | 0.63 | 0.9882 |
| GTP - GDP | SEPT7 | MgCl<sub>2</sub> | 0.814 | 0.096 | 5.2 | 8.46 | 0.0023 |
| GTP - (GTP/GDP) | SEPT7 | MgCl<sub>2</sub> | 0.208 | 0.121 | 5.1 | 1.72 | 0.5719 |
| GTP - GTPyS | SEPT7 | MgCl<sub>2</sub> | 0.939 | 0.093 | 4.9 | 10.06 | 0.0013 |
| GTP - GppNHp | SEPT7 | MgCl<sub>2</sub> | 0.945 | 0.093 | 4.9 | 10.12 | 0.0013 |
| GDP - (GTP/GDP) | SEPT7 | MgCl<sub>2</sub> | -0.606 | 0.082 | 6.2 | -7.43 | 0.0020 |
| GDP - GTPyS | SEPT7 | MgCl<sub>2</sub> | 0.125 | 0.028 | 28.5 | 4.53 | 0.0012 |
| GDP - GppNHp | SEPT7 | MgCl<sub>2</sub> | 0.130 | 0.027 | 28.3 | 4.77 | 0.0007 |
| (GTP/GDP) - GTPyS | SEPT7 | MgCl<sub>2</sub> | 0.731 | 0.078 | 5.8 | 9.34 | 0.0008 |
| (GTP/GDP) - GppNHp | SEPT7 | MgCl<sub>2</sub> | 0.737 | 0.078 | 5.8 | 9.42 | 0.0008 |
| GTPyS - GppNHp | SEPT7 | MgCl<sub>2</sub> | 0.005 | 0.015 | 71.0 | 0.36 | 0.9992 |
| noGXP - GTP | SEPT7 | EDTA | -0.005 | 0.011 | 51.0 | -0.43 | 0.9980 |
| noGXP - GDP | SEPT7 | EDTA | -0.208 | 0.032 | 18.4 | -6.43 | 0.0001 |
| noGXP - (GTP/GDP) | SEPT7 | EDTA | -0.134 | 0.025 | 30.3 | -5.37 | 0.0001 |
| noGXP - GTPyS | SEPT7 | EDTA | -0.020 | 0.013 | 64.3 | -1.57 | 0.6197 |
| noGXP - GppNHp | SEPT7 | EDTA | 0.001 | 0.010 | 46.1 | 0.11 | 1.0000 |
| GTP - GDP | SEPT7 | EDTA | -0.203 | 0.032 | 18.7 | -6.26 | 0.0001 |
| GTP - (GTP/GDP) | SEPT7 | EDTA | -0.130 | 0.025 | 30.7 | -5.14 | 0.0002 |
| GTP - GTPyS | SEPT7 | EDTA | -0.015 | 0.013 | 65.7 | -1.16 | 0.8538 |
| GTP - GppNHp | SEPT7 | EDTA | 0.006 | 0.011 | 50.2 | 0.54 | 0.9944 |
| GDP - (GTP/GDP) | SEPT7 | EDTA | 0.073 | 0.039 | 19.1 | 1.86 | 0.4555 |
| GDP - GTPyS | SEPT7 | EDTA | 0.188 | 0.033 | 19.3 | 5.68 | 0.0002 |
| GDP - GppNHp | SEPT7 | EDTA | 0.209 | 0.032 | 18.3 | 6.48 | 0.0001 |
| (GTP/GDP) - GTPyS | SEPT7 | EDTA | 0.115 | 0.026 | 31.9 | 4.41 | 0.0014 |
| (GTP/GDP) - GppNHp | SEPT7 | EDTA | 0.135 | 0.025 | 30.1 | 5.43 | 0.0001 |
| GTPyS - GppNHp | SEPT7 | EDTA | 0.021 | 0.012 | 64.0 | 1.67 | 0.5555 |

**Supplementary Table 7.** Cross-subunit meta-analysis of RMSD95 effects. For each mechanistic contrast, per-subunit Hodges-Lehmann estimates of the RMSD95 difference (median of all pairwise differences between conditions) were summarized across the eight septins. For the two marginal contrasts, dimer - monomer (avg GXP + apo) and GXP - apo (avg dimer + monomer), the per-subunit effect was obtained by averaging the two stratum-specific Hodges-Lehmann estimates (e.g., HL of dimer - monomer in GXP and HL of dimer - monomer in apo) to give a single marginal effect per subunit collapsed across the second factor. The four remaining rows report contrasts evaluated within a single stratum and use the per-subunit Hodges-Lehmann estimate directly. n, number of subunits included; Negative/Positive, number of subunits showing a decrease versus an increase in RMSD95 for the first condition relative to the second; Median ΔRMSD95, median Hodges-Lehmann effect across subunits; Range, observed range of per-subunit Hodges-Lehmann effects [min, max]; Sign test *P*, exact binomial *P*-value for directional consistency (null: 50% negative); Signed-rank *P*, one-sided Wilcoxon signed-rank *P*-value testing whether the median effect is less than zero (whether the first condition stabilizes the complexes compared to the second). Despite limited power at the level of individual subunits, the direction of effects was fully consistent across all eight septins for each contrast, with dimerization producing a median stabilizing effect of −0.37 nm and nucleotide binding −0.15 nm.

| Contrast | n | Negative/<br>Positive | Median<br>ΔRMSD95 [nm] | Range<br>ΔRMSD95 [nm] | Sign<br>test *P* | Signed-rank *P* |
| --- | --- | --- | --- | --- | --- | --- |
| dimer - monomer (avg GXP + apo) | 8 | 8/0 | -0.366 | [-0.472, -0.133] | 0.0039 | 0.0071 |
| GXP - apo (avg dimer + monomer) | 8 | 8/0 | -0.153 | [-0.264, -0.033] | 0.0039 | 0.0071 |
| dimer - monomer (GXP only) | 8 | 8/0 | -0.316 | [-0.571, -0.155] | 0.0039 | 0.0071 |
| dimer - monomer (apo only) | 8 | 8/0 | -0.372 | [-0.772, -0.111] | 0.0039 | 0.0071 |
| GXP - apo (dimer only) | 8 | 8/0 | -0.097 | [-0.285, -0.039] | 0.0039 | 0.0071 |
| GXP - apo (monomer only) | 8 | 8/0 | -0.123 | [-0.424, -0.021] | 0.0039 | 0.0071 |

**Supplementary Table 8.** List of plasmids used in this study. pAc and pES are in-house constructed, pET15A-based plasmids for the expression of His6-tagged proteins in *E. coli*. pRS304-X3 is an in-house-constructed, pRS304-based vector containing the EasyClone 2.0 integration site 3 in chromosome X of *S. cerevisiae*^54^. aa-amino acid residue, fl - full-length, S - S tag, PMET - MET17 promoter, PCUP - CUP1 promoter, mNG - mNeonGreen, mCh - mCherry.

|  | Construct | Vector backbone | Insert | Source |
| --- | --- | --- | --- | --- |
| #1 | His<sub>6</sub>-MBP-Cdc10G | pAc | aa 30-end | Grupp *et al.*<sup>27</sup> |
| #2 | His<sub>6</sub>-MBP-Cdc12G | pAc | aa 32-314 | <sup>27</sup> |
| #3 | His<sub>6</sub>-Cdc3G | pETDuet-1 | aa 117-410 | <sup>27</sup> |
| #4 | His<sub>6</sub>-Cdc11G | pETDuet-1 | aa 20-298 | <sup>27</sup> |
| #5 | His<sub>6</sub>-Shs1G | pETDuet-1 | aa 21-339 | <sup>27</sup> |
| #6 | His<sub>6</sub>-Cdc10G, Cdc3G-S | pETDuet-1 | aa 30-end (Cdc10), aa 117-410 (Cdc3) | This study |
| #7 | His<sub>6</sub>-Cdc12G, Cdc11G-S | pETDuet-1 | aa 32-314 (Cdc12), aa 20-298 (Cdc11) | This study |
| #8 | His<sub>6</sub>-Cdc12G, Shs1G-S | pETDuet-1 | aa 32-314 (Cdc12), aa 21-339 (Shs1) | This study |
| #9 | His<sub>6</sub>-MBP-Cdc10G-LgBiT | pAc | aa 30-302 | This study |
| #10 | His<sub>6</sub>-MBP-Cdc12G-LgBiT | pAc | aa 32-314 | This study |
| #11 | His<sub>6</sub>-Cdc3G-SmBiT-SUMO | pE-SUMOprom Kan | aa 117-410 | This study |
| #12 | His<sub>6</sub>-Cdc11G-SmBiT-SUMO | pE-SUMOprom Kan | aa 20-298 | This study |
| #13 | His<sub>6</sub>-Shs1G-SmBiT-SUMO | pE-SUMOprom Kan | aa 21-339 | This study |
| #14 | His<sub>6</sub>-MBP-Cdc10 | pAc | fl | This study |
| #15 | His<sub>6</sub>-MBP-Cdc12ΔC | pAc | aa 1-314 | This study |
| #16 | His<sub>6</sub>-Cdc3ΔNC | pETDuet-1 | aa 81-410 | This study |
| #17 | His<sub>6</sub>-MBP-Cdc10ΔC-LgBiT | pAc | aa 1-302 | This study |
| #18 | His<sub>6</sub>-MBP-Cdc10ΔC-SmBiT | pAc | aa 1-302 | This study |
| #19 | His<sub>6</sub>-MBP-Cdc12ΔC-LgBiT | pAc | aa 1-314 | This study |
| #20 | His<sub>6</sub>-Cdc3ΔNC-SmBiT-SUMO | pE-SUMOprom Kan | aa 81-410 | This study |
| #21 | P<sub>MET</sub>-Cdc10 | pRS316 | fl | <sup>31</sup> |
| #22 | P<sub>MET</sub>-Cdc3 | pRS316 | fl | <sup>27</sup> |
| #23 | P<sub>MET</sub>-Cdc12 | pRS316 | fl | <sup>27</sup> |
| #24 | P<sub>MET</sub>-Cdc11 | pRS316 | fl | <sup>27</sup> |
| #25 | P<sub>MET</sub>-Shs1 | pRS316 | fl | <sup>27</sup> |
| #26 | P<sub>MET</sub>-Cdc10 | pRS313 | fl | <sup>31</sup> |
| #27 | P<sub>MET</sub>-Cdc10(K45E) | pRS313 | fl | This study |
| #28 | P<sub>MET</sub>-Cdc10(S46A) | pRS313 | fl | This study |
| #29 | P<sub>MET</sub>-Cdc10(K180D, D182A) | pRS313 | fl | This study |
| #30 | P<sub>MET</sub>-Cdc10(E188R) | pRS313 | fl | This study |
| #31 | P<sub>MET</sub>-Cdc10(W255D) | pRS313 | fl | This study |
| #32 | P<sub>MET</sub>-Cdc10ΔN | pRS313 | aa 30-end | <sup>31</sup> |
| #33 | P<sub>MET</sub>-Cdc10ΔC | pRS313 | aa 1-302 | This study |
| #34 | P<sub>MET</sub>-Cdc3 | pRS313 | fl | <sup>27</sup> |
| #35 | P<sub>MET</sub>-Cdc3(K132E) | pRS313 | fl | This study |
| #36 | P<sub>MET</sub>-Cdc3(T133A) | pRS313 | fl | This study |
| #37 | P<sub>MET</sub>-Cdc3(K287D, D289A) | pRS313 | fl | This study |
| #38 | P<sub>MET</sub>-Cdc3(E295R) | pRS313 | fl | This study |
| #39 | P<sub>MET</sub>-Cdc3(W364D) | pRS313 | fl | This study |
| #40 | P<sub>MET</sub>-Cdc3ΔN | pRS313 | aa 117-end | This study |
| #41 | P<sub>MET</sub>-Cdc3ΔC | pRS313 | aa 1-410 | This study |
| #42 | P<sub>MET</sub>-Cdc12 | pRS313 | fl | <sup>27</sup> |
| #43 | P<sub>MET</sub>-Cdc12(K47E) | pRS313 | fl | This study |
| #44 | P<sub>MET</sub>-Cdc12(T48A) | pRS313 | fl | This study |
| #45 | P<sub>MET</sub>-Cdc12(K180D, D182A) | pRS313 | fl | This study |
| #46 | P<sub>MET</sub>-Cdc12(E188R) | pRS313 | fl | This study |
| #47 | P<sub>MET</sub>-Cdc12(W267D) | pRS313 | fl | This study |
| #48 | P<sub>MET</sub>-Cdc12ΔN | pRS313 | aa 32-end | This study |
| #49 | P<sub>MET</sub>-Cdc12ΔC | pRS313 | aa 1-314 | This study |
| #50 | P<sub>MET</sub>-Cdc11 | pRS313 | fl | <sup>27</sup> |
| #51 | P<sub>MET</sub>-Cdc11(R35E) | pRS313 | fl | This study |
| #52 | P<sub>MET</sub>-Cdc11(S36A) | pRS313 | fl | This study |
| #53 | P<sub>MET</sub>-Cdc11(K172D, D174A) | pRS313 | fl | This study |
| #54 | P<sub>MET</sub>-Cdc11(E180R) | pRS313 | fl | This study |
| #55 | P<sub>MET</sub>-Cdc11(W251D) | pRS313 | fl | This study |
| #56 | P<sub>MET</sub>-Cdc11ΔN | pRS313 | aa 20-end | This study |
| #57 | P<sub>MET</sub>-Cdc11ΔC | pRS313 | aa 1-298 | This study |
| #58 | P<sub>MET</sub>-Shs1 | pRS313 | fl | <sup>27</sup> |
| #59 | P<sub>MET</sub>-Shs1(K36E) | pRS313 | fl | This study |
| #60 | P<sub>MET</sub>-Shs1(T37A) | pRS313 | fl | This study |
| #61 | P<sub>MET</sub>-Shs1(R218D, D220A) | pRS313 | fl | This study |
| #62 | P<sub>MET</sub>-Shs1(E226R) | pRS313 | fl | This study |
| #63 | P<sub>MET</sub>-Shs1(W292D) | pRS313 | fl | This study |
| #64 | P<sub>MET</sub>-Shs1ΔN | pRS313 | aa 21-end | This study |
| #65 | P<sub>MET</sub>-Shs1ΔC | pRS313 | aa 1-339 | This study |
| #66 | P<sub>MET</sub>-Cdc10-mNG | pRS306 | fl | This study |
| #67 | P<sub>MET</sub>-Cdc3-mNG | pRS306 | fl | This study |
| #68 | P<sub>MET</sub>-Cdc12-mNG | pRS306 | fl | This study |
| #69 | P<sub>MET</sub>-mNG-Cdc11 | pRS306 | fl | This study |
| #70 | P<sub>MET</sub>-Shs1-mNG | pRS306 | fl | This study |
| #71 | P<sub>CUP</sub>-Pex3(M1-K45)-mCh | pRS304-X3 | - | This study |
| #72 | P<sub>CUP</sub>-Pex3(M1-K45)-mCh-Cdc3 | pRS304-X3 | fl | This study |
| #73 | P<sub>CUP</sub>-Pex3(M1-K45)-mCh-Cdc3(K287D, D289A) | pRS304-X3 | fl | This study |
| #74 | P<sub>CUP</sub>-Pex3(M1-K45)-mCh-Cdc3(W364D) | pRS304-X3 | fl | This study |
| #75 | P<sub>CUP</sub>-Pex3(M1-K45)-mCh-Cdc3ΔN | pRS304-X3 | aa 117-end | This study |
| #76 | P<sub>CUP</sub>-Pex3(M1-K45)-mCh-Cdc12 | pRS304-X3 | fl | This study |
| #77 | P<sub>CUP</sub>-Pex3(M1-K45)-mCh-Cdc12(K180D, D182A) | pRS304-X3 | fl | This study |
| #78 | P<sub>CUP</sub>-Pex3(M1-K45)-mCh-Cdc12(R263A) | pRS304-X3 | fl | This study |
| #79 | P<sub>CUP</sub>-Pex3(M1-K45)-mCh-Cdc12ΔN | pRS304-X3 | aa 32-end | This study |
| #80 | His<sub>6</sub>-hSEPT9G | pETDuet-1 | aa 296-end | This study |
| #81 | His<sub>6</sub>-hSEPT6G | pETDuet-1 | aa 40-309 | This study |
| #82 | His<sub>6</sub>-MBP-SEPT2G | pAc | aa 35-308 | This study |
| #83 | His<sub>6</sub>-MBP-SEPT7G | pAc | aa 48-318 | <sup>27</sup> |
| #84 | His<sub>6</sub>-SEPT7G | pES | aa 48-318 | <sup>27</sup> |
| #85 | His<sub>6</sub>-MBP-SEPT7G-LgBiT | pAc | aa 48-318 | This study |
| #86 | His<sub>6</sub>-SEPT2G-LgBiT | pAc | aa 35-308 | This study |
| #87 | His<sub>6</sub>-hSEPT9G-SmBiT-SUMO | pE-SUMOpro Kan | aa 296-568 | This study |
| #88 | His<sub>6</sub>-SUMO-hSEPT6G-SmBiT | pE-SUMOpro Kan | aa 40-309 | This study |
| #89 | His<sub>6</sub>-hSEPT9ΔN | pETDuet-1 | aa 263-end | This study |
| #90 | His<sub>6</sub>-MBP-SEPT9ΔNC-LgBiT | pAc | aa 263-568 | This study |
| #91 | His<sub>6</sub>-MBP-SEPT9ΔNC-SmBiT | pAc | aa 263-568 | This study |
| #92 | His<sub>6</sub>-Cdc12G, Cdc11ΔC | pETDuet-1 | aa 32-314 (Cdc12), aa 1-298 (Cdc11) | This study |
| #93 | His<sub>6</sub>-Cdc11G(C43A, E74C, C137A, C138A, K248C) | pETDuet-1 | aa 20-298 | This study |
| #94 | His<sub>6</sub>-Cdc11G(C43A, C137A, C138A, E151C, K248C) | pETDuet-1 | aa 20-298 | This study |
| #95 | His<sub>6</sub>-Cdc11G(C43A, C137A, C138A, K182C, K248C) | pETDuet-1 | aa 20-298 | This study |
| #96 | His<sub>6</sub>-Cdc11G(C43A, C137A, C138A, E215C, K248C) | pETDuet-1 | aa 20-298 | This study |

**Supplementary Table 9.** List of all constructs appearing in the main and supplementary figures.

| Figure | Construct |
| --- | --- |
| Fig. 1a | #1–#8 |
| Fig. 1b | #1–#8 |
| Fig. 1c | #1 |
| Fig. 1d | #6–#8 |
| Fig. 2a | #1, #3 |
| Fig. 2b | #2, #4 |
| Fig. 2c | #2, #5 |
| Fig. 2d | #9, #11 |
| Fig. 2e | #10, #12 |
| Fig. 2c | #10, #13 |
| Fig. 2g | #9, #11 |
| Fig. 2h | #10, #12 |
| Fig. 2i | #10, #13 |
| Fig. 3a | #1, #3, #14 |
| Fig. 3b | #3, #14 |
| Fig. 3c | #1, #4–#5, #15–#16 |
| Fig. 3d | #3, #17–#18 |
| Fig. 3e | #1, #4–#5, #19–#20 |
| Fig. 3f | - |
| Fig. 3g | #67, #69–#71, #76–#79 |
| Fig. 3h | #66, #68, #71–#75 |
| Fig. 4a | #80–#84 |
| Fig. 4b | #83–#84 |
| Fig. 4c | #81–#82 |
| Fig. 4d | #80, #83 |
| Fig. 4e | #86, #88 |
| Fig. 4f | #85, #87 |
| Fig. 4g | #85, #87 |
| Fig. 4h | #83, #89 |
| Fig. 4i | #83, #90–#91 |
| Fig. 5a | #7, #92 |
| Fig. 5b | #92 |
| Fig. 5c | #92 |
| Fig. 5d | #92 |
| Fig. 5e | #92 |
| Fig. 5f | - |
| Fig. 5g | #92 |
| Fig. 5h | #92–#96 |
| Fig. 5i | #92–#96 |
| Fig. 5j | #92–#96 |
| Fig. 6a | - |
| Fig. 6b | - |
| Fig. 6c | - |
| Supplementary Fig. 1a | - |
| Supplementary Fig. 1b | - |
| Supplementary Fig. 1c | - |
| Supplementary Fig. 2a | #1–#5 |
| Supplementary Fig. 2b | #1–#2 |
| Supplementary Fig. 2c | #2 |
| Supplementary Fig. 2d | #4 |
| Supplementary Fig. 2e | #3 |
| Supplementary Fig. 2f | #5 |
| Supplementary Fig. 3a | #9, #11 |
| Supplementary Fig. 3b | #10, #12 |
| Supplementary Fig. 3c | #10, #13 |
| Supplementary Fig. 3d | #85, #87 |
| Supplementary Fig. 4a | #2, #4 |
| Supplementary Fig. 4b | #2, #5 |
| Supplementary Fig. 4c | #2, #4 |
| Supplementary Fig. 4d | #2, #5 |
| Supplementary Fig. 4e | #1, #3 |
| Supplementary Fig. 4f | #3, #14 |
| Supplementary Fig. 4g | #1, #4, #15–#16 |
| Supplementary Fig. 4h | #1, #4, #15–#16 |
| Supplementary Fig. 4i | #1, #4, #15–#16 |
| Supplementary Fig. 4j | #1, #5, #15–#16 |
| Supplementary Fig. 4k | #1, #5, #15–#16 |
| Supplementary Fig. 4l | #1, #5, #15–#16 |
| Supplementary Fig. 4m | #4, #14–#16 |
| Supplementary Fig. 4n | #4, #14–#16 |
| Supplementary Fig. 4o | #4, #14–#16 |
| Supplementary Fig. 4p | #5, #14–#16 |
| Supplementary Fig. 4q | #5, #14–#16 |
| Supplementary Fig. 4r | #5, #14–#16 |
| Supplementary Fig. 5a | #21–#65 |
| Supplementary Fig. 5b | - |
| Supplementary Fig. 5c | #67, #69–#71, #76–#79 |
| Supplementary Fig. 6a | #92 |
| Supplementary Fig. 6b | #92 |
| Supplementary Fig. 6c | #92 |
| Supplementary Fig. 6d | #92 |
| Supplementary Fig. 7a | - |
| Supplementary Fig. 7b | - |
| Supplementary Fig. 7c | - |
| Supplementary Fig. 8a | #2, #4, #93–#96 |
| Supplementary Fig. 8b | #4, #93–#96 |
| Supplementary Fig. 8c | #4, #93–#96 |
| Supplementary Fig. 8d | #4 |
| Supplementary Fig. 9a | #1–#8 |
| Supplementary Fig. 9b | #6–#8 |
| Supplementary Fig. 9c | #1–#5 |
| Supplementary Fig. 9d | #6–#8 |
| Supplementary Fig. 9e | #9–#13 |
| Supplementary Fig. 9f | #9–#13 |
| Supplementary Fig. 9g | #14–#16 |
| Supplementary Fig. 9h | #14–#16 |
| Supplementary Fig. 9i | #17–#20 |
| Supplementary Fig. 9j | #17–#20 |
| Supplementary Fig. 9k | #92 |
| Supplementary Fig. 9l | #92 |
| Supplementary Fig. 10a | #80–#84, #89 |
| Supplementary Fig. 10b | #80–#84, #89 |
| Supplementary Fig. 10c | #85–#88 |
| Supplementary Fig. 10d | #85 |
| Supplementary Fig. 10e | #85–#88 |
| Supplementary Fig. 10f | #90–#91 |
| Supplementary Fig. 10g | #90–#91 |
| Supplementary Fig. 11a | #92 |
| Supplementary Fig. 11b | #92 |
| Supplementary Fig. 11c | #92 |
| Supplementary Fig. 11d | #92 |
| Supplementary Fig. 12a | #92 |
| Supplementary Fig. 12b | #92 |
| Supplementary Fig. 13a | #93 |
| Supplementary Fig. 13b | #94 |
| Supplementary Fig. 13c | #95 |
| Supplementary Fig. 13d | #96 |

**Supplementary Table 10.** Cryo-EM data collection. , **refinement and validation statistics.**

|  | Cdc12-Cdc11-Cdc11-Cdc12 |
| --- | --- |
| **Data collection and processing** |  |
| Microscope | Thermo Fisher Titan Krios G4 |
| Detector | Falcon 4i |
| Magnification | 215,000× |
| Voltage (kV) | 300 |
| Electron exposure (e<sup>-</sup>/Å<sup>2</sup>) | 40 |
| Defocus range (μm) | −0.5 to −2.2 |
| Pixel size (Å) | 0.572 |
| Symmetry imposed | C<sub>2</sub> |
| Number of movies | 8195 |
| Final particle number | 70,904 |
| Map resolution (Å) | 2.52 |
| FSC threshold | 0.143 |
| Map resolution range (Å) | 2.2-2.9 |
| cFAR | 0.79 |
| **Refinement** |  |
| Initial model used | AlphaFold3 prediction of<br>Cdc12-Cdc11-Cdc11-Cdc12 |
| Model resolution (Å) | 2.9 |
| FSC threshold | 0.5 |
| Map sharpening B factor (Å<sup>2</sup>) | 47.9 |
| Model composition |  |
| Non-hydrogen atoms | 11,021 |
| Protein residues | 1,118 |
| Ligands | 4 |
| B factors (Å<sup>2</sup>) |  |
| Protein | 72.82 |
| Ligand | 88.27 |
| R.m.s. deviations |  |
| Bond lengths (Å) | 0.005 |
| Bond angles (°) | 0.459 |
| **Validation** |  |
| MolProbity score | 2.48 |
| Clashscore | 28.22 |
| Poor rotamers (%) | 1.93 |
| Ramachandran plot |  |
| Favored (%) | 95.28 |
| Allowed (%) | 4.36 |
| Disallowed (%) | 0.36 |

**Supplementary Table 11.**
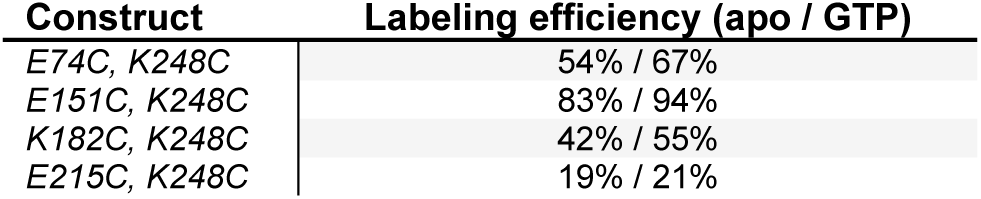
Spin-labeling efficiencies determined by cw-EPR.

| Construct | Labeling efficiency (apo / GTP) |
| --- | --- |
| E74C, K248C | 54% / 67% |
| E151C, K248C | 83% / 94% |
| K182C, K248C | 42% / 55% |
| E215C, K248C | 19% / 21% |

**Supplementary Table 12.** DEER acquisition parameters for spin-labeled Cdc11 variants. All measurements were performed at 80 K in Q-band on a Bruker ELEXSYS E580 spectrometer using a dielectric ring Q-band resonator, as described in Materials and Methods. *Δν*, frequency offset between pump and observer (*νpump* = *νobs* + *Δν*); *Δt*, dwell time of the pump pulse; *τ1*, first interpulse delay; *τ2*, dipolar evolution time; SRT, shot repetition time; *tgate*, integration gate width. ^a^For constructs #93, #95, and #96, *νpump* was set to the maximum of the nitroxide spectrum, and the observer was high-field shifted by ∼1.8 mT, corresponding to *Δν* ≈ 47–57 MHz across measurements. For construct #94, *Δν* was set to a fixed value of +80 MHz (see Materials and Methods).

| Parameter | #93<br>(E74C,K248C) |  | #94<br>(E151C,K248C) |  | #95<br>(K182C,K248C) |  | #96<br>(E215C,K248C) |  |
| --- | --- | --- | --- | --- | --- | --- | --- | --- |
|  | apo | GTP | apo | GTP | apo | GTP | apo | GTP |
| $t_{\pi/2,\text{obs}}$ (ns) | 18 | 18 | 32 | 32 | 18 | 18 | 18 | 18 |
| $t_{\pi,\text{obs}}$ (ns) | 36 | 36 | 32 | 32 | 36 | 36 | 36 | 36 |
| $t_{\pi,\text{pump}}$ (ns) | 36 | 36 | 14 | 14 | 36 | 36 | 36 | 36 |
| $\Delta\nu$ (MHz)<sup>a</sup> | +51 | +53 | +80 | +80 | +51 | +57 | +51 | +47 |
| $\Delta t$ (ns) | 16 | 16 | 12 | 12 | 8 | 8 | 8 | 16 |
| $\tau_1$ (μs) | 0.20 | 0.20 | 2.4 | 2.4 | 0.20 | 0.20 | 0.20 | 0.20 |
| $\tau_2$ (μs) | 3.6 | 3.2 | 3.8 | 3.8 | 2.4 | 2.4 | 1.6 | 3.2 |
| SRT (ms) | 3 | 3 | 1.5 | 1.5 | 3 | 3 | 3 | 3 |
| $t_{\text{gate}}$ (ns) | 36 | 108 | 32 | 32 | 114 | 124 | 102 | 100 |

**Supplementary Table 13.**
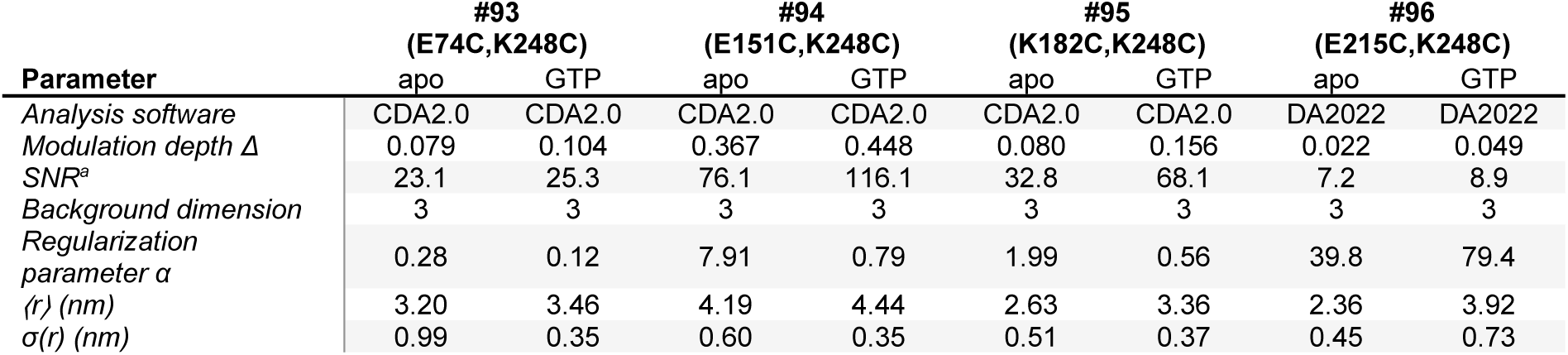
DEER analysis parameters and results. CDA2.0, ComparativeDeerAnalyzer 2.0; DA2022, DeerAnalysis2022; *Δ*, modulation depth of the dipolar evolution function; ⟨*r*⟩, mean of the distance distribution *P(r)*; *σ(r)*, standard deviation of *P(r)*. For constructs #93–#95, the regularization parameter *α* refers to the Tikhonov regularization component selected based on the best overlap with the DEERNet^93^ neural network solution within CDA2.0^91^, whereas ⟨*r*⟩ and *σ(r)* were obtained from the CDA2.0 consensus distance distributions. For construct #96, *α*, ⟨*r*⟩, and *σ(r)* were obtained from L-curve-selected Tikhonov regularization in DeerAnalysis2022^92^. ^a^Signal-to-noise ratio with respect to modulation, calculated as *Δ/rmsd*, where *rmsd* is the root-mean-square deviation of the time-domain fit normalized to the maximum of the signal.

## Supplementary Movies

**Supplementary Movie 1.**
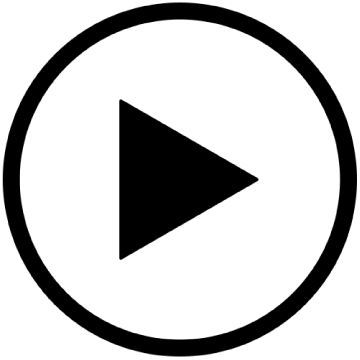
Flexibility of the SUE-βββ in Cdc3:apo. Molecular dynamics trajectory (700 ns) of Cdc3 in the apo state showing pronounced Cdc11-like unfolding of the SUE-βββ motif (pale green).

**Supplementary Movie 2.**
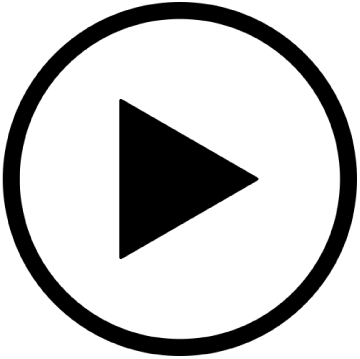
Flexibility of the SUE-βββ in SEPT7:GDP. Molecular dynamics trajectory (700 ns) of SEPT7 bound to GDP illustrating marked unfolding of the SUE-βββ (pale green) accompanied by partial GDP dissociation.

**Supplementary Movie 3.**
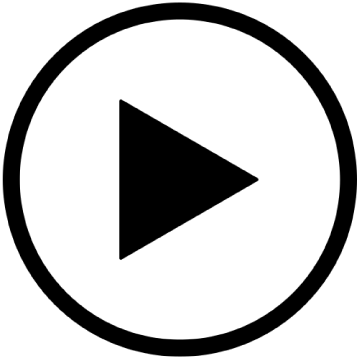
Nucleotide ejection by Cdc10. Molecular dynamics trajectory (700 ns) of Cdc10:GDP showing complete dissociation of GDP. The SUE-βββ is highlighted in pale green.

**Supplementary Movie 4.**
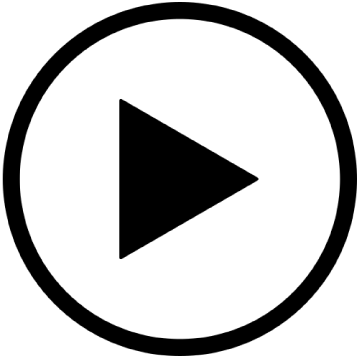
Nucleotide ejection by SEPT7. Molecular dynamics trajectory (700 ns) of SEPT7:GDP showing full release of GDP. The SUE-βββ is highlighted in pale green.

## References

1. Hartwell, L. H. Genetic control of the cell division cycle in yeast. IV. Genes controlling bud emergence and cytokinesis. Exp. Cell Res. 69, 265–276 (1971).

2. Longtine, M. S. et al. The septins: Roles in cytokinesis and other processes. Curr. Opin. Cell Biol. 8, 106–119 (1996).

3. Shuman, B. & Momany, M. Septins From Protists to People. Front. Cell Dev. Biol. 9, 1–7 (2022).

4. Grupp, B. & Gronemeyer, T. A biochemical view on the septins, a less known component of the cytoskeleton. Biol. Chem. 404, 1–13 (2023).

5. Cavini, I. A. et al. The Structural Biology of Septins and Their Filaments: An Update. Front. Cell Dev. Biol. 9, 1–25 (2021).

6. Sirajuddin, M. et al. Structural insight into filament formation by mammalian septins. Nature 449, 311–315 (2007).

7. Bertin, A. et al. Saccharomyces cerevisiae septins: Supramolecular organization of heterooligomers and the mechanism of filament assembly. Proc. Natl. Acad. Sci. U. S. A. 105, 8274–8279 (2008).

8. Mendonça, D. C. et al. A revised order of subunits in mammalian septin complexes. Cytoskeleton 76, 457–466 (2019).

9. Soroor, F. et al. Revised subunit order of mammalian septin complexes explains their in vitro polymerization properties. Mol. Biol. Cell 32, 289–300 (2021).

10. Bi, E. & Park, H. O. Cell polarization and cytokinesis in budding yeast. Genetics 191, 347–387 (2012).

11. Marquardt, J., Chen, X. & Bi, E. Architecture, remodeling, and functions of the septin cytoskeleton. Cytoskeleton 76, 7–14 (2019).

12. Mostowy, S. & Cossart, P. Septins: The fourth component of the cytoskeleton. Nat. Rev. Mol. Cell Biol. 13, 183–194 (2012).

13. Field, C. M. & Kellogg, D. Septins: Cytoskeletal polymers or signalling GTPases? Trends Cell Biol. 9, 387–394 (1999).

14. Hecht, M., Alber, N., Marhoffer, P., Johnsson, N. & Gronemeyer, T. The concerted action of SEPT9 and EPLIN modulates the adhesion and migration of human fibroblasts. Life Sci. Alliance 7, 1–17 (2024).

15. Akhmetova, K. A., Chesnokov, I. N. & Fedorova, S. A. Functional Characterization of Septin Complexes. Mol. Biol. (Moscow) 52, 155–171 (2018).

16. Kozubowski, L., Larson, J. R. & Tatchell, K. Role of the Septin Ring in the Asymmetric Localization of Proteins at the Mother-Bud Neck in *Saccharomyces cerevisiae*. Mol. Biol. Cell 16, 3455–3466 (2005).

17. Mitchison, T. J. & Field, C. M. Cytoskeleton: What does GTP do for septins? Current Biology 12, 788–790 (2002).

18. Zent, E. & Wittinghofer, A. Human septin isoforms and the GDP-GTP cycle. Biol. Chem. 395, 169–180 (2014).

19. Mendoza, M., Hyman, A. A. & Glotzer, M. GTP Binding Induces Filament Assembly of a Recombinant Septin. Current Biology 12, 1858–1863 (2002).

20. Vrabioiu, A. M., Gerber, S. A., Gygi, S. P., Field, C. M. & Mitchison, T. J. The Majority of the Saccharomyces cerevisiae Septin Complexes Do Not Exchange Guanine Nucleotides. Journal of Biological Chemistry 279, 3111–3118 (2004).

21. Versele, M. & Thorner, J. Septin collar formation in budding yeast requires GTP binding and direct phosphorylation by the PAK, Cla4. Journal of Cell Biology 164, 701–715 (2004).

22. Abbey, M., Gaestel, M. & Menon, M. B. Septins: Active GTPases or just GTP-binding proteins? Cytoskeleton 76, 55–62 (2019).

23. Kuo, Y. C. et al. SEPT12 mutations cause male infertility with defective sperm annulus. Hum. Mutat. 33, 710–719 (2012).

24. Angelis, D. & Spiliotis, E. T. Septin mutations in human cancers. Front. Cell Dev. Biol. 4, 1–17 (2016).

25. Brognara, G., Pereira, H. D. M., Brandão-Neto, J., Araujo, A. P. U. & Garratt, R. C. Revisiting SEPT7 and the slippage of β-strands in the septin family. J. Struct. Biol. 207, 67–73 (2019).

26. Zeraik, A. E. et al. Crystal structure of a Schistosoma mansoni septin reveals the phenomenon of strand slippage in septins dependent on the nature of the bound nucleotide. Journal of Biological Chemistry 289, 7799–7811 (2014).

27. Grupp, B. et al. Interface integrity in septin protofilaments is maintained by an arginine residue conserved from yeast to man. Mol. Biol. Cell 36, (2025).

28. Grupp, B., Lemkul, J. A. & Gronemeyer, T. An in silico approach to determine inter-subunit affinities in human septin complexes. Cytoskeleton 80, 141–152 (2023).

29. Mendonça, D. C. et al. Structural Insights into Ciona intestinalis Septins: Complexes Suggest a Mechanism for Nucleotide-dependent Interfacial Cross-talk. J. Mol. Biol. 436, (2024).

30. Mendonça, D. C. et al. An atomic model for the human septin hexamer by cryo-EM. J. Mol. Biol. 433, (2021).

31. Grupp, B. et al. The structure of a tetrameric septin complex reveals a hydrophobic element essential for NC-interface integrity. Commun. Biol. 7, 1–15 (2024).

32. Brausemann, A. et al. Crystal structure of Cdc11, a septin subunit from Saccharomyces cerevisiae. J. Struct. Biol. 193, 157–161 (2016).

33. Farkasovsky, M., Herter, P., Voß, B. & Wittinghofer, A. Nucleotide binding and filament assembly of recombinant yeast septin complexes. Biol. Chem. 386, 643–656 (2005).

34. Valadares, N. F., d’ Muniz Pereira, H., Ulian Araujo, A. P. & Garratt, R. C. Septin structure and filament assembly. Biophys. Rev. 9, 481–500 (2017).

35. Sirajuddin, M., Farkasovsky, M., Zent, E. & Wittinghofer, A. GTP-induced conformational changes in septins and implications for function. Proc. Natl. Acad. Sci. U. S. A. 106, 16592–16597 (2009).

36. Field, C. M. et al. A purified Drosophila septin complex forms filaments and exhibits GTPase activity. J. Cell Biol. 133, 605–616 (1996).

37. Baur, J. D., Rösler, R., Wiese, S., Johnsson, N. & Gronemeyer, T. Dissecting the nucleotide binding properties of the septins from S. cerevisiae. Cytoskeleton 76, 45–54 (2019).

38. Dixon, A. S. et al. NanoLuc Complementation Reporter Optimized for Accurate Measurement of Protein Interactions in Cells. ACS Chem. Biol. 11, 400–408 (2016).

39. Weems, A. & McMurray, M. The step-wise pathway of septin hetero-octamer assembly in budding yeast. Elife 6, (2017).

40. Johnson, C. R., Weems, A. D., Brewer, J. M., Thorner, J. & McMurray, M. A. Cytosolic chaperones mediate quality control of higher-order septin assembly in budding yeast. Mol. Biol. Cell 26, 1323–1344 (2015).

41. Sellin, M. E., Sandblad, L., Stenmark, S. & Gullberg, M. Deciphering the rules governing assembly order of mammalian septin complexes. Mol. Biol. Cell 22, 3152–3164 (2011).

42. Hassell, D. et al. Chaperone requirements for de novo folding of Saccharomyces cerevisiae septins. Mol. Biol. Cell 33, (2022).

43. McMurray, M. A. et al. Septin Filament Formation Is Essential in Budding Yeast. Dev. Cell 20, 540–549 (2011).

44. DeRose, B. T. et al. Production and analysis of a mammalian septin hetero-octamer complex. Cytoskeleton 77, 485–499 (2020).

45. Weirich, C. S., Erzberger, J. P. & Barral, Y. The septin family of GTPases: Architecture and dynamics. Nat. Rev. Mol. Cell Biol. 9, 478–489 (2008).

46. Fischer, M., Frank, D., Rösler, R., Johnsson, N. & Gronemeyer, T. Biochemical Characterization of a Human Septin Octamer. Front. Cell Dev. Biol. 10, (2022).

47. Marques da Silva, R. et al. A key piece of the puzzle: The central tetramer of the Saccharomyces cerevisiae septin protofilament and its implications for self-assembly. J. Struct. Biol. 215, (2023).

48. Hamilton, G. E., Wadkovsky, K. N. & Gladfelter, A. S. A single septin from a polyextremotolerant yeast recapitulates many canonical functions of septin heterooligomers. Mol. Biol. Cell 35, (2024).

49. Saint-Marc, C. et al. Phenotypic consequences of purine nucleotide imbalance in Saccharomyces cerevisiae. Genetics 183, 529–538 (2009).

50. Rudoni, S., Colombo, S., Coccetti, P. & Martegani, E. Role of guanine nucleotides in the regulation of the Ras/cAMP pathway in Saccharomyces cerevisiae. Biochimica et Biophysica Acta (BBA) - Molecular Cell Research 1538, 181–189 (2001).

51. Kim, M. S., Froese, C. D., Xie, H. & Trimble, W. S. Uncovering principles that control septin-septin interactions. Journal of Biological Chemistry 287, 30406–30413 (2012).

52. Sikorski, R. S. & Hieter, P. A system of shuttle vectors and yeast host strains designed for efficient manipulation of DNA in Saccharomyces cerevisiae. Genetics 122, 19–27 (1989).

53. Grinhagens, S. et al. A time-resolved interaction analysis of Bem1 reconstructs the flow of Cdc42 during polar growth. Life Sci. Alliance 3, (2020).

54. Stovicek, V., Borja, G. M., Forster, J. & Borodina, I. EasyClone 2.0: expanded toolkit of integrative vectors for stable gene expression in industrial Saccharomyces cerevisiae strains. J. Ind. Microbiol. Biotechnol. 42, 1519–1531 (2015).

55. Glomb, O. et al. The cell polarity proteins Boi1 and Boi2 direct an actin nucleation complex to sites of exocytosis in Saccharomyces cerevisiae. J. Cell Sci. 133, (2020).

56. Andersen, K. R., Leksa, N. C. & Schwartz, T. U. Optimized E. coli expression strain LOBSTR eliminates common contaminants from His-tag purification. Proteins: Structure, Function and Bioinformatics 81, 1857–1861 (2013).

57. Studier, F. W. Protein production by auto-induction in high density shaking cultures. Protein Expr. Purif. 41, 207–234 (2005).

58. Gasteiger, E. et al. Protein Identification and Analysis Tools on the ExPASy Server. in The Proteomics Protocols Handbook 571–607 (Humana Press, Totowa, NJ, 2005). doi:10.1385/1-59259-890-0:571.

59. da Silveira Tomé, C., Foucher, A., Jault, J. & Housset, D. High concentrations of GTP induce conformational changes in the essential bacterial GTPase EngA and enhance its binding to the ribosome. FEBS J. 285, 160–177 (2018).

60. Armbruster, D. A. & Pry, T. Limit of blank, limit of detection and limit of quantitation. Clin. Biochem. Rev. 29 Suppl 1, S49–S52 (2008).

61. Vogt, A. D. & Di Cera, E. Conformational Selection or Induced Fit? A Critical Appraisal of the Kinetic Mechanism. Biochemistry 51, 5894–5902 (2012).

62. Paul, F. & Weikl, T. R. How to Distinguish Conformational Selection and Induced Fit Based on Chemical Relaxation Rates. PLoS Comput. Biol. 12, e1005067 (2016).

63. Wessel, D. & Flügge, U. I. A method for the quantitative recovery of protein in dilute solution in the presence of detergents and lipids. Anal. Biochem. 138, 141–143 (1984).

64. Hecht, M., Rösler, R., Wiese, S., Johnsson, N. & Gronemeyer, T. An Interaction Network of the Human SEPT9 Established by Quantitative Mass Spectrometry. G3 Genes|Genomes|Genetics 9, 1869–1880 (2019).

65. Cox, J. & Mann, M. MaxQuant enables high peptide identification rates, individualized p.p.b.-range mass accuracies and proteome-wide protein quantification. Nat. Biotechnol. 26, 1367–1372 (2008).

66. Cox, J. et al. Andromeda: A Peptide Search Engine Integrated into the MaxQuant Environment. J. Proteome Res. 10, 1794–1805 (2011).

67. Schindelin, J. et al. Fiji: an open-source platform for biological-image analysis. Nat. Methods 9, 676–682 (2012).

68. Punjani, A., Rubinstein, J. L., Fleet, D. J. & Brubaker, M. A. cryoSPARC: algorithms for rapid unsupervised cryo-EM structure determination. Nat. Methods 14, 290–296 (2017).

69. Punjani, A., Zhang, H. & Fleet, D. J. Non-uniform refinement: adaptive regularization improves single-particle cryo-EM reconstruction. Nat. Methods 17, 1214–1221 (2020).

70. Punjani, A. & Fleet, D. J. 3D variability analysis: Resolving continuous flexibility and discrete heterogeneity from single particle cryo-EM. J. Struct. Biol. 213, 107702 (2021).

71. Emsley, P., Lohkamp, B., Scott, W. G. & Cowtan, K. Features and development of Coot. Acta Crystallogr. D Biol. Crystallogr. 66, 486–501 (2010).

72. Abramson, J. et al. Accurate structure prediction of biomolecular interactions with AlphaFold 3. Nature 630, 493–500 (2024).

73. Liebschner, D. et al. Macromolecular structure determination using X-rays, neutrons and electrons: recent developments in *Phenix*. Acta Crystallogr. D Struct. Biol. 75, 861–877 (2019).

74. Meng, E. C. et al. UCSF ChimeraX: Tools for structure building and analysis. Protein Science 32, (2023).

75. Durell, S. R., Brooks, B. R. & Ben-Naim, A. Solvent-Induced Forces between Two Hydrophilic Groups. J. Phys. Chem. 98, 2198–2202 (1994).

76. Lu, C. et al. OPLS4: Improving force field accuracy on challenging regimes of chemical space. J. Chem. Theory Comput. 17, 4291–4300 (2021).

77. Lee, J. et al. CHARMM-GUI Input Generator for NAMD, GROMACS, AMBER, OpenMM, and CHARMM/OpenMM Simulations Using the CHARMM36 Additive Force Field. J. Chem. Theory Comput. 12, 405–413 (2016).

78. Huang, J. et al. CHARMM36m: an improved force field for folded and intrinsically disordered proteins. Nat. Methods 14, 71–73 (2017).

79. Šali, A. & Blundell, T. L. Comparative Protein Modelling by Satisfaction of Spatial Restraints. J. Mol. Biol. 234, 779–815 (1993).

80. Abraham, M. J. et al. GROMACS: High performance molecular simulations through multi-level parallelism from laptops to supercomputers. SoftwareX 1–2, 19–25 (2015).

81. Jorgensen, W. L., Chandrasekhar, J., Madura, J. D., Impey, R. W. & Klein, M. L. Comparison of simple potential functions for simulating liquid water. J. Chem. Phys. 79, 926–935 (1983).

82. Neria, E., Fischer, S. & Karplus, M. Simulation of activation free energies in molecular systems. J. Chem. Phys. 105, 1902–1921 (1996).

83. Hess, B. P-LINCS: A parallel linear constraint solver for molecular simulation. J. Chem. Theory Comput. 4, 116–122 (2008).

84. Hess, B., Bekker, H., Berendsen, H. J. C. & Fraaije, J. G. E. M. LINCS: A Linear Constraint Solver for molecular simulations. J. Comput. Chem. 18, 1463–1472 (1997).

85. Essmann, U. et al. A smooth particle mesh Ewald method. J. Chem. Phys. 103, 8577–8593 (1995).

86. Darden, T., York, D. & Pedersen, L. Particle mesh Ewald: An N·log(N) method for Ewald sums in large systems. J. Chem. Phys. 98, 10089–10092 (1993).

87. Bussi, G., Donadio, D. & Parrinello, M. Canonical sampling through velocity rescaling. Journal of Chemical Physics 126, (2007).

88. Bernetti, M. & Bussi, G. Pressure control using stochastic cell rescaling. Journal of Chemical Physics 153, (2020).

89. Parrinello, M. & Rahman, A. Polymorphic transitions in single crystals: A new molecular dynamics method. J. Appl. Phys. 52, 7182–7190 (1981).

90. Bahrenberg, T., Jahn, S. M., Feintuch, A., Stoll, S. & Goldfarb, D. The decay of the refocused Hahn echo in double electron–electron resonance (DEER) experiments. Magnetic Resonance 2, 161–173 (2021).

91. Russell, H., Cura, R. & Lovett, J. E. DEER Data Analysis Software: A Comparative Guide. Front. Mol. Biosci. 9, (2022).

92. Jeschke, G. et al. DeerAnalysis 2006—a comprehensive software package for analyzing pulsed ELDOR data. Appl. Magn. Reson. 30, 473–498 (2006).

93. Worswick, S. G., Spencer, J. A., Jeschke, G. & Kuprov, I. Deep neural network processing of DEER data. Sci. Adv. 4, (2018).

94. Jeschke, G. MMM: A toolbox for integrative structure modeling. Protein Science 27, 76–85 (2018).

95. Jeschke, G. Protein ensemble modeling and analysis with MMMx. Protein Science 33, (2024).

96. Schrödinger LLC. The PyMOL Molecular Graphics System, Version 3.1.6.1.

97. Schwede, T., Kopp, J., Guex, N. & Peitsch, M. C. SWISS-MODEL: An automated protein homology-modeling server. Nucleic Acids Res. 31, 3381–3385 (2003).

98. Ashkenazy, H., Erez, E., Martz, E., Pupko, T. & Ben-Tal, N. ConSurf 2010: Calculating evolutionary conservation in sequence and structure of proteins and nucleic acids. Nucleic Acids Res. 38, (2010).

99. Castro, D. K. S. do V. et al. A complete compendium of crystal structures for the human SEPT3 subgroup reveals functional plasticity at a specific septin interface. IUCrJ 7, 462–479 (2020).

